# Novel integrative elements and genomic plasticity in ocean ecosystems

**DOI:** 10.1101/2020.12.28.424599

**Authors:** Thomas Hackl, Raphaël Laurenceau, Markus J. Ankenbrand, Christina Bliem, Zev Cariani, Elaina Thomas, Keven D. Dooley, Aldo A. Arellano, Shane L. Hogle, Paul Berube, Gabriel E. Leventhal, Elaine Luo, John Eppley, Ahmed A. Zayed, John Beaulaurier, Ramunas Stepanauskas, Matthew B. Sullivan, Edward F. DeLong, Steven J. Biller, Sallie W. Chisholm

## Abstract

Horizontal gene transfer accelerates microbial evolution, promoting diversification and adaptation. The globally abundant marine cyanobacterium *Prochlorococcus* has a highly streamlined genome with frequent gene exchange reflected in its extensive pangenome. The source of its genomic variability, however, remains elusive since most cells lack the common mechanisms that enable horizontal gene transfer, including conjugation, transformation, plasmids and prophages. Examining 623 genomes, we reveal a diverse system of mobile genetic elements – cargo-carrying transposons we named tycheposons – that shape *Prochlorococcus*’ genomic plasticity. The excision and integration of tycheposons at seven tRNA genes drive the remodeling of larger genomic islands containing most of *Prochlorococcus*’ flexible genes. Most tycheposons carry genes important for niche differentiation through nutrient acquisition; others appear similar to phage parasites. Tycheposons are highly enriched in extracellular vesicles and phage particles in ocean samples, suggesting efficient routes for their dispersal, transmission and propagation. Supported by evidence for similar elements in other marine microbes, our work underpins the role of vesicle- and virus-mediated transfer of mobile genetic elements in the diversification and adaptation of microbes in dilute aquatic environments – adding a significant piece to the puzzle of what governs microbial evolution in the planet’s largest habitat.

---

*Prochlorococcus* is the smallest and numerically most abundant cyanobacterium in the oceans. It possesses a large pangenome and contains hypervariable genomic islands that have been linked to niche differentiation and phage defense^1–4^. Adaptations to light and temperature broadly define clades in the genus, which are subdivided into a mosaic of coexisting subpopulations in the wild^5–7^. This structure provides stability and resilience to the global population in the face of viral predation and changing environmental conditions^8–10^. While the importance of variable genomic islands in the ecology of *Prochlorococcus* is well-known, how they are formed and acquire new genes remains an open question because most cells lack common means of horizontal gene transfer, such as conjugative systems^11^ and genes for natural competence^12^. Island diversification via gene exchange with cyanophages has been observed for genes involved in photosynthesis, high-light adaptation, and other metabolic functions^13–15^, but evidence for prophage-mediated transduction is rare - some cyanophages carry integrase genes and putative attachment sites have been linked to recombination hotspots in *Prochlorococcus* genomes^16,17^, but only a single partial prophage has been observed in hundreds of available genomes^18^. *Prochlorococcus* cells, moreover, appear devoid of any common mobile genetic elements including plasmids, transposons, insertion sequences, or integrative and conjugative elements^19–21^, with the exception of a few transposons and insertion sequences previously identified in the most basal *Prochlorococcus* clade LLIV^17^. This overall limited ability to utilize canonical horizontal gene transfer mechanisms – seemingly inconsistent with the large and widely distributed pangenome^22,23^ – motivated us to explore genomes of cultured and wild *Prochlorococcus* cells^21^ for evidence of mechanisms promoting genomic island diversity in this group.

## Mobile genetic elements that target genomic islands

We focused our analysis on flexible genes – genes present in only a subset of the population – and on genomic islands – distinct regions in the genome with a high density of these flexible genes. The term “genomic island” is sometimes used in reference to individual mobile genetic elements^24^. We use the term here, however, to refer to larger chromosomal regions with high inter-strain variability, consistent with a widely used definition and earlier descriptions of genomic islands in *Prochlorococcus*^*1*^. These regions in most cases do not represent an individual mobile genetic element, but rather are heterogeneous aggregations of horizontally transferred material, that typically cannot mobilize as an integral unit. We searched for clues as to how such hotspots of variability arise and are maintained over evolutionary timescales.

Horizontal gene transfer promoting genomic island variability is typically associated with two mechanisms: the integration of mobile genetic elements, such as prophages, giving rise to *additive islands*, and the exchange of material through homologous recombination by conserved flanking core genes creating *replacement islands*^*25*^. The two island types have distinct characteristics, as detailed below. We searched for common signatures of both types in 623 *Prochlorococcus* genomes obtained from cultured isolates and single-cells from the wild^21^ (Supplementary Table 1). In all of these genomes, we identified well-defined genomic islands – typically around 8-10 islands per genome between 4 and 200 kbp in size – that comprise about one-quarter of all genes per genome and harbor more than two-thirds of all flexible genes of the *Prochlorococcus* pangenome (Supplementary Fig. 1/2).

We first asked if these islands were replacement islands, i.e. those derived via homologous recombination. Islands of this type are usually found in all lineages of a group and are syntenic, but different strains often carry different gene clusters encoding equivalent biological functions^26^. They are, moreover, exchanged among closely related strains through recombination among their flanking core genes, replacing the entire existing gene cluster with the incoming material^27^. *Prochlorococcus* islands have highly conserved locations and are usually present in all strains (Fig. 1, Supplementary Fig. 3). However, we never observed a complete replacement, only the gain and loss of some genes from one strain to the next (Supplementary Fig. 4). Moreover, recombination rates inferred from island-flanking regions were similar to those of other parts of the genomes (Supplementary Fig. 5), thus, also arguing against whole-island homologous recombination as a key driver of *Prochlorococcus* genomic islands formation.

**Figure 1.**
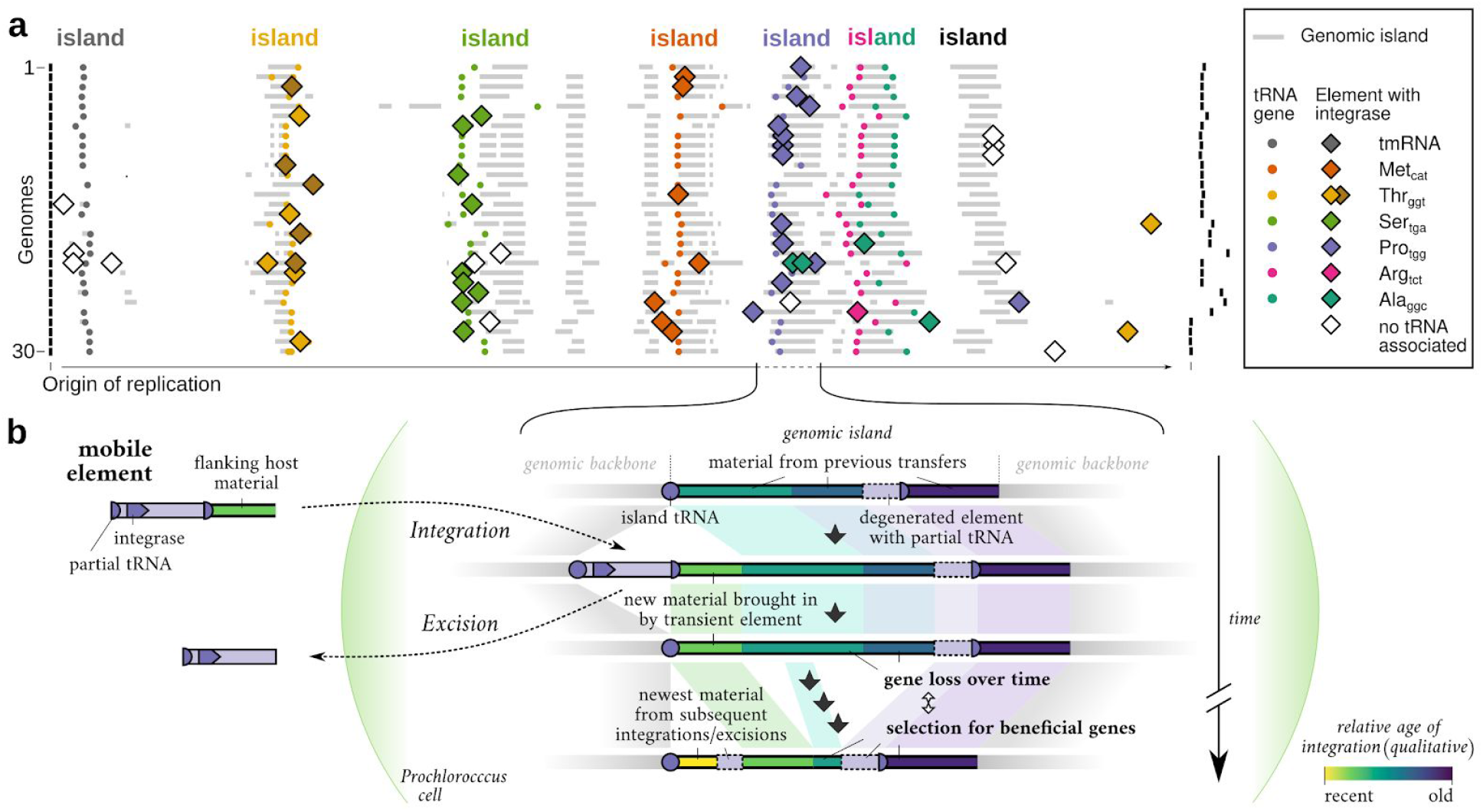
Chromosomal organization of genomic islands and associated mobile genetic elements in *Prochlorococcus*. **a)** 30 selected circular *Prochlorococcus* genomes shown relative to their origin of replication (left to rightmost vertical black bar). Vertical column-like features indicate the predicted genomic islands in conserved locations across the genomes (grey bars). Most islands are associated with one or two specific full-length tRNA genes (colored points, header color). These tRNAs are targeted by mobile genetic elements carrying integrases specific to the different islands (colored diamonds). The genomes shown here are a representative subset of a single *Prochlorococcus* clade (HLII) and are among the most complete genomes in the dataset. The genomes are ordered according to their phylogenetic relationships, i.e. the most closely related genomes are plotted next to each other. **b)** A model for genomic island formation promoted by mobile element activity. Mobile elements integrate and excise at the tRNA gene at the proximal end of the island. Genetic material brought in but not excised later, such as flanking DNA from other hosts, and degenerated elements, accumulates next to the tRNA leading to gene gain and island growth. Gene gain is countered by gene loss and selection, preserving only beneficial acquisitions which may, in turn, become fixed in descending lineages. Due to the directionality of the process, the observed intra-strain heterogeneity is highest at the proximal end of the island right next to the tRNA and decreases towards the distal end of the island.

Next, we searched for signs of mobile genetic element activity – which would be expected for islands of the additive type^25^. Consistent with previous studies^1,28^ most *Prochlorococcus* islands are directly adjacent to 7 tRNA genes (Proline_tgg_, Serine_tga_, Alanin_ggc_, Threonine_ggt_, Arginine_tct_, one of three Methionines_cat_, and the tmRNA gene – a bifunctional transfer-messenger RNA important for releasing stalled ribosomes)(Supplementary Fig. 3). tRNAs are known integration hotspots for a variety of mobile genetic elements^29^. The elements usually encode integrases that recognize a specific attachment site, typically around 40 nucleotides long, in both the excised element and the host genome, and often within tRNA genes^30^. The integrases catalyze site-specific recombination between the two DNA molecules splicing the element into (or out of) the host chromosome. This leaves the integrated elements flanked on both sides with the attachment site motif (one copy from the host, one from the incoming element) – partial tRNAs in this case. Guided by these hallmarks of mobility, we screened *Prochlorococcus* genomes for integrases, other genes linked to a mobile lifestyle, and the presence of full-length and partial tRNAs.

We did not find any of the well-known types of mobile genetic elements except a few previously described insertion sequences and transposons in the most basal *Prochlorococcus* clade (LLIV)^17^. Our search, however, yielded 937 putative integrase-carrying elements – most of which belong to a new family of cargo-carrying DNA transposons that we describe in detail in the next section. These elements specifically integrate at the 7 island-associated tRNAs mentioned above, and in the following, we explore their impact on *Prochlorococcus* genomic islands and niche-differentiation.

The vast majority of all the elements we found were located within genomic islands and in most cases adjacent to the island-associated tRNA at the proximal end of the island (Fig. 1a, Supplementary Fig. 4). In a few cases, we even found multiple elements located right next to each other, separated only by a partial copy of the island-tRNA (Supplementary Doc. 1, examples: AG−363−G23_00382/00397, AG−355−N18_01633/01647, AG−388−A01_00476/00484). These cases likely represent subsequent integrations at the same integration site, with newly incoming elements pushing existing ones further into the island. We also identified remnants of degenerated elements, such as fragmented integrases and more partial tRNAs scattered throughout the islands, suggestive of additional, older integration events.

Compared to the about 8-10 islands that were present among all *Prochlorococcus* genomes in their conserved locations – usually next to a tRNA gene – the elements were much rarer and more transient: on average we observed only 1-2 elements per genome, meaning that most individual islands did not contain any elements. Moreover, 62% of the elements were unique, i.e. only present in a single genome. About 26% were present in two to five – and in rare cases up to 25 – closely related strains likely representing a single integration event in a common ancestor and subsequent vertical inheritance; 12% were present in at least two distantly related genomes – indicating independent integration events. This high diversity and patchy distribution suggest a dynamic system with a large pool of elements that transiently “visit” the islands, thereby, greatly contributing to intrapopulation heterogeneity (Supplementary Fig. 6; elements were defined as the “same” if they shared at least 90% identity over 50% of the shorter element; we did not factor in location because independent integrations will occur at the same tRNA due to the site-specificity of the integrase).

Overall, we found that these putatively active elements only made up 3.6% of the total island material. Degenerated fragments, which lack clear boundaries and are harder to delineate from surrounding material, appear to account for a similar fraction, based on hallmark genes present in non-element parts of islands. Thus, the major fraction of the islands appear not to be made up of the elements, but rather are composed of additional transferred material. Given the location of this material – inside islands abutting tRNAs – we expect that this material was also acquired through site-specific recombination, most likely carried out by transient elements. We hypothesize that the material is brought in as flanking material temporarily captured onto an excising element by imprecise excision from a donor genome, comparable to specialized transduction in lysogenic phages^17,31^ (Fig. 1b). Subsequent precise excision of the element leaves this extra material behind in its new host. In support of this hypothesis, we observed full-length elements in viral metagenomes that carried sequences that appear to be adjacent host material (Supplementary Fig. 7).

As a consequence of this bi-partition, *Prochlorococcus* islands comprise two gene pools with different transfer and turn-over rates: Element genes – which include hallmark genes for integration and replication as well as additional cargo genes – and islands genes that are not part of a functional element. Element genes are highly mobile and can be gained and lost dynamically to quickly match selective pressures. Island genes still likely have higher rates of transfer than core genes but rely on more stochastic processes such as co-transfer of element-flanking material or recombination with active elements. The two pools, moreover, are linked by active gene flow as indicated by a strong overlap in enriched functions between elements and adjacent island regions (Supplementary Fig. 8).

Thus, the majority of material in *Prochlorococcus* islands is not derived from elements directly, but rather appears to be brought in by them as flanking material, suggesting that this type of horizontal transfer is crucial to the island formation process. While genomic data is not direct evidence for such a model of element-driven island formation, all of the described element-associated processes – degeneration, transduction and recombination – fit the observed features including the accretion of horizontally acquired material at the distal end of the islands (the end furthest from the island tRNA). In this model, the tRNAs function as seeds and the transient elements as vectors, gradually contributing to the accumulation of transferred material and, thus, the formation of large, persistent islands.

## New elements for defense and adaptation

To gain a better understanding of the specific roles these novel elements carry out in their hosts we characterized them in two ways: by their hallmark genes related to their function as independent genetic units (recombination and DNA-replication; Supplementary Table 4) and by their cargo genes that appear to convey their broader ecological functions. Below we provide evidence that about half these elements (501 of 937) belong to a single, cohesive and new family of DNA transposons that are likely to confer a fitness advantage on their host cell in particular environments and niches. We named these elements tycheposons – referring to the Greek deity Tyche, a daughter of Oceanus and a guardian of fortune and prosperity. The other half of the elements were more cryptic, lacking identifiable hallmark genes other than integrases and excisionases, and we excluded them from analyses beyond those two genes.

While the diversity of prokaryotic mobile genetic elements, such as phages, plasmids and numerous types of transposons, is immense, tycheposons differ from all of the known classes and families in significant ways, including a key feature in transposable elements classification: the enzyme that catalyzes the elements’ movement^32^. Arguably the most abundant prokaryotic transposable elements are insertion sequence-like transposons. Their autonomous representatives – insertion sequences (IS) and composite transposons – encode transposases for their integration and excision. These enzymes usually exhibit little to no site-selectivity enabling the encoding elements to transpose, i.e. integrate at different locations within the same genome. Moreover, these classic transposons lack genes for independent replication. By contrast, tycheposons and the cryptic elements we identified, encode site-specific integrases (910 out of 937 contain a phage-integrase-like tyrosine recombinase and the rest a large serine recombinase) and many of them carry genes associated with autonomous replication (polymerases, primases, helicases) (Fig. 2, Supplementary Table 5).

**Figure 2.**
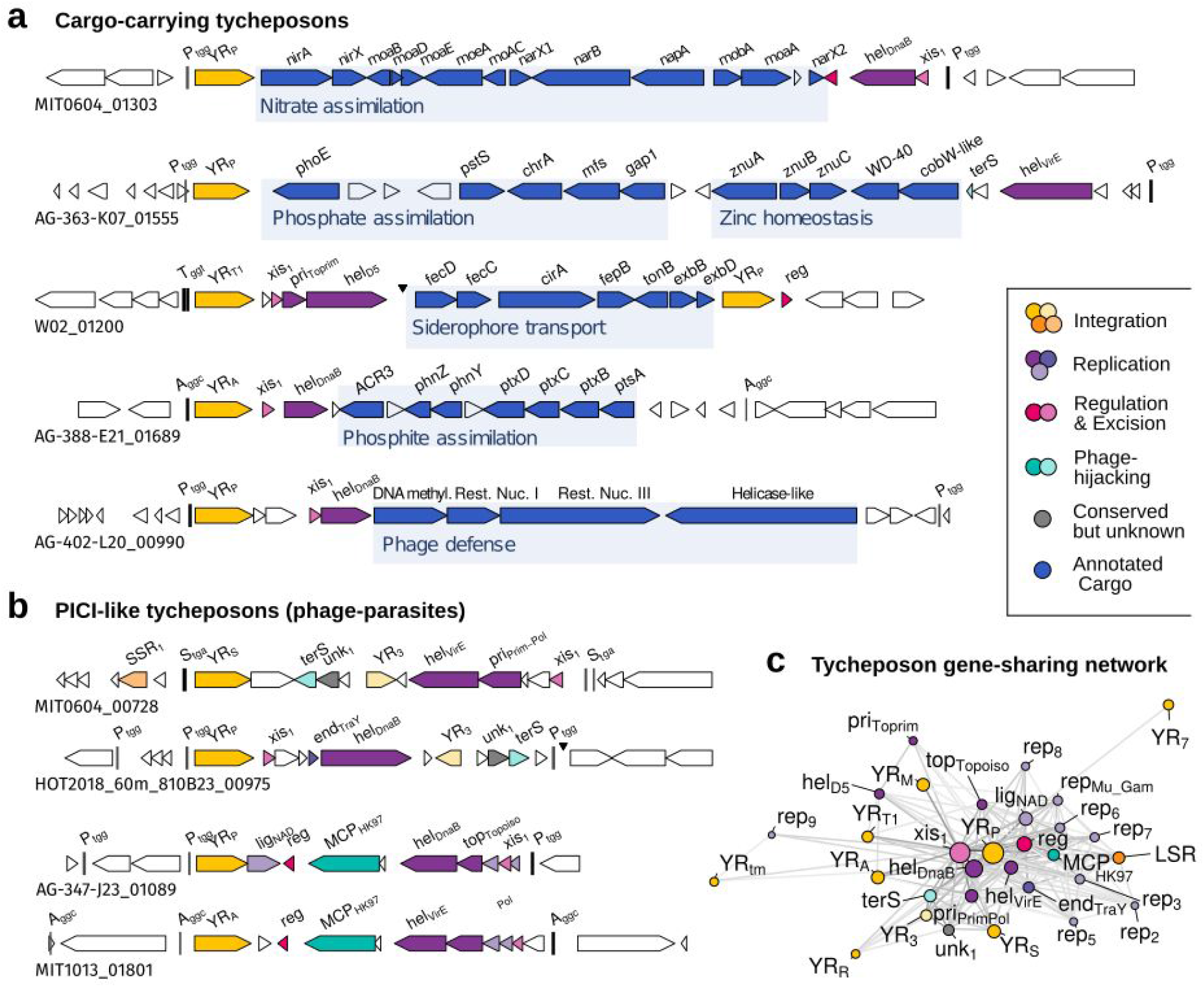
Structure and function of tycheposons in *Prochlorococcus*. Examples of two types of tycheposons illustrating their modular structure: **a)** Cargo-carrying tycheposons selected here because of their clear ecological relevance in ocean ecosystems, and **b)** PICI-like tycheposons, carrying either a terS or MCP viral packaging gene likely used to hijack phage capsids for dispersal. Cargo modules were annotated with roles in nitrate assimilation^36^, siderophore transport^18^, phosphate assimilation^37^, zinc homeostasis^38^ and phosphite assimilation^39^. **c)** A gene sharing network of tycheposon genes indicating that the functionally different tycheposon types (PICI-like and cargo-carrying) are evolutionarily linked through their integration and replication modules (all genes form a single cluster). Gene labels: full-length and partial tRNA genes are labeled with single-letter amino acid code and their anticodon, e.g. S_tga_ = tRNA-Serin_TGA_. YR = tyrosine recombinase; MCP = major capsid protein; terS = terminase small subunit; xis = excisionase; hel = helicase; top = topoisomerase; lig = ligase; reg = transcriptional regulator; pri = primase or primase/polymerase or primase/helicase; end = endonuclease; SSR = small serine recombinase; unk = conserved unknown. Some annotations are further labeled with more specific subprofiles indicating specific families of helicases, for example (hel_DnaB_ or hel_VirE_). Node size corresponds to the number of genes matching individual profiles. More detailed information on the gene profiles is available in Supplementary Table 4.

Site-specific integrases and autonomous replication are not at all uncommon in mobile genetic elements, however, they are typically associated with either eukaryotic elements^33^ or large prokaryotic elements with distinctive life-styles: lysogenic phages, which encode their own capsids, plasmids, which typically exists as extrachromosomal DNA molecules, and integrative conjugative elements (ICE), which encode conjugation machinery that enable them to move between cells^34,35^. Tycheposons, however, lack these complex functional traits.

Based on their hallmark functions and overall genomic organization, tycheposons appear most similar to phage-inducible chromosomal islands (PICIs)^40–42^. PICIs are mobile genetic elements similar in size to tycheposons (10-20 kbp), that also make use of site-specific integrases and carry genes associated with autonomous replication. The key characteristic of PICIs is their life-stye: they are parasites of phages - reactivating from the host genome during phage infection, replicating, interfering with the replication of the phage, hijacking newly produced phage capsids so they can spread to other cells. PICIs encode regulatory genes to sense phage infections, genes to interfere with phage replication, and either a predicted major capsid protein, a building block of a phage capsid, or a small terminase subunit, a part of the protein complex that pumps phage DNA into newly formed phage capsids. As such, PICIs increase host population resistance to infections by diverting resources away from the infecting phage.

In comparing PICIs and tycheposons, we examined both functional traits and evolutionary history. We found that all tycheposons are part of one evolutionarily linked group sharing a set of hallmark genes not shared with other groups of mobile genetic elements, including PICIs, ICEs, or phages (Fig. 2c). In terms of their broader ecological roles, however, tycheposons fall into two categories: shuttles for dedicated cargo such as operons for the acquisition of growth-limiting resources and putative PICI-like phage parasites (Fig. 2a/b, Supplementary Table 5). Interestingly, this functional specialization is reflected neither in different hallmark gene sets nor in the phylogeny of the integrases, suggesting that both types evolve as one cohesive group linked by active gene flow.

Analogous to previously described PICIs, the PICI-like tycheposons in our system also carry either a predicted major capsid protein or a small terminase subunit, and we anticipate a life-style similar to that of known PICIs, ultimately providing population-level resistance against certain phage infections. Interestingly, however, we never observed PICI-like tycheposons with additional cargo genes, while this appears to be common in known PICIs^42^. PICI-like tycheposons also account for only about 15% of the elements in our dataset (81 out of 501).

The majority of tycheposons in our dataset (419 out of 501) lack genes for viral packaging or interference and their cargo appears to reflect selection pressures of the local environment (Fig. 2, Supplementary Table 5). They often carry only the two small gene modules for integration (∼1 kbp) and replication (∼2-4 kbp) in addition to up to 10 kbp or more of dedicated cargo. In many cases, the cargo of this set of tycheposons is comprised of genes of unknown functions – a standing challenge of annotation for wild microbes^43^. The genes we could annotate, however, revealed a broad spectrum of functions associated with ecologically relevant processes such as nutrient uptake. Remarkably, those functions include the three key nutrients that limit primary productivity in the global oceans: nitrogen, phosphorus and iron (Fig. 2a). These complex metabolic traits – some of them typically encoded in the core genome^36,44^ – are captured in tycheposons, turning them into mobile units able to boost host fitness in environments where a particular nutrient is a strong selective agent. These findings underpin the role of tycheposons in accelerating microbial adaptation and genome evolution, especially with respect to key metabolic traits, which are typically thought to be more rarely gained and lost. The elements’ mobility is likely also an important contributor to the ocean-wide stability of the *Prochlorococcus* collective^7^.

A phylogenetic comparison of the tyrosine integrases encoded by tycheposons with those of other mobile genetic elements, viruses, bacteria and archaea^45^ revealed that, with few exceptions, they form a monophyletic clade; they are more closely related to each other than to integrases from any other system (Fig. 3). This is unexpected because integrases often jump between viruses, elements, and hosts, resulting in more convoluted evolutionary histories^26^. At the same time, these new integrases account for more than 10% of the phylogenetic diversity observed among known integrases across various domains of life, further supporting that they constitute an evolutionarily ancient and diverse group, and suggesting that the elements carrying them evolved independently over similar timescales.

**Figure 3.**
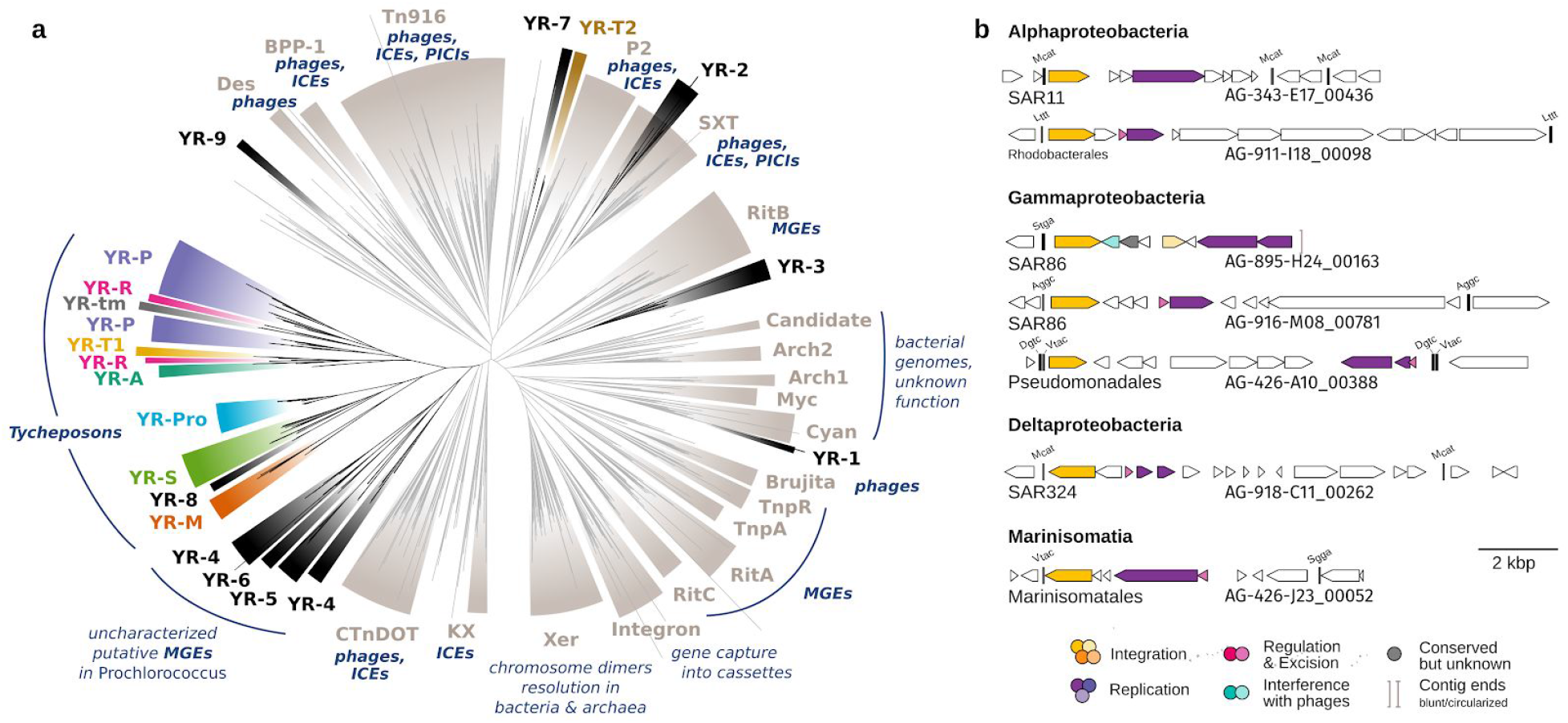
Phylogenetic placement of tycheposon tyrosine recombinases and presence of putative tycheposons in other marine bacteria. **a**) Maximum likelihood phylogeny of a comprehensive, representative set of Xer-like tyrosine recombinases from bacterial, archaeal and phage genomes as well as those identified in *Prochlorococcus* genomes in this study. Common types (grey) are labeled according to the classification introduced by Smyshlyaev et al.^45^. Integrases found in phages, ICEs (integrative conjugative elements) or other mobile genetic elements (MGEs) and integrases associated with other functions are indicated by navy blue labels. Integrases from known PICIs (phage-inducible chromosomal islands)^42,46,47^ match two known types: Tn916 and SXT. Most *Prochlorococcus* tRNA-associated (colored) and non-tRNA-associated (black) tyrosine recombinases form a monophyletic group, together with a *Prochlorococcus* core gene integrase (light blue), and are distinct from the previously described common types. 474 of the 501 tycheposons described here carry one of the tRNA-associated tyrosine recombinases. **b**) Examples of PICI-like and cargo-carrying tycheposons in a variety of other marine bacterial groups. Gene labels: tRNA genes and snippets are labeled with single-letter amino acid code and their anticodon, e.g. S_tga_ = tRNA-Serin_TGA_.

Within the tycheposons, the integrases further cluster by the different tRNA genes they target (Fig. 3) such that each island has its specific, proximal tRNA, and each tRNA its specific integrase making integrases island-specific. Thus, the integrases determine into which island a tycheposon will integrate and partition the pool of tycheposons into subpopulations that affect different islands. This mechanism has the potential to control which tycheposons and flanking genomic regions are more likely to recombine. It has previously been shown that different genes of similar ecological functions appear to occur in the same islands across different strains^1,3^. The island specificity of tycheposons could be one important factor for promoting this differentiation of islands with respect to broader ecological themes.

Based on this evidence we define tycheposons as an independent lineage of mobile genetic elements with the following features: a site-specific integrase – in the majority of cases a site-specific tyrosine recombinase lineage-specific to tycheposons (Fig. 3); a common set of shared hallmark genes facilitating mobility and replication (Fig. 2c, Supplementary Data for gene profiles); a conserved gene organization with an inward-facing integrase at one end, and an optional, small replication module either next to the integrase or at the opposite end (Fig. 2a/b); cargo providing adaptation to the local environment – e. g. nutrient assimilation – or phage interference (Fig. 2a/b).

To determine whether tycheposons were specific to *Prochlorococcus* or might be a feature of genomically streamlined ocean microbes, we examined a large number of other marine bacterial genomes for similar elements by looking for the same hallmarks we have associated with tycheposons. This search revealed related elements in a broad variety of bacterial groups including Alpha-, Gamma- and Deltaproteobacteria (Fig 3b), where they likely play similar roles in promoting genomic plasticity and adaptability.

## Evidence of activity and mobility

To get a more detailed picture of the activity of tycheposons, we turned to *Prochlorococcus* strain MIT0604, which is particularly interesting as it contains 7 tycheposons including two with 100% sequence identity encoding the complete gene cluster for nitrate assimilation^36,44^ (Supplementary Fig. 9). We took advantage of the fact that we had maintained this strain through serial transfer in two separate liquid cultures for 10 years – one propagated on media with NH_4_^+^ as the only N-source, the other on the same media but with NO_3_^-^ as the only N-source. The latter culture was sequenced shortly after separation in 2011, and then both independent cultures were sequenced again in 2019. Comparison of the genomes of all three revealed significant genomic rearrangements, all of which centered around the tycheposons: we observed gain, loss and duplication of tycheposons as well as an associated chromosomal inversion between two identical copies of the nitrate assimilation cluster they carry (Fig. 4, Supplementary Fig. 10). Because the tycheposon attachment sites reside within genomic islands, the modifications were confined to islands, while the rest of the genome remained unaltered. Importantly, the rearrangements also point to the selective advantage provided by tycheposons carrying cargo with adaptive metabolic functions: The most conspicuous changes were linked to the tycheposon containing the nitrate assimilation gene cluster when cultures were maintained with NO_3_^-^ as the sole N source. That is, the cargo metabolic function, nitrate assimilation has become an independent ‘plug-in’ cassette that can be duplicated under the selective pressure (NO_3_^-^, in this case), lost when useless (if NH_4_^+^ is the only N-source), and, in the wild, likely also more flexibly transferred between cells than would be a core genome trait.

**Figure 4.**
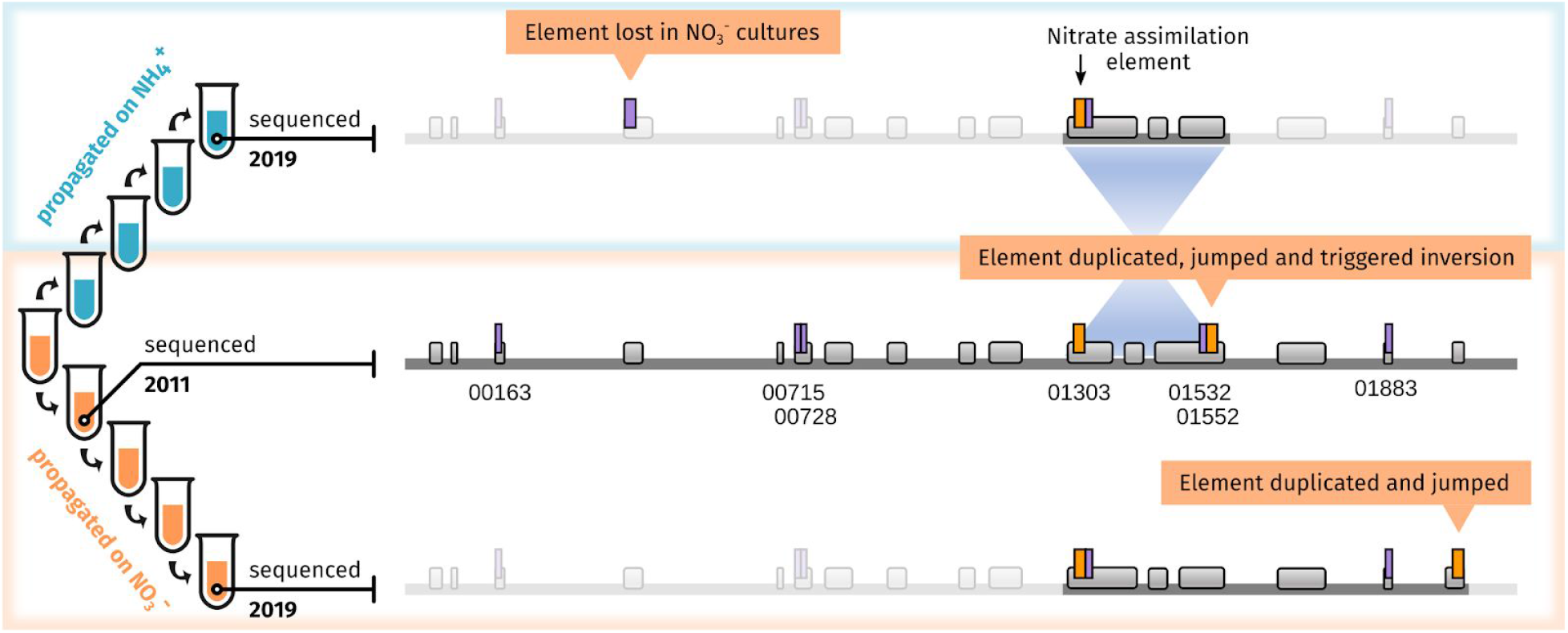
*Prochlorococcus* MIT0604 genome and genomic islands remodeling induced by its integrated tycheposons. Comparison of 3 *Prochlorococcus* MIT0604 genomes from two liquid cultures maintained through serial transfers for 10 years: MIT0604 (NO3 ^-^) was kept growing on nitrate as the sole nitrogen source while MIT0604 (NH4 ^+^) was transferred onto and maintained on ammonia as the sole nitrogen source. The sketch on the left gives a simplified representation of the sequence of propagation of the two lineages, and when the genomes were sequenced. The reference MIT0604 genome sequenced in 2011 carries seven integrated tycheposons including two identical copies of a tycheposon containing the nitrate assimilation gene cluster (Fig. 2) in two different genomic islands. The dark grey line represents the reference genome (1.78 Mbp) and regions that changed in the other lineages relative to the reference; faded regions remained unchanged. Grey boxes represent genomic islands, purple boxes represent different tycheposons, the orange boxes the tycheposons carrying the nitrate assimilation cluster (see details in Supplementary Fig. 9). The genomes sequenced in 2019, furthermore, showed that identical copies of a tycheposon in a single genome can trigger chromosomal inversions probably through homologous recombination ^48^ thereby, revealing yet another mechanism by which these elements can promote genomic plasticity – in this case even at the chromosomal scale. Comparing the cultures maintained on different N-sources, we observe that the duplication and movement of the nitrate-assimilation tycheposon happened in both lineages maintained on nitrate. Each jump happened in two distinct genomic island locations, inserting next to a short segment of the same tRNA abutting the original copy, supporting our model of site-specific recombination.

We also tested four additional tycheposon-containing *Prochlorococcus* strains for tycheposon mobility using PCR amplification of the circular and excised element intermediates. We found that all cultures are internally heterogeneous, with subpopulations of cells missing a tycheposon that excised, or possessing tandem repeat junctions (Supplementary Fig. 11), adding to our understanding of how hundreds of coexisting subpopulations varying by small sets of genes might arise in the environment^5–7^.

To better understand what conditions would trigger tycheposon mobility, we measured the transcriptional response of an integrase associated with a cargo-carrying tycheposon in MIT0604 to a wide range of stressors (Supplementary Fig. 12) – some of which a cell might experience in the environment, and others artificial. The only significant relative increase in transcripts occurred in response to mitomycin C, a DNA alkylating agent commonly used to trigger DNA damage. Mitomycin C, well known to induce prophage excision, also induced other integrases in other strains (Supplementary Fig. 13). This treatment triggered detectable mobilization of some tycheposons (Supplementary Fig. 11), showing that the effect is not specific to the MIT0604 strain.

Further, a genome-wide transcriptome analysis of MIT0604 (Fig. 5) revealed that most integrases, putative excisionases and replication genes all had elevated transcripts when subjected to mitomycin C, indicating a universal regulatory mechanism. We note that other DNA damaging treatments (Supplementary Fig. 12) including UV shock – a known source of DNA damage in the surface ocean^50–53^ – did not significantly increase relative transcript abundance of the integrase (Fig. 5). This suggests something distinctive about the mitomycin C inhibitory mechanism, which we suspect lies in its ability to cause lethal DNA crosslinks^54,55^ that lead to replication fork arrests. Thus we postulate that this tight regulatory mechanism would ensure that mobility genes remain mostly silent, avoiding the toxicity generally linked to their activity^34,56^, and are only induced in a fraction of the population^57^, even in times of stress such as nutrient starvation or high UV exposure.

**Figure 5.**
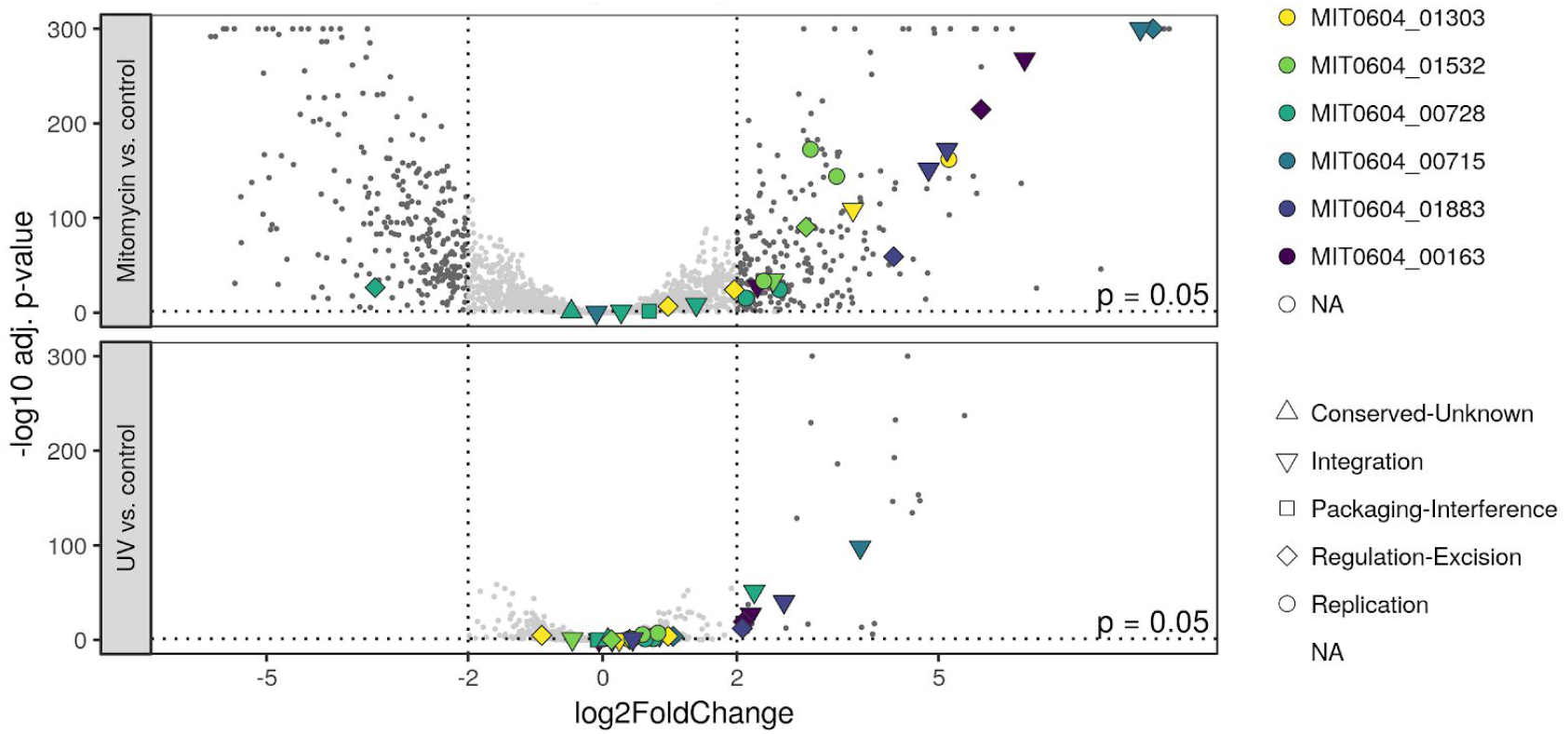
Differential gene expression of tycheposon hallmark genes under DNA damage stress. Volcano plot showing the log_2_ fold change of gene expression of cultures treated with either mitomycin C (top) or a UV shock (bottom) relative to the control treatment. The y-axis shows the false discovery rate corrected p-value. Coloured points show hallmark genes with the shape indicating their function. Overall, the mitomycin C treatment produced a much stronger signal (a phenomenon already observed in other bacteria, 30% of *E. coli* genome is differentially expressed by mitomycin C^49^), indicating that the cells are better equipped to handle UV-induced damage. The majority of tycheposon hallmark genes are strongly upregulated by mitomycin C but not UV shock. In the mitomycin C treatment, several tycheposons show co-upregulation of their hallmark genes (integrase, excisionase, primase/helicase).

## Tycheposons in viral capsids and vesicles

Next, we asked how tycheposons might move from cell to cell in the dilute oceans. First, we looked at viral particles as potential vectors by screening viral-fraction metagenomic libraries from ocean samples where *Prochlorococcus* is abundant. We examined both long nanopore reads^58^ and short-read contigs^59,60^, and indeed, found an abundance of PICI-like tycheposons (Supplementary Fig. 7). Because they carry the same type of phage-packaging genes as previously described PICIs – we also anticipate they exhibit the same life-style: the tycheposons likely reactivate upon phage infection, replicate, and hijack phage capsids with the help of their phage-like packaging genes. In turn, they may reduce, either through competition for limited resources or through specific interference, the number of infective phage particles produced in the infected cell, thereby conveying resistance against a further spread of the infection on the population. At the same time, by hijacking phage particles the PICI-like tycheposons promote their own dissemination leading to their prevalence in marine viral metagenomes. We could not, however, identify cargo-carrying tycheposons in the phage fraction samples, suggesting alternative transfer routes for them and motivating us to look elsewhere.

Because *Prochlorococcus* does not appear to engage in natural conjugation or transformation, we wondered if extracellular vesicles, known to contain DNA and to be released by *Prochlorococcus* and other marine microbes^61^, might serve as vectors. Comparing vesicle- and cellular-fraction metagenomic data from seawater we observed a specific enrichment of almost all predicted tycheposon hallmark genes in vesicles (Fig. 6). Similarly, most segments of tRNAs acting as tycheposon attachment sites and, thereby, as genomic island seeds, were also significantly enriched over other tRNA segments in vesicles. The specific enrichments of both of these features strongly suggest that vesicles serve as a common means of dispersal for these elements, thereby casting new light on the importance of vesicles as vectors for horizontal gene transfer in natural microbial communities.

**Figure 6.**
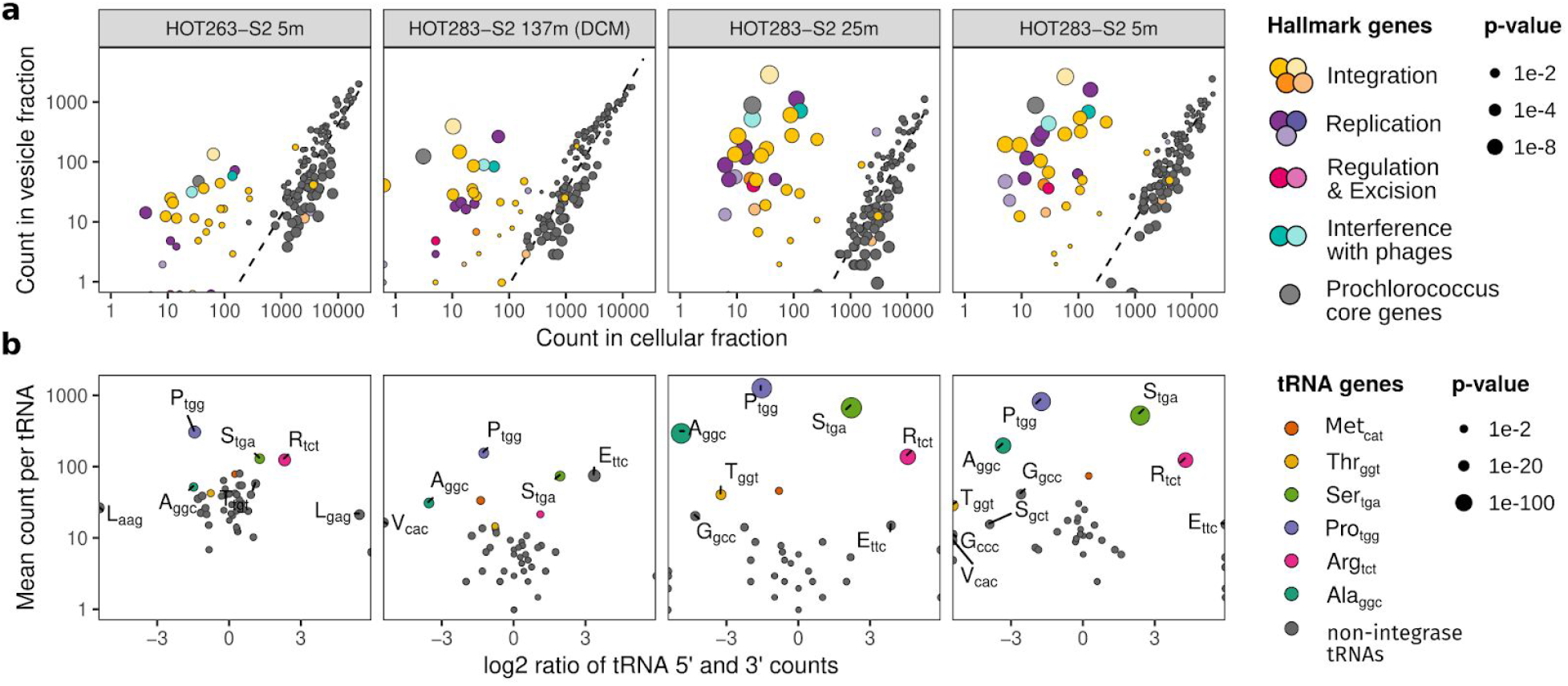
Differential abundance of tycheposon signatures in vesicle-fraction metagenomes from the ocean. **a)** Read counts obtained for the same gene profiles from vesicle- and cellular fraction metagenomes from the North Pacific Subtropical Gyre (dark grey: *Prochlorococcus* core gene profiles, colored: tycheposon hallmark genes, plot labels: Hawai’i Ocean Time-series cruise, station and depth the samples were collected from, DCM: deep chlorophyll maximum). Larger points correspond to a higher degree of deviation (lower p-value in edgeR differential abundance analysis) from the expected abundance ratio (dotted line). **b)** Abundances of tRNA 5’ and 3’ segments in vesicle-fraction metagenomes. Deviation from a 1:1 ratio on the x-axis indicates an over- or underrepresentation of the respective ends of the tRNA sequence in vesicles, suggesting in the most extreme cases an excess of partial, integrase-targeted tRNAs of more than 10-fold (A_gcc_, R_tct_). Higher mean counts for both ends on the y-axis indicate higher overall abundance. Larger points correspond to a higher degree of deviation scaled according to p-values after Bonferroni correction.

## Discussion and Conclusion

Given that *Prochlorococcus*, and likely other oligotrophic bacteria, appear to generally lack the canonical modes of horizontal gene transfer, the mechanism through which their extraordinarily diverse pangenomes have evolved has been elusive. Here, we present evidence – using genomic, experimental, and field data – arguing that a unique set of mobile genetic elements, which we named tycheposons is involved. Tycheposons can carry functional modules important for niche-differentiation among sub-populations of cells along environmental gradients. Strikingly, the modules include genes involved in the uptake of nitrogen, phosphorus and iron – the three key nutrients limiting primary productivity in the global ocean^62^. Rarely does one see such a strong signal highlighting the important agents of selection over a vast ecosystem.

A subset of the tycheposons appear to be phage satellite elements known as PICIs^42,47^ and those reported here are the first of their kind described in cyanobacteria. The different evolutionary histories of tycheposons and PICI integrases suggests that their structural similarity could have arisen from convergent evolution. It is unclear whether PICI-like tycheposons were initially introduced in a *Prochlorococcus* ancestor, and diverged into the larger family of cargo-carrying tycheposons, or if cargo-carrying tycheposons acquired phage interference genes – however, all tycheposons are interconnected by the swapping of the integrases and replication genes. That the two broad functional categories of tycheposons involve both relief from growth limitation and mortality defense is elegant in its ecological simplicity.

The role of tycheposons, however, appears to go beyond just the function of their cargo. They appear to be at the center of a system that is responsible for the formation and remodeling of *Prochlorococcus’* much larger reservoir of genomic variability – its genomic islands. Our analyses of genes present in and near genomic islands suggest that the horizontally acquired regions inside islands that are not part of tycheposons were brought in with them. Moreover, the presence and activity of tycheposons appear to stimulate rearrangements inside islands and beyond, thereby, promoting the genomic diversity underlying the population structure of *Prochlorococcus* cells in the wild, consisting of hundreds to thousands of coexisting subpopulations varying by small cassettes of genes within those islands^5,6^.

Our metagenomic analysis further shows that tycheposons are contained within both phage capsids and membrane vesicles, implicating these structures in the movement of tycheposons among cells in the marine environment. As structured particles that can diffuse through aqueous environments, vesicles and phage are both well-suited to transporting genetic information and delivering it into other ocean microbes. Extracellular vesicle-mediated horizontal gene transfer - recently proposed to be referred to as vesiduction^63^ - occurs in diverse microbial systems^64–70^ and may mediate exchanges between more distantly related taxa than do viruses^71^, which frequently exhibit narrow host ranges^72^. Given the abundance of vesicles in the oceans^73^ and the enrichment of tycheposons within natural vesicle samples, we propose that vesicle-mediated exchange of these elements could be a prevalent mechanism for horizontal gene exchange among bacteria in the marine ecosystem. They are in effect, a family of minimalistic gene shuttles. Though much remains to be learned concerning how such elements are packaged within vesicles, as well as the factors influencing their delivery and integration into other cells, our findings highlight the potential contribution of extracellular vesicles in shaping the genetic structure of marine microbial populations. They have left their signatures all over the genomes of marine microbes implicating themselves as a potentially important mechanism for diversification and adaptation in dilute environments, and possibly beyond.

## Material and Methods

See supplementary information.

## Data availability

Raw Prochlorococcus isolate and single-cell assemblies are available through public databases such as NCBI Genbank, the reference-scaffolded assemblies with our lifted annotations are available from GitHub (https://github.com/thackl/pro-tycheposons) and Zendodo^74^. Global Ocean Reference Genomes can be accessed at https://osf.io/pcwj9 and as EBI/NCBI bioproject PRJEB33281. The Tara Oceans viral-fraction short-read metagenome contigs can be accessed through the Data Commons portal of iVirus under GOV2.0 (filename: Tara_assemblies.tar.gz). Station ALOHA viral-fraction nanopore reads are available from the NCBI Sequence Read Archive (SRX7079550). Vesicle-fraction metagenomes are available from the NCBI Sequence Read Archive (SRP272691).

## Code availability

Custom code used to detect and annotate genomic islands and tycheposons is available via GitHub (https://github.com/thackl/pro-tycheposons) and Zenodo^74^.

## Acknowledgments

This study was supported in part by the Simons Foundation (Life Sciences Project Award IDs 337262, 509034SCFY17, 647135 to SWC, SCOPE Award ID 329108 to EFD and SWC, SCOPE ALOHA award ID 721223 to EFD), Gordon and Betty Moore Foundation Award IDs 3777 to EFD and 3790 to MBS, the Department of Energy (248445 to MBS), and the National Science Foundation (DBI 0424599, OCE-1153588, OCE-1356460 and IOS-1645061 to SWC and OCE-1829831 to MBS). We thank Wei Ding for support with the panX pangenome software, Gera Smyshlyaev for access to the HMM used to classify known tyrosine-recombinases, Kathryn Kauffman for feedback on the analysis of PICI-like tycheposons, and Kristina Haslinger, Daniel S. Fisher and Jed Fuhrman for comments on the manuscript. We also thank Jamie Becker, Jessie Berta-Thompson, and Elena Kazamia for assistance with vesicle metagenome sampling.

## Author information

### Contributions

TH, RL and MJA conceived the study. TH led the bioinformatics analyses. TH, MJA and ET carried out the computational analyses with contributions from SLH, PB and GEL. RL led the experimental work on tycheposon mobility and activity. RL, CB, ZC carried out those experiments. SJB led the vesicle-related experiments and carried them out together with KDD and AA. EL, JB, JME, and EFD generated and provided the viral-fraction nanopore data and Station ALOHA time-series metagenomic contigs. AAZ and MBS generated and provided the viral-fraction metagenomic contigs from Tara Oceans. RS provided the Global Ocean Reference Genomes database. TH, RL, MJA and SWC wrote the manuscript with contributions from all coauthors. SWC supervised the project.

## Additional information

Supplementary Information is available for this paper. Correspondence and requests for materials should be addressed to TH and SWC.

## Supplementary information

Supplementary Figures, Supplementary Tables, Supplementary Data - including scripts, reference genomes, gene profiles and annotations of all identified mobile genetic elements in *Prochlorococcus*, other bacteria and viral metagenomes.

## References

1. Coleman, M. L. et al. Genomic islands and the ecology and evolution of Prochlorococcus. Science 311, 1768–1770 (2006).

2. Rodriguez-Valera, F. et al. Explaining microbial population genomics through phage predation. Nat. Rev. Microbiol. 7, 828–836 (2009).

3. Avrani, S., Wurtzel, O., Sharon, I., Sorek, R. & Lindell, D. Genomic island variability facilitates Prochlorococcus-virus coexistence. Nature 474, 604–608 (2011).

4. Delmont, T. O. & Eren, A. M. Linking pangenomes and metagenomes: the Prochlorococcus metapangenome. PeerJ 6, e4320 (2018).

5. Kashtan, N. et al. Single-cell genomics reveals hundreds of coexisting subpopulations in wild Prochlorococcus. Science 344, 416–420 (2014).

6. Kashtan, N. et al. Fundamental differences in diversity and genomic population structure between Atlantic and Pacific Prochlorococcus. ISME J. 11, 1997–2011 (2017).

7. Biller, S. J., Berube, P. M., Lindell, D. & Chisholm, S. W. Prochlorococcus: the structure and function of collective diversity. Nat. Rev. Microbiol. 13, 13–27 (2015).

8. Ahlgren, N. A., Perelman, J. N., Yeh, Y. & Fuhrman, J. A. Multi-year dynamics of fine-scale marine cyanobacterial populations are more strongly explained by phage interactions than abiotic, bottom-up factors. Environ. Microbiol. 21, 2948–2963 (2019).

9. Ribalet, F. et al. Light-driven synchrony ofProchlorococcusgrowth and mortality in the subtropical Pacific gyre. Proceedings of the National Academy of Sciences vol. 112 8008–8012 (2015).

10. García-García, N., Tamames, J., Linz, A. M., Pedrós-Alió, C. & Puente-Sánchez, F. Microdiversity ensures the maintenance of functional microbial communities under changing environmental conditions. ISME J. 101, 1–15 (2019).

11. Guglielmini, J., de la Cruz, F. & Rocha, E. P. C. Evolution of conjugation and type IV secretion systems. Mol. Biol. Evol. 30, 315–331 (2013).

12. Johnston, C., Martin, B., Fichant, G., Polard, P. & Claverys, J.-P. Bacterial transformation: distribution, shared mechanisms and divergent control. Nature Publishing Group 12, 1–16 (2014).

13. Lindell, D. et al. Transfer of photosynthesis genes to and from Prochlorococcus viruses. Proc. Natl. Acad. Sci. U. S. A. 101, 11013–11018 (2004).

14. Zeidner, G. et al. Potential photosynthesis gene recombination between Prochlorococcus and Synechococcus via viral intermediates. Environ. Microbiol. 7, 1505–1513 (2005).

15. Sullivan, M. B. et al. Prevalence and evolution of core photosystem II genes in marine cyanobacterial viruses and their hosts. PLoS Biol. 4, e234 (2006).

16. Sullivan, M. B., Coleman, M. L., Weigele, P., Rohwer, F. & Chisholm, S. W. Three Prochlorococcus Cyanophage Genomes: Signature Features and Ecological Interpretations. PLoS Biology 3, e144 (2005).

17. Sullivan, M. B. et al. The genome and structural proteome of an ocean siphovirus: a new window into the cyanobacterial ‘mobilome’. Environ. Microbiol. 11, 2935–2951 (2009).

18. Malmstrom, R. R. et al. Ecology of uncultured Prochlorococcus clades revealed through single-cell genomics and biogeographic analysis. ISME J. 7, 184–198 (2013).

19. Rocap, G. et al. Genome divergence in two Prochlorococcus ecotypes reflects oceanic niche differentiation. Nature 424, 1042–1047 (2003).

20. Biller, S. J. et al. Genomes of diverse isolates of the marine cyanobacterium Prochlorococcus. Scientific Data 1, 140034 (2014).

21. Berube, P. M. et al. Single cell genomes of Prochlorococcus, Synechococcus, and sympatric microbes from diverse marine environments. Scientific Data 5, 180154 (2018).

22. Bentkowski, P., Van Oosterhout, C. & Mock, T. A Model of Genome Size Evolution for Prokaryotes in Stable and Fluctuating Environments. Genome Biol. Evol. 7, 2344–2351 (2015).

23. Karcagi, I. et al. Indispensability of Horizontally Transferred Genes and Its Impact on Bacterial Genome Streamlining. Mol. Biol. Evol. 33, 1257–1269 (2016).

24. Boyd, E. F., Almagro-Moreno, S. & Parent, M. A. Genomic islands are dynamic, ancient integrative elements in bacterial evolution. Trends Microbiol. 17, 47–53 (2009).

25. López-Pérez, M., Gonzaga, A. & Rodriguez-Valera, F. Genomic Diversity of ‘Deep Ecotype’ Alteromonas macleodii Isolates: Evidence for Pan-Mediterranean Clonal Frames. Genome Biol. Evol. 5, 1220–1232 (2013).

26. Rodriguez-Valera, F., Martin-Cuadrado, A.-B. & López-Pérez, M. Flexible genomic islands as drivers of genome evolution. Curr. Opin. Microbiol. 31, 154–160 (2016).

27. López-Pérez, M., Martin-Cuadrado, A.-B. & Rodriguez-Valera, F. Homologous recombination is involved in the diversity of replacement flexible genomic islands in aquatic prokaryotes. Front. Genet. 5, 147 (2014).

28. Liu, H.-L. & Zhu, J. Analysis of the 3’ ends of tRNA as the cause of insertion sites of foreign DNA in Prochlorococcus. J. Zhejiang Univ. Sci. B 11, 708–718 (2010).

29. Williams, K. P. Integration sites for genetic elements in prokaryotic tRNA and tmRNA genes: sublocation preference of integrase subfamilies. Nucleic Acids Res. 30, 866–875 (2002).

30. Grindley, N. D. F., Whiteson, K. L. & Rice, P. A. Mechanisms of site-specific recombination. Annu. Rev. Biochem. 75, 567–605 (2006).

31. Sato, K. & Campbell, A. Specialized transduction of galactose by lambda phage from a deletion lysogen. Virology 41, 474–487 (1970).

32. Siguier, P., Gourbeyre, E. & Chandler, M. Bacterial insertion sequences: their genomic impact and diversity. FEMS Microbiol. Rev. 38, 865–891 (2014).

33. Hackl, T., Duponchel, S., Barenhoff, K., Weinmann, A. & Fischer, M. G. Endogenous virophages populate the genomes of a marine heterotrophic flagellate. doi:10.1101/2020.11.30.404863.

34. Johnson, C. M. & Grossman, A. D. Integrative and Conjugative Elements (ICEs): What They Do and How They Work. Annu. Rev. Genet. 49, 577–601 (2015).

35. Sobecky, P. A. & Hazen, T. H. Horizontal gene transfer and mobile genetic elements in marine systems. Methods Mol. Biol. 532, 435–453 (2009).

36. Berube, P. M. et al. Physiology and evolution of nitrate acquisition in Prochlorococcus. ISME J. 9, 1195–1207 (2014).

37. Martiny, A. C., Coleman, M. L. & Chisholm, S. W. Phosphate acquisition genes in Prochlorococcus ecotypes: evidence for genome-wide adaptation. 103, 12552–12557 (2006).

38. Barnett, J. P. et al. Mining genomes of marine cyanobacteria for elements of zinc homeostasis. Front. Microbiol. 3, 142 (2012).

39. Martínez, A., Osburne, M. S., Sharma, A. K., DeLong, E. F. & Chisholm, S. W. Phosphite utilization by the marine picocyanobacterium Prochlorococcus MIT9301. Environ. Microbiol. 14, 1363–1377 (2012).

40. Maiques, E. et al. Role of Staphylococcal Phage and SaPI Integrase in Intra- and Interspecies SaPI Transfer. J. Bacteriol. 189, 5608–5616 (2007).

41. Novick, R. P., Christie, G. E. & Penadés, J. R. The phage-related chromosomal islands of Gram-positive bacteria. Nat. Rev. Microbiol. 8, 541–551 (2010).

42. Fillol-Salom, A. et al. Phage-inducible chromosomal islands are ubiquitous within the bacterial universe. ISME J. 12, 2114–2128 (2018).

43. Lobb, B., Tremblay, B. J.-M., Moreno-Hagelsieb, G. & Doxey, A. C. An assessment of genome annotation coverage across the bacterial tree of life. Microb Genom 6, (2020).

44. Berube, P. M., Rasmussen, A., Braakman, R., Stepanauskas, R. & Chisholm, S. W. Emergence of trait variability through the lens of nitrogen assimilation in Prochlorococcus. Elife 8, (2019).

45. Smyshlyaev, G., Barabas, O. & Bateman, A. Sequence analysis allows functional annotation of tyrosine recombinases in prokaryotic genomes. bioRxiv 542381 (2019) doi:10.1101/542381.

46. Hiramatsu, K. et al. Genomic Basis for Methicillin Resistance in Staphylococcus aureus. Infect Chemother 45, 117–136 (2013).

47. Martínez-Rubio, R. et al. Phage-inducible islands in the Gram-positive cocci. ISME J. 11, 1029–1042 (2017).

48. Achaz, G., Coissac, E., Netter, P. & Rocha, E. P. C. Associations between inverted repeats and the structural evolution of bacterial genomes. Genetics 164, 1279–1289 (2003).

49. Khil, P. P. & Camerini-Otero, R. D. Over 1000 genes are involved in the DNA damage response of Escherichia coli. Mol. Microbiol. 44, 89–105 (2002).

50. Llabrés, M. & Agustí, S. Picophytoplankton cell death induced by UV radiation: Evidence for oceanic Atlantic communities. Limnol. Oceanogr. 51, 21–29 (2006).

51. Agustí, S. & Llabrés, M. Solar radiation-induced mortality of marine pico-phytoplankton in the oligotrophic ocean. Photochem. Photobiol. 83, 793–801 (2007).

52. Kolowrat, C. et al. Ultraviolet stress delays chromosome replication in light/dark synchronized cells of the marine cyanobacterium Prochlorococcus marinus PCC9511. BMC Microbiol. 10, 204 (2010).

53. Mella-Flores, D. et al. Prochlorococcus and Synechococcus have Evolved Different Adaptive Mechanisms to Cope with Light and UV Stress. Front. Microbiol. 3, 285 (2012).

54. Iyer, V. N. & Szybalski, W. Mitomycins and Porfiromycin: Chemical Mechanism of Activation and Cross-linking of DNA. Science 145, 55–58 (1964).

55. Tomasz, M. Mitomycin C: small, fast and deadly (but very selective). Chem. Biol. 2, 575–579 (1995).

56. Turner, S. L., Bailey, M. J., Lilley, A. K. & Thomas, C. M. Ecological and molecular maintenance strategies of mobile genetic elements. FEMS Microbiol. Ecol. 42, 177–185 (2002).

57. Minoia, M. et al. Stochasticity and bistability in horizontal transfer control of a genomic island in Pseudomonas. Proc. Natl. Acad. Sci. U. S. A. 105, 20792–20797 (2008).

58. Beaulaurier, J. et al. Assembly-free single-molecule nanopore sequencing recovers complete virus genomes from natural microbial communities. bioRxiv 619684 (2019) doi:10.1101/619684.

59. Gregory, A. C. et al. Marine DNA Viral Macro- and Microdiversity from Pole to Pole. Cell 177, 1109–1123.e14 (2019).

60. Luo, E., Eppley, J. M., Romano, A. E., Mende, D. R. & DeLong, E.F. Double-stranded DNA virioplankton dynamics and reproductive strategies in the oligotrophic open ocean water column. ISME J. 14, 1304–1315 (2020).

61. Biller, S. J. et al. Bacterial Vesicles in Marine Ecosystems. Science vol. 343 183–186 (2014).

62. Moore, C. M. et al. Processes and patterns of oceanic nutrient limitation. Nat. Geosci. 6, 701–710 (2013).

63. Soler, N. & Forterre, P. Vesiduction: the fourth way of HGT. Environ. Microbiol. 22, 2457–2460 (2020).

64. Dorward, D. W., Garon, C. F. & Judd, R. C. Export and intercellular transfer of DNA via membrane blebs of Neisseria gonorrhoeae. J. Bacteriol. 171, 2499–2505 (1989).

65. Klieve, A. V. et al. Naturally occurring DNA transfer system associated with membrane vesicles in cellulolytic Ruminococcus spp. of ruminal origin. Appl. Environ. Microbiol. 71, 4248–4253 (2005).

66. Gaudin, M. et al. Extracellular membrane vesicles harbouring viral genomes. Environ. Microbiol. 16, 1167–1175 (2014).

67. Renelli, M., Matias, V., Lo, R. Y. & Beveridge, T. J. DNA-containing membrane vesicles of Pseudomonas aeruginosa PAO1 and their genetic transformation potential. Microbiology 150, 2161–2169 (2004).

68. Kolling, G. L., Simon, L. & Matthews, K. R. Vesicle-Mediated Transfer of Virulence Genes fromEscherichia coli O157: H7 to Other Enteric Bacteria. Applied and (2000).

69. Erdmann, S., Tschitschko, B., Zhong, L., Raftery, M. J. & Cavicchioli, R. A plasmid from an Antarctic haloarchaeon uses specialized membrane vesicles to disseminate and infect plasmid-free cells. Nat Microbiol 2, 1446–1455 (2017).

70. Tran, F. & Boedicker, J. Q. Genetic cargo and bacterial species set the rate of vesicle-mediated horizontal gene transfer. Sci. Rep. 7, 8813 (2017).

71. Nazarian, P., Tran, F. & Boedicker, J. Q. Modeling Multispecies Gene Flow Dynamics Reveals the Unique Roles of Different Horizontal Gene Transfer Mechanisms. Front. Microbiol. 9, 2978 (2018).

72. Kauffman, K. M. et al. A major lineage of non-tailed dsDNA viruses as unrecognized killers of marine bacteria. Nature 554, 118–122 (2018).

73. Biller, S. J. et al. Bacterial vesicles in marine ecosystems. Science 343, 183–186 (2014).

74. Hackl, T. & Ankenbrand, M. J. Supplementary code and data for “Novel integrative elements and genomic plasticity in ocean ecosystems”. Zenodo (2020) doi:10.5281/zenodo.4393629.

