## Supplement for "Novel integrative elements and genomic plasticity in ocean ecosystems"

**Materials & Methods** **3**
Genomic datasets 3
Prochlorococcus genomes 3
Global Ocean Reference Genomes (GORG) - Tropics 3
Vesicle-fraction metagenomes 3
Tara Oceans viral-fraction short-read metagenome contigs 3
Station ALOHA viral-fraction nanopore reads 4
Computational analyses 4
Gene prediction and ortholog detection 4
Phylogenetic tree reconstruction for Prochlorococcus 4
Reference-based genome-scaffolding 4
Genomic island predictions using orthogroups frequencies and a Hidden-Markov-Model 5
Estimation of homologous recombination rates for island-flanking and non-island-flanking genomic regions 5
Identification element hallmark genes in Prochlorococcus 5
Automatic detection, annotation and visualisation of integrative elements in genomic data 6
Detection of element-containing reads in viral-fraction nanopore reads 7
Functional enrichment analysis 7
Phylogeny of tyrosine integrases 7
Differential abundance analysis of element signatures in vesicle- and cellular-fraction metagenomes 7
Differential abundance analysis of tRNA attachment sites in vesicle- and cellular-fraction metagenomes 7
Oxford Nanopore genome assembly of Prochlorococcus MIT0604 cultures 8
Experimental procedures 8
Oxford Nanopore sequencing of Prochlorococcus MIT0604 cultures 8
Strain isolation and genome sequencing 8
Culture conditions 9
RTqPCR of integrase genes under various treatments 9
RNA-sequencing experiments and analyses 10
Detection of element mobility in lab isolates 10

**Supplementary Figures** **11**
Supplementary Figure 1 | HMM-based genomic island prediction using abundances of orthologous genes. 11
Supplementary Figure 2 | Distribution of core and flexible genes in the genomic backbone and islands. 12
Supplementary Figure 3 | Chromosomal organization of genomic islands and associated mobile genetic elements in Prochlorococcus. 13
Supplementary Figure 4 | An example of increasing content and structure variability in and around a genomic island. 14
Supplementary Figure 5 | Homologous recombination in island-flanking and non-island-flanking genomic regions. 15
Supplementary Figure 6 | Diversity and distribution of mobile elements across 623 Prochlorococcus genomes. 16
Supplementary Figure 7 | Tycheposons in viral particles. 17
Supplementary Figure 8 | Gene functions in elements and other island parts resemble each other. 18
Supplementary Figure 9 | Prochlorococcus MIT0604 integrative elements. 19
Supplementary Figure 10 | Whole-genome alignment of Prochlorococcus MIT0604 strains showing element-related rearrangements 20
Supplementary Figure 11 | Detection of integration and excision of elements in cultures. 21
Supplementary Figure 12 | Transcriptional response of integrase genes associated with cargo-carrying elements in Prochlorococcus MIT0604 in response to a wide range of shock treatments. 22
Supplementary Figure 13 | Effect of mitomycin C treatment on various tycheposon integrases. 23
References 24

### Materials & Methods

#### Genomic datasets

***Prochlorococcus* genomes**

For our analyses, we selected a set of 623 publicly available *Prochlorococcus* genome assemblies, 73 obtained from cultured isolates, 540 generated through single-cell sequencing and 10 extracted from metagenome assemblies (Supplementary Table 1). We quality-screened the assemblies using checkm v1.0.7[1]. The selected assemblies have minimum completeness of 25% and median completeness of 74%; isolates and single-cell genomes have less than 4% contaminations, metagenome-assembled genomes less than 8%. Raw assemblies are available through public databases such as NCBI Genbank, the reference-scaffolded assemblies with our lifted annotations are available from https://github.com/thackl/pro-tycheposons and Zendodo[2].

**Global Ocean Reference Genomes (GORG) - Tropics**

This dataset consists of 12,715 single amplified genomes (SAGs) of Bacteria and Archaea, which were obtained through a randomized cell selection from 28 globally distributed samples of tropical and subtropical, epipelagic ocean water[3]. GORG-Tropics SAGs represent all major lineages of surface ocean prokaryoplankton. Used as a reference database, GORG-Tropics recruits an average of 40% reads from tropical and subtropical epipelagic metagenomes with >95% nucleotide identity. The dataset can be accessed at https://osf.io/pcwj9 and https://www.ebi.ac.uk/ena/data/view/PRJEB33281.

**Vesicle-fraction metagenomes**

We generated paired metagenomes from the cellular and vesicle/small particle fractions of four oligotrophic water samples. All samples were collected at Station ALOHA (22.75 °N, 158 °W) in the North Pacific Subtropical Gyre on cruises HOT263 (June 2014; 5m depth) and HOT283 (April 2016; 5m, 25m, and 137m depth). Each set of samples was derived from a total of ~100-200 L of water retrieved from Niskin bottles. For cellular metagenomes, 3-6 L of water was filtered onto a 0.2 µm Sterivex filter (Millipore), preserved, and DNA later extracted as previously described[4]. The <0.2 µm fraction of the remaining water was concentrated on a 100 kDa tangential flow filter and further fractionated using an Optiprep gradient; vesicle-enriched fractions were identified, DNA outside the vesicles removed using TURBO DNAse, and the remaining DNA (presumably within vesicles) was extracted as previously described[5]. Sequencing libraries for the purified cellular and vesicle/particle-associated DNA were prepared from ~1 ng of DNA using the NextEra XT kit (Illumina) and 150+150nt paired-end sequences generated by an Illumina NextSeq 500 at the MIT BioMicro Center. All data are available from the NCBI Sequence Read Archive (SRP272691).

**Tara Oceans viral-fraction short-read metagenome contigs**

The short-read viral data were obtained from 131 samples collected during the *Tara* Oceans and *Tara* Oceans Polar Circle expeditions and span water layers from the surface (5m deep) to the mesopelagic ocean (up to 1000m deep). The full description of the collection locations and depths as well as the viral enrichment, DNA sequencing, and contig assembly protocols for these samples can be found in the methods section of the Global Ocean Viromes 2.0 (GOV2.0) dataset (Gregory et al. 2019). GOV2.0's deep ocean samples from the Malaspina expedition (n=14) were not included in this study. All the assembled contigs from the 131 *Tara* samples were screened for tycheposons prior to any bioinformatic viral selection carried out in (Gregory et al. 2019) to establish the GOV2.0 dataset. These assembled contigs can be accessed through the Data Commons portal of iVirus under GOV2.0 (filename: Tara_assemblies.tar.gz)

**Station ALOHA viral-fraction nanopore reads**

The 25 m deep sample was collected on the HOT-314 cruise on August 5, 2019 at Station ALOHA (22°45' N, 158° W; http://hahana.soest.hawaii.edu/hot/). Sample collection, filtration, viral concentration, extraction, and sequencing have all been previously described[6]. Briefly, the seawater was pre-filtered by peristatic pumping through a 0.22 um filter (Sterivex GV) then concentrated by tangential flow filtration (TFF) over a 30 kDa filter (Biomax 30 kDa) membrane, catalogue #: P3B030D01, Millipore). Following multiple rounds of centrifugal concentration, lysis and DNA purification were performed in a single tube using the Qiagen Genomic-tip 20/G protocol following manufacturer's recommendations. Virus-enriched samples from 110 L of 0.22um pre-filtered seawater yielded a total of 3.2 µg of purified, high molecular weight DNA. Sequencing was conducted on a GridION X5 with FLO-MIN106 (R 9.4.1) flowcells (Oxford Nanopore Technologies, Ltd.). The resulting 701,515 reads were basecalled using Guppy v3.0.4, generating 10.38Gb of sequencing data with a read N50 length of 29.70 Kb. All data are available from the NCBI Sequence Read Archive (SRX7079550).

#### Computational analyses

**Gene prediction and ortholog detection**

To ensure consistent gene annotations, all 623 *Prochlorococcus* genomes (see Genomic datasets for details) were reannotated using PROKKA v1.12-beta[7]. The annotated genes were clusters into groups of orthologs using panX sha-0a4dfce[8] with customised settings to account for the incompleteness of the single-cell genomes in the collection: core genes are defined by being present in at least 70% of all genomes and in every genome of >98% completeness (-cg 0.70 -csf strains_complete_98plus.txt).

**Phylogenetic tree reconstruction for *Prochlorococcus***

The phylogenetic reference tree for all 623 *Prochlorococcus* genomes analysed was constructed using the maximum likelihood method from 109 concatenated single-copy core proteins of all. The markers were identified using HMMER3 v3.2.1[9] based on the 120 TIGRFAM[10] profiles previously described as ubiquitous bacterial single-copy core genes ("bac120")[11]. 11 proteins found to be not single-copy in this specific data set were excluded. Sequence files were manipulated with seqkit[12] and individual protein alignments were generated with mafft v7.310[13], trimmed with trimAl v1.4[14] (-gappyout), and concatenated with msa-concatenate (https://github.com/thackl/phylo-scripts/, sha-2334c67). The maximum likelihood phylogeny was inferred with FastTree v2.1.10[15]. The tree was rooted at the LLIV grade and visualised using the R packages phytools[16] and ggtree[17].

**Reference-based genome-scaffolding**

All 34 complete genomes comprising only a single contig were considered as references. For better comparability, all reference genomes were oriented and rotated - they are circular chromosomes - to start with the origin of replication near the dnaN gene on the plus strand. This is consistent with the convention used for most published *Prochlorococcus* genomes. OriC prediction was performed with the command line version of Ori-Finder v1.0 [18] with dnaA box sequence set to "TTTTCCACA" as suggested for cyanobacteria[19]. The start positions of 11 genomes were adjusted, and 3 of them were also reverse complemented. For all other genomes, the closest reference genome was determined by the smallest cophenetic distance in the reference phylogenetic tree described above. For each pair of draft genome and closest reference, 1x1 anchor maps were created by matching genes belonging to the same cluster of orthologous genes (see above). These anchor maps were used to orient and order contigs in a way that maximises collinearity using ALLMAPS v0.7.7 [20]. If necessary the contig spanning the beginning and end of the reference sequence was split into two parts at the position of the dnaN gene. Gaps between contigs were estimated by minimising the sum of absolute distances between corresponding genes from the 1x1 map. Contigs were weighted by the log of their length and minimum gap size was set to 100bp. The optimisation was implemented using the "L-BFGS-B" method in R v3.5.1[21]. See Prochlorococcus genomes for access to the data.

**Genomic island predictions using orthogroups frequencies and a Hidden-Markov-Model**

To automate the annotation of genomic islands across the entire dataset, we devised an HMM-based approach that uses frequencies of orthologous genes as input. We define the frequency of a gene and its respective orthogroup, as the number of strains the gene is present in. Duplications within the same genome are ignored. We are working off the observation that islands are enriched in non-core genes, and hence their composition in terms of gene frequencies is different from non-island regions[22]. We used previously described genomic islands from 2 *Prochlorococcus* HLI strains (MED4, MIT9515), 2 *Prochlorococcus* HLII strains (MIT9312, MIT9215), 1 *Prochlorococcus* LLII strain (SS120) and 1 *Prochlorococcus* LLIV strain (MIT9313)[22–24] (Supplementary Table 2) to generate four profiles of the island and non-island gene frequencies. Based on comparisons of these profiles we then defined different states for the HMM: core, flex, and inconclusive depending on their prevalence in or outside of islands. Using the R-package HMM v1.0[25] we built four HMMs with two hidden states (island, non-island). Start, transition, and emission probabilities were estimated from the literature annotations. For all genomes, each gene was assigned the category (core, flex, and inconclusive) depending on the gene frequency as described above. Using these labels in the order they appear on the scaffolded sequence (see above) as observations the Viterbi algorithm was applied to predict the hidden state for each gene (island, non-island). This gene-level resolution of genomic islands was used in further analyses.

We first reference-scaffolded and reorganised all assemblies to obtain single-chromosome scaffolds with consistent start and orientation. Using coordinates of known genomic islands[22–24] and abundances of orthologous gene clusters[8] we trained a Hidden-Markov-Model to predict islands across all genomes (Extended Data Fig. 1).

**Estimation of homologous recombination rates for island-flanking and non-island-flanking genomic regions**

Genomic islands formation is often driven by homologous recombination between flanking core genes[26,27]. To test if this is also the case for the islands in *Prochlorococcus* we estimated recombination rates for genomic regions relative to their distance to genomic islands. First, we identified backbone core genes (orthogroups not found in islands) individually for the 5 large monophyletic clades HLI, HLII/VI, LLI, LLII/III and LLIV. We then estimated a proxy for the average distance of each cluster to the closest island across the entire clade. For that, we took the 25% quantile of all gene-to-closest-island distances we obtained from each individual genome. Next, we generated protein alignments for all of those clade-wise clusters with mafft v7.310[13], mapped back the nucleotide codons onto the amino-acid alignments (msa-codon https://github.com/thackl/phylo-scripts/) and trimmed positions with more than 70% gaps with trimAl v1.4[14] (-gt .3). We then concatenated all alignments ordered by their estimated distance to the closest island, with island-flanking genes at the beginning and genes furthest from islands at the end. Finally, we partitioned those clade-wise concatenated alignments into 99999 nucleotide long blocks. For each of those blocks, we estimated recombination rates and related variables using mcorr v20180102[28], and compared those block-wise results with respect to the proximity of contained genes to genomic islands.

**Identification element hallmark genes in *Prochlorococcus***

To enable a comprehensive Prochlorococcus-focused search for integrative elements we compiled and curated protein HMM-profiles of genes typically found in candidate elements using an iterative, explorative approach: Starting with a handful of manually annotated high-confidence element candidates we identified relevant orthogroups (see Gene prediction, ortholog detection and functional annotation) based on one of the two following criteria:

a) orthogroups with some annotations associated with the excision-replication-packaging life-cycle typical for elements (integrases, primases, helicases, capsid genes, terminases), and
b) orthogroups appearing in multiple candidates with syntenically conserved patterns.

From these orthogroups we created and curated, redundancy-reduced alignments and HMM-profiles using combinations of the following tools: mafft v7.310[13], msa-trim (https://github.com/thackl/phylo-scripts/, sha-2334c67), AliView[29], FastTree v2.1.10[15], FigTree[30], and HMMER3 v3.2.1[9]. Orthogroups with good reciprocal hits in all-versus-all comparisons and overall consistent multiple sequence alignment, when aligned all together, were merged. We then scanned all annotated proteins of our collection of 623 Prochlorococcus genomes using these profiles and rudimentary versions of the R-scripts described in more detail in the next section. We identified, ranked and visualized genomic regions with multiple hits to different hallmark genes within close proximity to each other (multiple hits within a 10-20kb window). From the thus obtained highest ranking clusters we selected new high-confidence candidates and used those together with previously selected candidates to repeat the identification of hallmark genes, expand the overall size of the gene set and refine the existing profiles.

In addition to curating profiles for proteins often found in putative elements in *Prochlorococcus*, we also added generic Pfam profiles [31] with a strong overlap to some of the other hallmark profiles and gathered proteins from four sets of published PICI elements [32–35]. We grouped these genes into clusters based on the annotations provided in the respective publications, manually curated the alignments, and merged clusters into single profiles if they had good reciprocal hits in all-vs-all comparisons and consistent multiple sequence alignment when aligned together.

**Automatic detection, annotation and visualisation of integrative elements in genomic data**

Ultimately we devised a small pipeline to automate the detection of elements in genomic data sets. The scripts are available at https://github.com/thackl/pro-tycheposons/element-finder or Zendodo[2], and perform the following steps:

- Gene prediction with prodigal[36] if no annotations are provided
- Full-length tRNA annotation with ARAGORN[37] if no external tRNA database is provided
- Detection of element hallmark genes and viral hallmark genes (VirSorter profiles[38]) using HMMER3
- Partial tRNA annotations using full-length tRNAs and BLAST+[39]
- Attachment site detection in integrase-flanking regions with BLAST+
- Scoring of candidate elements based on the presence of different hallmark genes, attachments sites and tRNAs
- Visualisation of elements using ggplot and https://github.com/thackl/gggenomes

For this study, we analysed three datasets with this pipeline:

1) 623 Prochlorococcus genomes
2) 2344 single-cell assemblies of the Global Ocean Reference Genomes (GORG)
3) up to 1000 randomly selected 5-20 kbp long contigs from 262 different viral-fraction samples from Tara Oceans (128,566 contigs in total)

For more information on the datasets, see the section Genomic datasets, The results from the analyses are available at https://github.com/thackl/pro-tycheposons or Zenodo[2].

**Detection of element-containing reads in viral-fraction nanopore reads**

To annotate element hallmark genes in raw nanopore reads, which have error-rates too high to predict open reading frames, we used a different strategy: We converted the alignments used to generate the HMM-profiles for hallmark genes into a protein reference database and used DIAMOND[40] in long-read mode to align nanopore reads to this database. We then identified and visualised reads with multiple hits to different hallmark genes.

**Functional enrichment analysis**

For the functional enrichment analysis, all genes were split into three sets: backbone, island, elements. Detected hallmark genes and those smaller than 201bp were excluded. The sets are mutually exclusive so genes within elements within islands are only assigned the element category. Pairwise functional enrichment analyses were performed between backbone-element and between backbone-island using the R-package topGO v2.34.0[41]. Significantly enriched GO terms ($p<1e-10$, only GO terms with at least 100 occurrences are considered) in the two analyses were compared using GOView[42] for each of the GO categories: molecular function, cellular component and biological process.

**Phylogeny of tyrosine integrases**

To put the tyrosine recombinases identified on most of the new elements into context with previously described integrases, we first compiled a representative set of known recombinase proteins. We used a HMM database of recently described Xer-like tyrosine-recombinases[43], which was kindly provided to us by the authors, to collect protein sequences from UniRef50 (https://www.uniprot.org/help/uniref). We then combined these sequences with a non-redundant subset of the integrase sequences found in our elements (max pairwise identity of 40%). We aligned the sequences with mafft v7.310[13] (--genafpair), computed a phylogenetic tree with FastTree v2.1.10[15] and visualised it with iTOL[44] followed by manual curation. For the comparison of the phylogenetic diversity contributed by the new integrases, we divided the sum of branch length of the new integrase clade by the total sum of branch lengths in the full tree. Before classifying new putative integrases as such, we also checked each subtype alignment for the presence of the characteristic residues at the catalytic sites to gain more confidence in the prediction of their functionality.

**Differential abundance analysis of element signatures in vesicle- and cellular-fraction metagenomes**

To assess the relative abundance of element hallmark genes in cellular- and vesicle-fraction metagenomes, we recruited reads translated into amino-acid space using orfm v0.7.1[45] with HMMER3 (--evalue 1e-20) to all element hallmark gene profiles and 109 *Prochlorococcus* core gene profiles. See Phylogenetic tree reconstruction for *Prochlorococcus* for details on the generation of the single-copy core marker gene set. We then analysed the obtained counts for differential abundance with edgeR[46]. We only considered profiles with a minimum count of 5 in at least two cellular- and two viral-fraction samples. In the absence of replicates, we estimated the dispersion from all core genes across all samples, assuming that this would provide us with a reasonable yet rather conservative estimate. We normalised for library size (method="TMM") and tested for differences between the two fractions using the exact test and the dispersion estimated from the core genes.

**Differential abundance analysis of tRNA attachment sites in vesicle- and cellular-fraction metagenomes**

To assess the abundance of tRNA sequences that might serve as attachment site in the vesicle- and cellular-fraction samples we applied a two-step process: First, we screened for reads with at least one almost exact 39bp match using bbduk[47] (k=39 edist=2) to a non-redundant reference database of marine tRNA genes we compiled using the GORG single-cell genomes (see Global Ocean Reference Genomes (GORG)). We then blasted those reads with settings optimised for short almost exact matches (-task blastn -reward 1 -penalty -4 -gapopen 5 -gapextend 2 -perc_identity 94 -evalue 10e-5) against both 5' and 3' halves of all reference tRNAs. We then further analysed the resulting counts in R: we filtered for a minimum alignment length of 38bp and tested for tRNAs with differentially abundant 5' and 3' regions with Fisher's exact test and the Bonferroni correction to adjust p-values for multiple testing. The resulting count distributions were visualized with ggplot2[48].

**Oxford Nanopore genome assembly of *Prochlorococcus* MIT0604 cultures**

The assemblies of two independently growing cultures of MIT0604 were generated using wtdbg2 sha-8926622 from nanopore reads longer than 50,000 bp for the nitrate-culture and 30,000 bp for the ammonia-culture, respectively. In both cases, a single contig matching the complete chromosome of MIT0604 was obtained and extracted for further analysis from the assembly. The contigs were each reorganised to match the start and orientation of the reference Illumina assembly generated in 2011. The assemblies and read data set were analysed for rearrangements using Geneious[49], Mauve[50], minimap2[51] and Ribbon[52]. The processed assemblies are available for download at https://github.com/thackl/pro-tycheposons.

#### Experimental procedures

##### Oxford Nanopore sequencing of *Prochlorococcus* MIT0604 cultures

Two independent cultures of *Prochlorococcus* MIT0604 were sequenced using Oxford Nanopore technologies. Samples were prepared following an ultra-long read sequencing protocol[53], and whole genomes were sequenced using the Rapid sequencing kit (SQK-RAD004) on a MinION sequencer according to the manufacturer's instructions (Oxford Nanopore).

##### Strain isolation and genome sequencing

*Prochlorococcus* strain MIT1013 was isolated from seawater obtained on the BiG-RAPA (Biogeochemical Gradients – Role in Arranging Planktonic Assemblages) expedition aboard the R/V Melville (MV1015) during the late austral spring of 2010 (18 November 2010 – 14 December 2010). Seawater was collected at Station 7 (Latitude: -26.25; Longitude -104) using a Niskin bottle rosette (Cast 68, 10 December 2010, 15:03 GMT) from a depth of 150m, corresponding to the subsurface chlorophyll maximum. Seventeen mL of seawater was aliquoted into an acid-washed 28 mL screw cap polycarbonate tube and amended with 20 µM ammonium chloride, 1 µM sodium phosphate, 1 mM sodium bicarbonate, 0.117 µM ethylenediaminetetraacetic acid, 0.117 µM iron (III) chloride, 0.009 µM manganese (II) chloride, 0.0008 µM zinc (II) sulfate, 0.0005 µM cobalt (II) chloride, 0.0003 µM sodium molybdate, 0.001 µM sodium selenite, and 0.001 µM nickel (II) chloride. The MIT1013 strain has been deemed unialgal based on observations of a single *Prochlorococcus* population using flow cytometry and by the presence of a single 16S–23S rRNA internal transcribed spacer (ITS) sequence as determined by direct sequencing of its ITS PCR amplicon. The complete genome sequence of MIT1013 was additionally determined. Cells were grown to mid-exponential phase and pelleted by centrifugation. DNA was isolated by phenol/chloroform extraction[54]. PacBio library preparation and sequencing was carried out by the MIT BioMicro Center and the UMass Worcester Medical School's Deep Sequencing Core Facility. Assembly of PacBio reads was performed using the hierarchical genome assembly process (Protocol = RS_HGAP_Assembly.2) as implemented in SMRT Analysis 2.3.0 (Chin et al. 2013) with the following parameters adjusted: Minimum Polymerase Read Quality = 0.85 and Genome Size = 2000000 bp (default settings were used for all other parameters). A single *Prochlorococcus* contig was identified as well as a 6513 bp contig most closely related to *Marinobacter* sp., a common heterotrophic contaminant of xenic cultures of *Prochlorococcus*. Overlapping ends of the *Prochlorococcus* contig were identified using BLAST, and the assembled contig was manually circularized. The circular assembly was corrected using the RS_Resequencing.1 protocol in SMRT Analysis 2.3.0 (Chin et al. 2013) with the following parameters: Minimum Polymerase Read Quality = 0.85 and Consensus Algorithm = Quiver. This genome was deposited with IMG (accession number 2681812904), annotated using IMG Annotation Pipeline version 4[55,56], and included in ProPortal CyCOGs 6.0[57].

##### Culture conditions

*Prochlorococcus* cells (axenic cultures, except for MIT1013 and PAC1, and MIT0604 when specified) were grown under constant light flux (30–40 µmol photons $m^{-2}$ $s^{-1}$) at 24°C in natural seawater-based Pro99 medium containing 0.2-µm-filtered Sargasso Sea water, amended with Pro99 nutrients (N, P, and trace metals) prepared as previously described[58]. Growth was monitored using bulk culture fluorescence measured with a 10AU fluorometer (Turner Designs).

**Supplementary Tab. 12.** *Prochlorococcus* isolate strains used in this study. HOTS: Hawaii Ocean Time Series station, Pacific oligotrophic gyre. BATS: Bermuda Atlantic Time Series station, Sargasso Sea.

| Strain | Clade | Location of isolation, depth | Reference |
|---|---|---|---|
| MIT0604 | HLII | HOTS, 175m | Biller et al. 2014[59] |
| MIT9202 | HLII | Tropical Pacific, 79m | Thompson et al. 2011[60] |
| MIT9312 | HLII | Gulf Stream, 135m | Coleman et al. 2006[61] |
| MIT9215 | HLII | Equatorial Pacific, surface | Kettler et al. 2007[62] |
| SB | HLII | Western Pacific, 40m | Shimada et al. 1995[63] |
| PAC1 | LLI | HOTS, 100m | Penno et al. 2000[64] |
| MIT1013 | LLI | Eastern South Pacific Subtropical Gyre, 150m | This study |
| MIT1306 | LLIV | HOTS, 150m | Cubillos-Ruiz et al. 2017[65] |

**RTqPCR of integrase genes under various treatments**

Reverse-Transcription qPCR analysis of integrase genes was performed on biological triplicates exposed to the treatment, compared to untreated controls. For RNA preparation, cells were collected by centrifugation (12,000g for 12 min, 20°C) and immediately resuspended in 500 µL of TRI reagent (Zymo Research). RNA was isolated using the Direct-zol RNA MicroPrep kit (Zymo Research) according to the manufacturer's instructions. A DNA removal step was added after elution using the TURBO DNA-free Kit (ThermoFisher). RNA samples were subsequently concentrated using RNA the Clean & Concentrator kit (Zymo Research), followed by reverse transcription using the SuperScript™ III First-Strand Synthesis System (ThermoFisher). Finally, triplicate qPCRs were performed using the QuantiTect Probe PCR Kit (Qiagen) - using the primer sets detailed in STab. 11 - and differential gene expression was calculated following the comparative CT (2-DDCT) method[66], with the expression of gene *rnpB* as the endogenous reference.

Description of the treatments from SFig. 4 graph: 'Alteromonas' = addition of the helper strain Alteromonas MIT1002 at $5x10^6$ cells $mL^{-1}$ for 1h ; 'Pyruvate' = addition of 5 mM pyruvate for 2h; 'Glucose' = addition of 5 mM glucose for 2h; 'Nitrate' = culture grown in Pro99 media with nitrate substituted to ammonium as the nitrogen source; 'N starvation' = cells are washed twice and resuspended in Pro99 media devoid of any nitrogen source, and incubated for 48h, while control cultures are washed but resuspended in replete Pro99; 'P starvation' = same process, using Pro99 media devoid of phosphate; 'Stationary phase' = cultures are left until they reach the stationary phase, while control cultures are harvested in exponential phase; 'Metal toxicity' = trace metals present in Pro99 are added at 5 times their concentration (toxic level) for 1h; 'Copper' = addition of $CuCl_2$ at 100 pM in the culture for 1h; 'Arsenate' = addition of $Na_3AsO_4$ at 10 µM for 16h; '$H_2O_2$' = Addition of hydrogen peroxide at 0.5 µM for 1h; 'DCMU' = addition of DCMU (Diuron herbicide) at 4 µM for 1h; 'Chloramphenicol' = addition of chloramphenicol at 1 µM (lethal dose) for 1h; 'Ciprofloxacin' = addition of ciprofloxacin at 1 µM (lethal dose) for 1h[67]; 'Mitomycin C' = addition of mitomycin C at 20 µM (lethal dose) for 2h; 'Cold shock' = cultures were cooled to 14°C for 1h; 'pH 9.3' = addition of a 0.1 M NaOH solution at 1.4 mM to reach pH 9.3, for 1h; 'pH 6.5' = addition of a 0.1M HCl solution to 2.1 mM to reach pH 6.5, for 1h; 'Dark' = culture tubes are placed in dark for 1h; 'High Light' = culture tubes are placed at a light intensity of 150 µE for 1h; 'UV shock' = cells were irradiated at for 30 sec at 254 nm (UV 100 µW $cm^{-2}$) in an uncovered sterile glass petri dish (150 mm diameter) containing 25 mL of culture. Cells were placed back in 25 mL culture tubes inside the incubator for 30 min, while control cultures were subjected to the same conditions, omitting the UV irradiation[68]; 'UV acclimation' = Cultures were acclimated to a light level of 50 µE in diel regime, receiving an extra 2h daily UV at midday from a Rayminder 15W UV lamp shining 302-316 nm UVB (16 in distance from lamp). Control cultures were grown in diel regime at 25 µE light intensity and without UV. RNA was harvested at midday in exponentially growing cultures[69]; 'Exogenous DNA' = 1 µg of pUC19 plasmid DNA was electroporated in concentrated cell samples[67], which were then left to recover for 24h before RNA extraction. The control cultures were also electroporated in the absence of plasmid DNA.

**RNA-sequencing experiments and analyses**

Nine 30 mL cultures of Prochlorococcus MIT0604 strain were grown to exponential phase. 3 cultures were treated with mitomycin C (Sigma-Aldrich) at a final concentration of 15 µg $mL^{-1}$ for 2 hours; 3 cultures were applied a UV shock by irradiating cells at room temperature for 30 s at 254 nm (UV 100 µW $cm^{-2}$) in an uncovered sterile glass Petri dish (150 mm diameter) - then placed back into their initial culture condition for 1h[68]; the 3 remaining cultures were kept as control with no treatment. Cells were harvested by centrifugation at 12,000 g for 12 min, 20°C and RNA was extracted using the mirVana microRNA (miRNA) extraction kit (Ambion, Carlsbad, CA, USA). All strand-specific transcriptome sequencing (RNA-seq) libraries were constructed using the KAPA RNA HyperPrep kit (Illumina) and used the RiboZero kit (Illumina) for ribosomal RNA depletion. Sequencing was carried out on an Illumina NextSeq 500 instrument at the BPF Next-Gen Sequencing Core Facility at Harvard Medical School, with a High-Output 75-cycle kit to obtain Single-Read 75bp reads.

Adapters were trimmed from the raw Illumina data with bbduk v38.16[70], with settings ktrim=r, k=23, mink=11, hdist=1. Low-quality regions were removed from the adapter-trimmed sequences using bbduk v38.16[70], with parameters qtrim=rl, trimq=6. The trimmed RNA-seq reads were aligned to a reference file containing the MIT0604 genome (available from https://github.com/thackl/pro-tycheposons/) with the Burrows-Wheeler Aligner v0.7.16a-r1181[71], using the BWA-backtrack algorithm. To determine the number of reads that aligned to each annotated ORF in the "sense" and "antisense" orientations, we parsed the mappings using the HTSeq package v0.11.2[72] with default parameters and the "nonunique all" option. We compiled the counts of reads that aligned to each ORF (excluding rRNAs and tRNAs because library preparation included ribosomal depletion) across replicates. We identified differentially expressed genes using the DESeq2 R package v1.24.0[73]. Using the standard DESeq2 functions and workflow, we normalised samples by library sequencing depth and estimated the dispersion of each gene. Differential expression tests were performed on mitomycin C vs. control and UV shock vs. control comparisons with the Wald test, using a negative binomial generalised linear model. P-values were corrected for multiple testing with the Benjamini–Hochberg procedure. As was suggested by the DESeq2 authors[73], genes with an adjusted p-value of <0.1 were considered to have significantly different expression between a given pair of treatments. We visualised the differential expression results with ggplot2[48].

**Detection of element mobility in lab isolates**

Excision of elements in lab isolates was probed using end-point PCR specific to the chromosomal vacated site ('excised'), or the circular/tandem repeats state of the elements (SFig. 7). Primers are listed in Stab. 11. 25 mL duplicate cultures were grown to exponential phase and sampled at $t_0$ (before the addition of mitomycin C) and $t_{2h}$ (2h post-addition of mitomycin C at 10 µg $mL^{-1}$ final concentration). For sampling, 10 mL of cells were harvested by centrifugation (7,000 g for 30 min, 20°c), and DNA was extracted using the DNeasy Blood & Tissue kit (Qiagen). PCR reactions were performed using the Quick Load Taq 2X master mix (New England Biolabs) with 2 min elongation time and 52°C annealing temperature. PCR products were directly loaded on 1% agarose gels for visualisation. Each band displayed on the figure was checked by Sanger sequencing and corresponded to the expected amplicon.

### Supplementary Figures

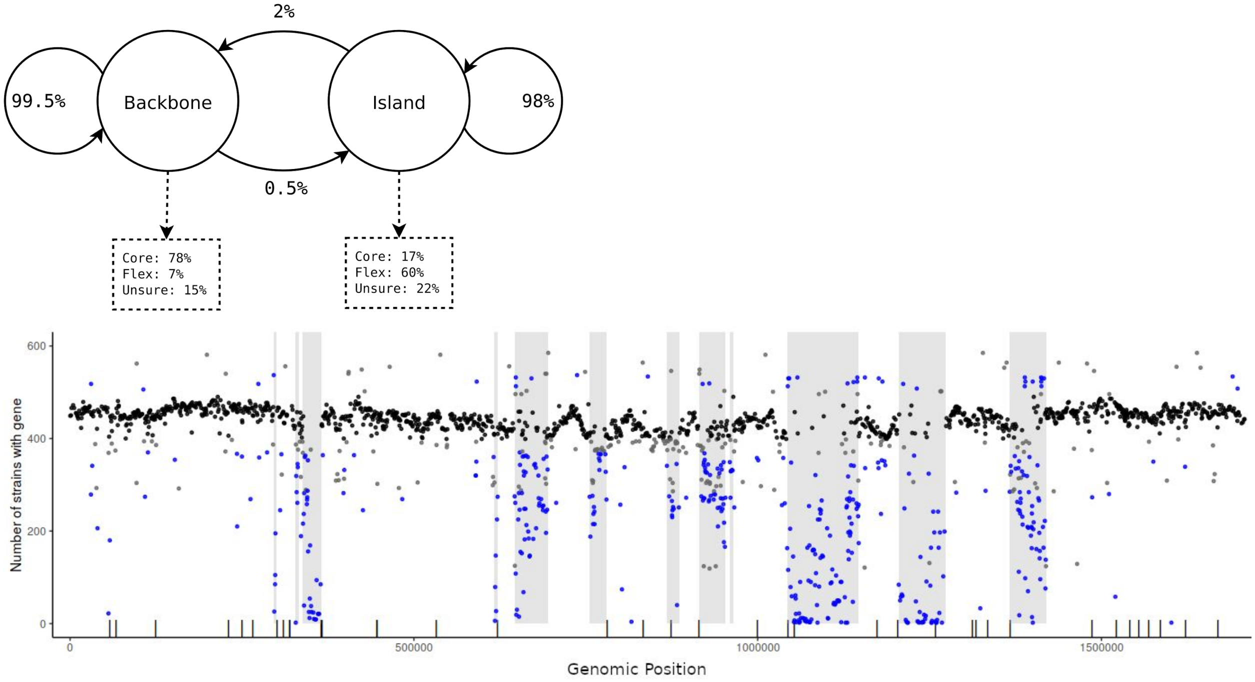


**Supplementary Figure 1 | HMM-based genomic island prediction using abundances of orthologous genes.**
Top: the two-state Hidden Markov Model to predict genomic islands with exemplary transition and emission possibilities. Bottom: result of the prediction on strain *Prochlorococcus* MIT9312 (HLII). Each gene is depicted as a dot at its genomic position with the number of strains that possess this gene on the y-axis. Genes are coloured by their class (core: black, flex: blue or unsure: grey). The sequence of genes and their class is fed to the clade-specific HMM as observations and hidden states are predicted through the Viterbi-algorithm. Resulting island regions are depicted as vertical grey bars. Additionally, the location of tRNA genes is shown as black ticks on the x-axis.

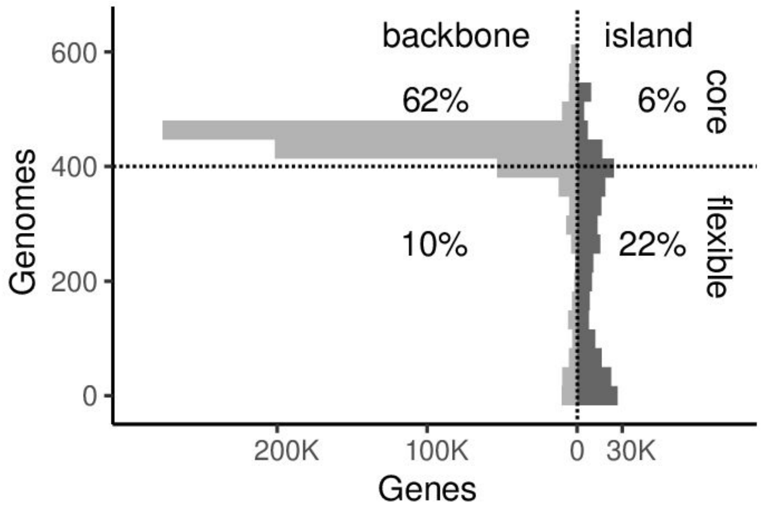


**Supplementary Figure 2 | Distribution of core and flexible genes in the genomic backbone and islands.**
A histogram showing the distribution of genes contained in the genomic backbone and genomic islands with respect to the number of genomes the respective genes are observed in (y-axis). Overall we analysed 623 genomes, with median completeness of 74%. For the purpose of this summary and to account for the incompleteness of the genomes and statistical variance, we define core genes as genes present in at least 65% of the genomes (dotted line). Based on this definition more than ⅔ of *Prochlorococcus* flexible genes are contained in genomic islands we annotated.

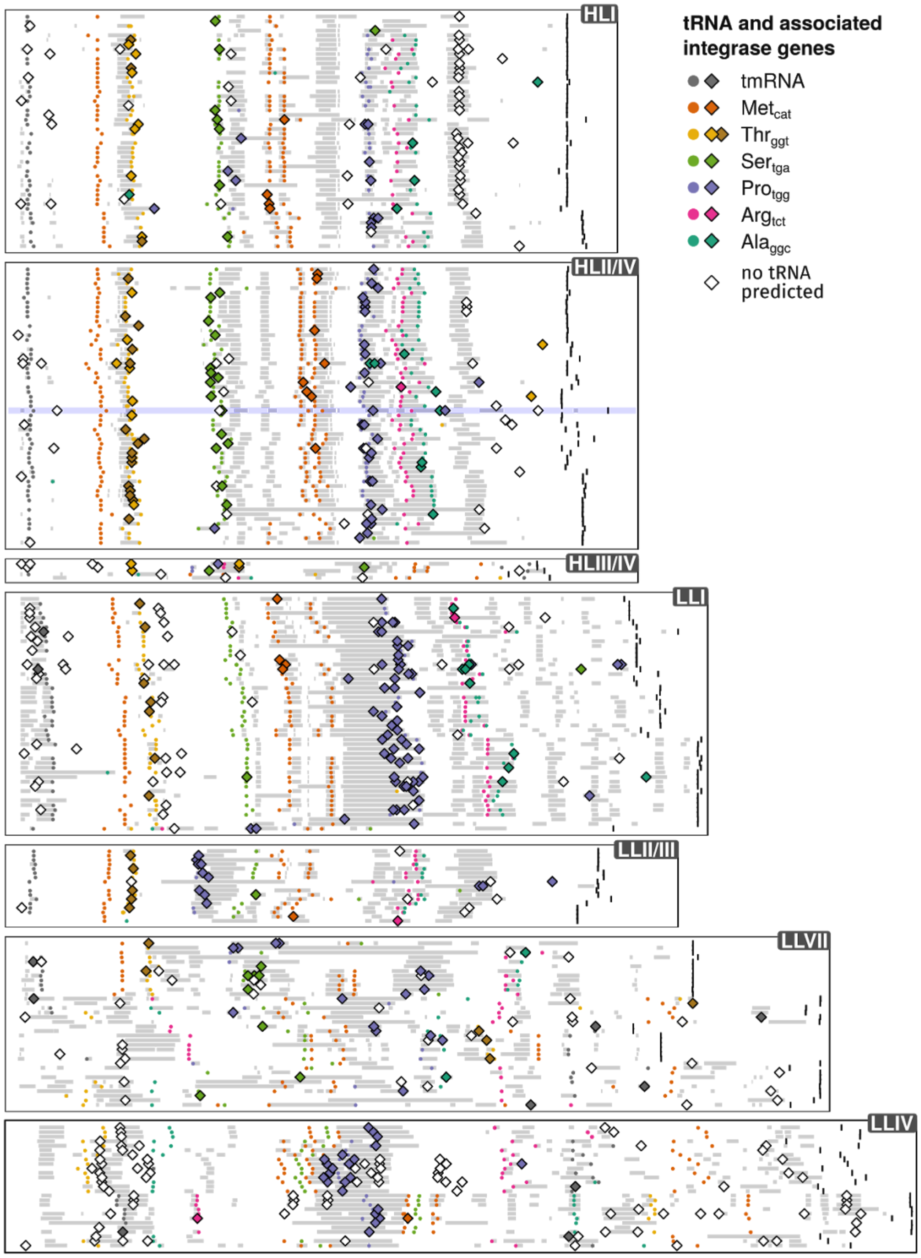


**Supplementary Figure 3 | Chromosomal organization of genomic islands and associated mobile genetic elements in *Prochlorococcus*.**

237 finished or reference-scaffolded circular *Prochlorococcus* shown relative to their origin of replication (left to rightmost black mark). Vertical column-like features indicate the predicted genomic islands in conserved locations across the genomes (grey bars). Most islands are associated with one or two specific full-length tRNA genes (colored points. These tRNAs are targeted by mobile genetic elements carrying integrases specific to the different islands (colored diamonds). Only the 50 most complete genomes of each clade out of all the 623 genomes used in this study are shown. The genomes are ordered according to their phylogenetic relationships, i.e. the most closely related genomes are plotted next to each other, and are grouped in different panels corresponding to known *Prochlorococcus* clades and grades of the low-light- (LL) and high-light-adapted (HL) ecotypes. The genome of strain MIT0604 used for experimental analyses is highlighted in blue.

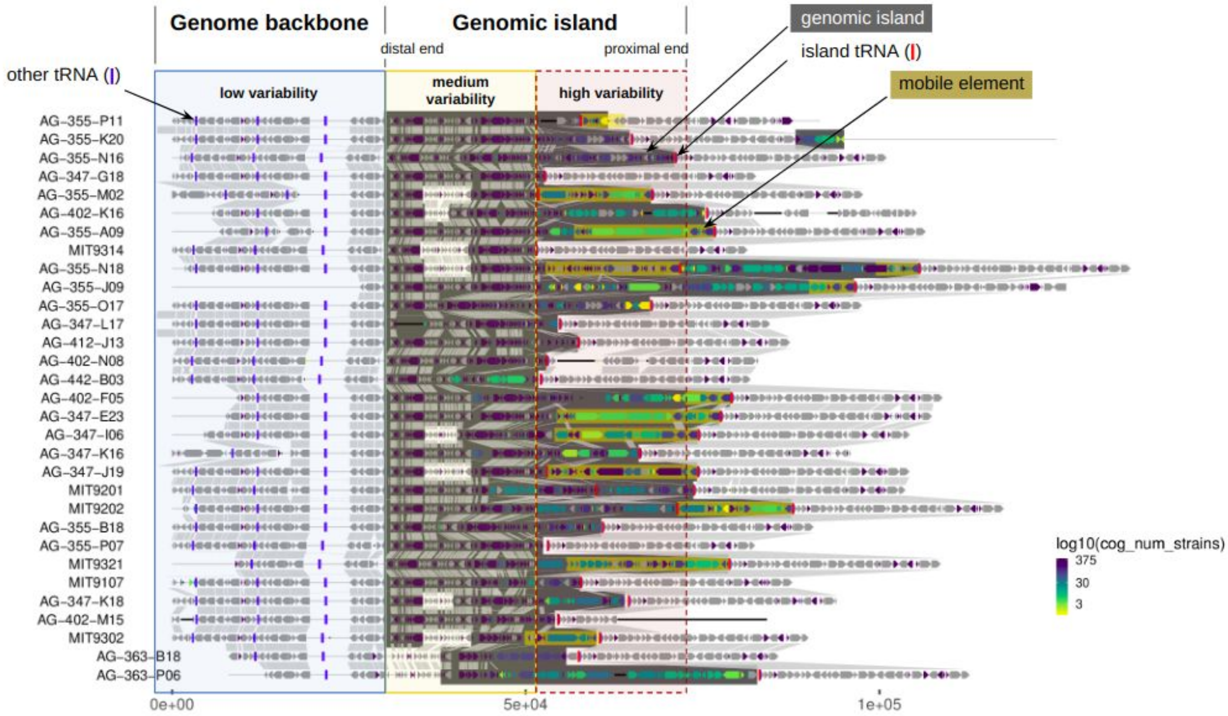


**Supplementary Figure 4 | An example of increasing content and structure variability in and around a genomic island.**

A gene synteny map showing the genomic regions around the Threonine$_{ggt}$-island in a selection of HLII-*Prochlorococcus* strains. Each row is a genome. Filled arrows indicate protein-coding genes, colours their abundance in the set of 623 genomes (grey: core genome). Orthologous genes are linked by vertical grey bars. Red and blue vertical bars indicate tRNA genes, with the red tRNA being the integration site for mobile elements (dark yellow boxes) "visiting" the island. The three large blocks highlight regions of increasing genomic variability, from the most conserved backbone genome (blue), through the distal end of the island with medium variability (yellow, furthest from island-tRNA), to the proximal end with high variability and in some cases integrated mobile elements (red, abutting Thr$_{ggt}$-tRNA gene)

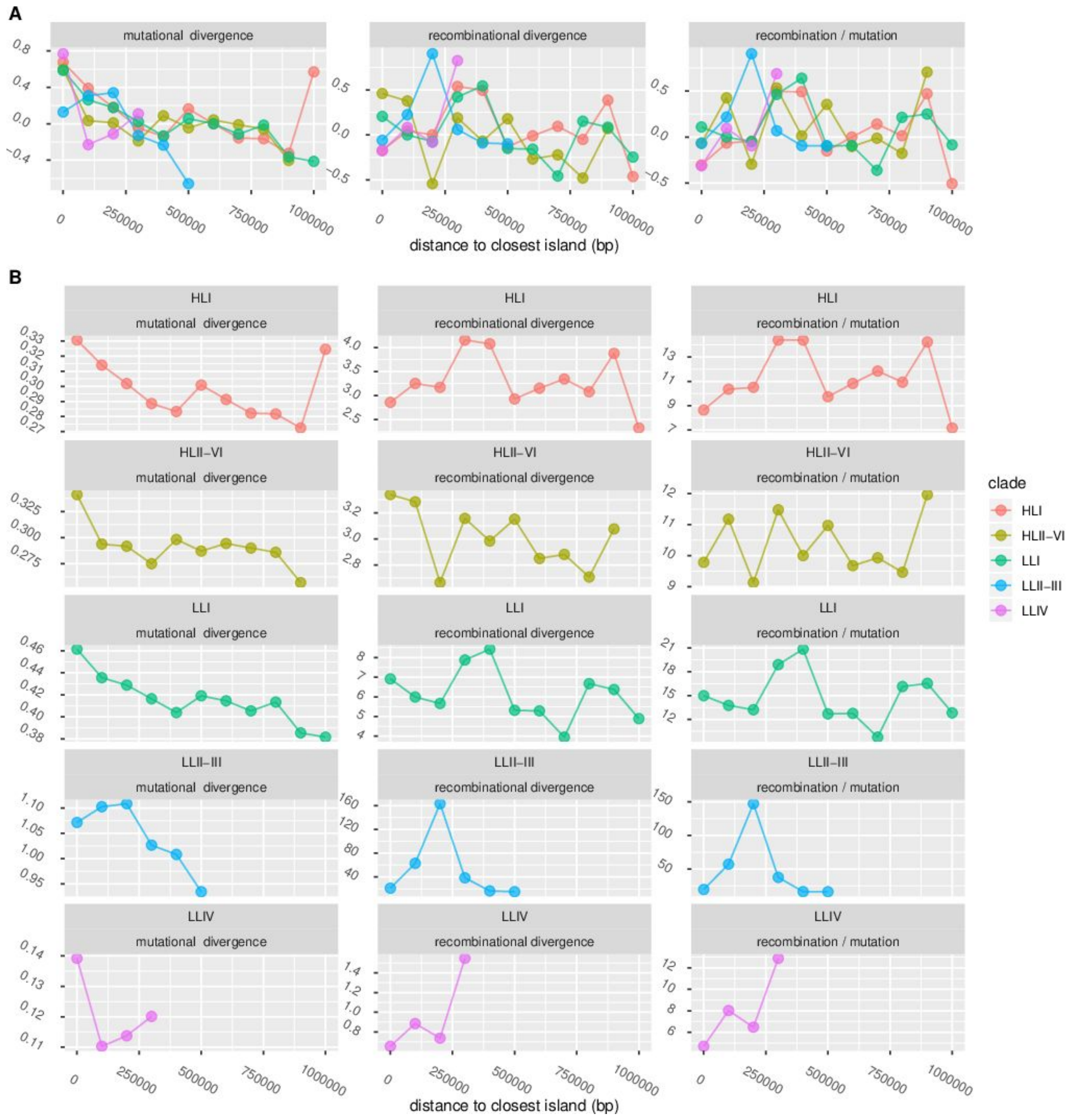


**Supplementary Figure 5 | Homologous recombination in island-flanking and non-island-flanking genomic regions.**
**A**) Across-clade comparison of three recombination-related parameters - mutational divergence, recombinational divergence and the ratio of the two - estimated using mcorr[28]. Values are derived from clade-specific estimates (B) and were mean-centred and rescaled for better visual comparison of trends across clades. Parameters were estimated on 99999 bp long partitions from a concatenated alignment of backbone genes, ordered by their proximity to genomic islands, from island-flanking genes (first partition) to genes furthest from islands (last partition). **B**) Absolute values of the same three recombination-related parameters for each individual clade. We note that in regions closest to genomic islands (first partition) the impact of recombination on divergence relative to mutation is among the lowest overall observed values, indicating that these locations are not hotspots of homologous recombination.

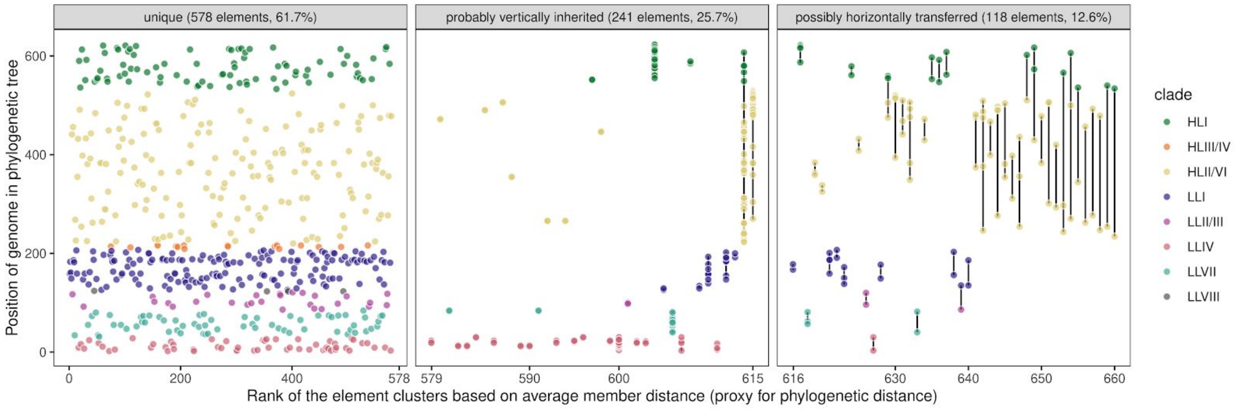


**Supplementary Figure 6 | Diversity and distribution of mobile elements across 623 *Prochlorococcus* genomes.**
Elements are clustered (coloured points linked by vertical black lines) based on significant pairwise similarity on the nucleotide level (at least 90% identity over 50% of the shorter element). The panels further group the mobile elements into three categories: elements found in only a single genome, elements found in some closely related genomes, likely representing vertical transmission, and elements found in multiple genomes with a noticeable difference in the phylogenetic position of their hosts, indicating possible horizontal transfer events.

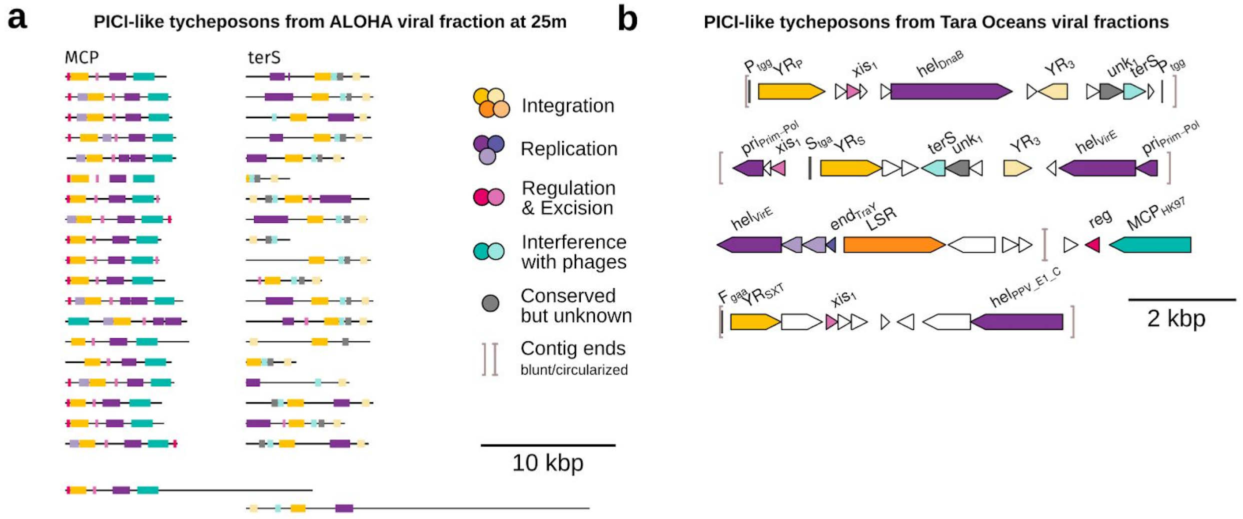


**Supplementary Figure 7 | Tycheposons in viral particles.**

**a**) Examples of nanopore reads from viral-fraction metagenomes from Station ALOHA appearing to be full-length tycheposons. Two reads also carry additional flanking material likely representing imprecise excision events and hinting at the ability of the tycheposons to promote the transfer of adjacent host material. **b**) PICI-like tycheposons in viral-fraction metagenomes from different Tara Ocean stations. Gene labels: tRNA genes and snippets are labeled with single-letter amino acid code and their anticodon, e.g. $S_{tga}$ = tRNA-$Serin_{TGA}$. YR = tyrosine recombinase; LSR = large serine recombinase; MCP = major capsid protein; terS = terminase small subunit; xis = excisionase; hel = helicase; top = topoisomerase; lig = ligase; reg = transcriptional regulator; pri = primase or primase/polymerase or primase/helicase; end = endonuclease; SSR = small serine recombinase; unk = conserved unknown. Some annotations are further labeled with their specific profile (e.g. $hel_{DnaB}$ or $hel_{VirE}$).

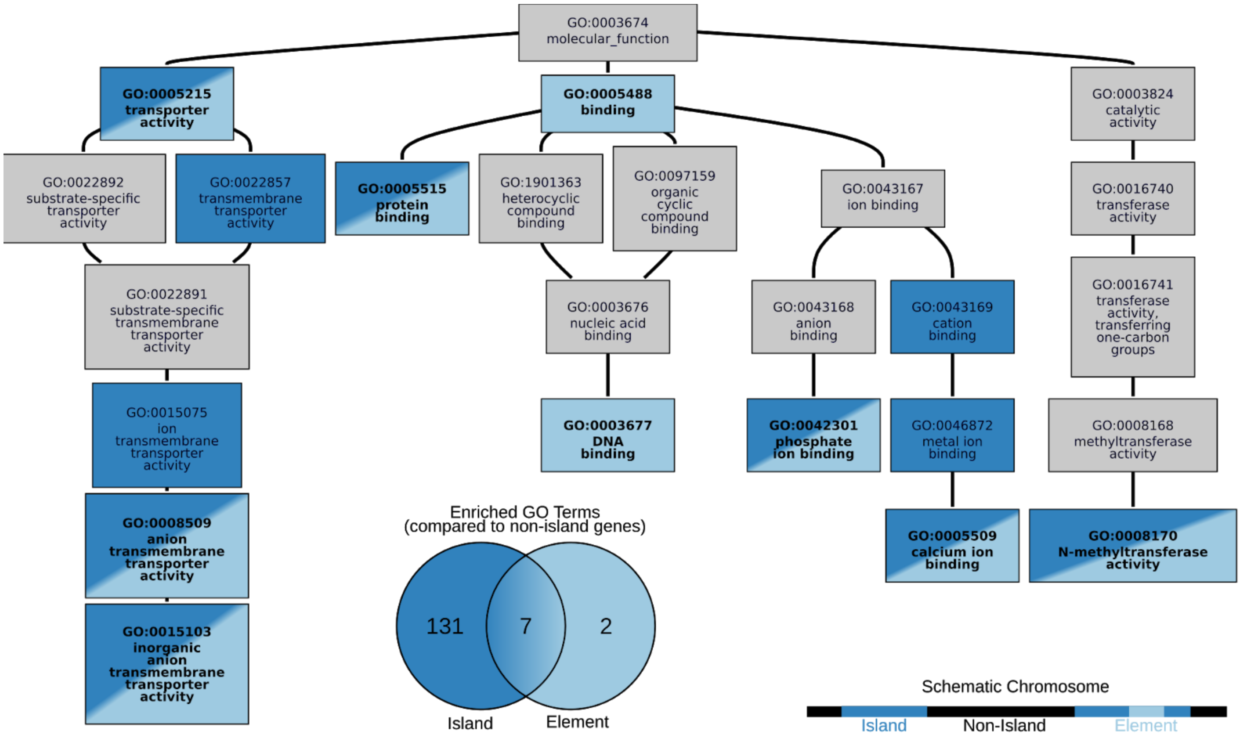


**Supplementary Figure 8 | Gene functions in elements and other island parts resemble each other.**

Subset of GO-terms in the Molecular Function sub-ontology. Enriched terms (compared to non-island genes) are coloured in shades of blue (see "Functional enrichment analysis" in methods for details). Light blue indicates enrichment of non-hallmark genes (genes not related to recombination and DNA-replication) on elements, while dark blue indicates that these terms are also enriched on genomic islands (excluding elements). There are another 127 terms in this sub-ontology that are enriched on genomic islands but not on elements (not shown).

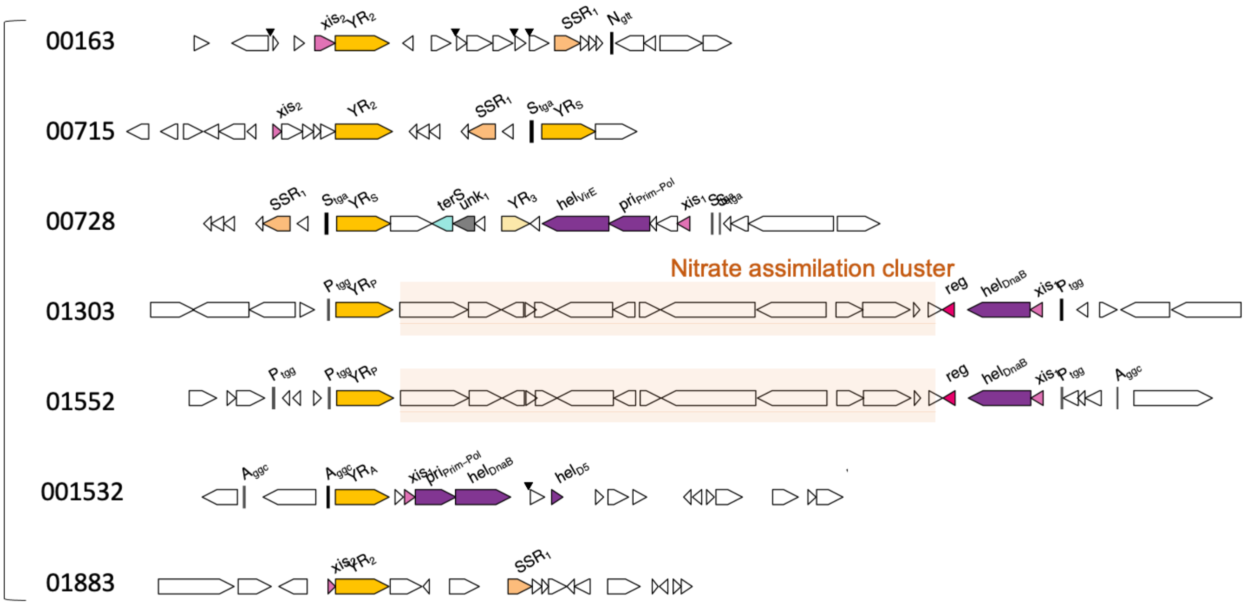


**Supplementary Figure 9 | *Prochlorococcus* MIT0604 integrative elements.**
Overview of the 7 different elements found in the *Prochlorococcus* MIT0604 reference strains. Elements 01303 and 01552 are the two identical copies of the tycheposon carrying the complete nitrate assimilation cluster[74] (orange box). The genes color code is identical to Figure 2.

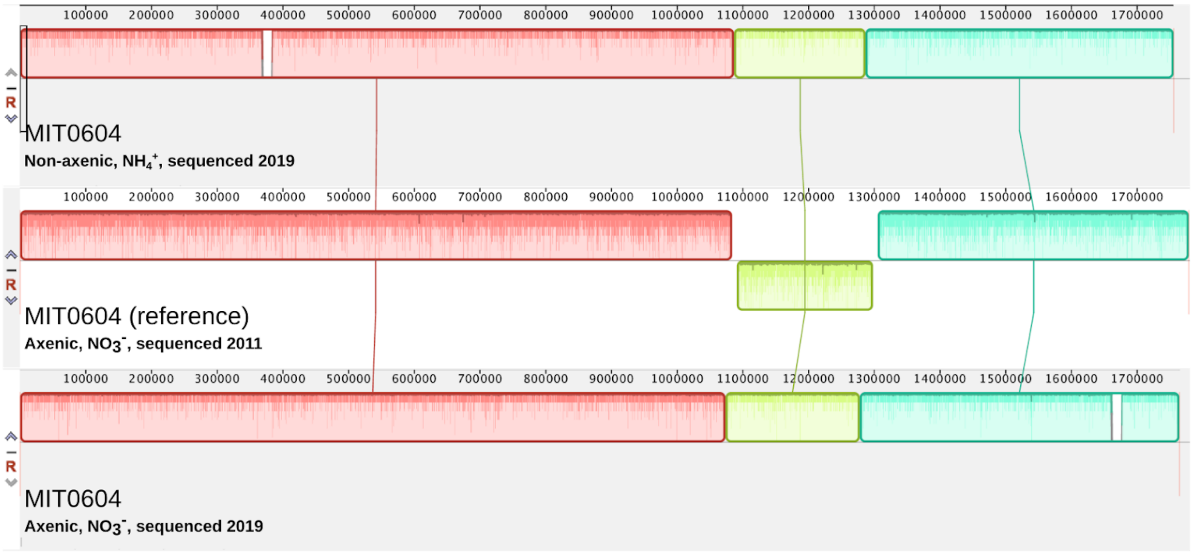


**Supplementary Figure 10 | Whole-genome alignment of *Prochlorococcus* MIT0604 strains showing element-related rearrangements**

Comparison of genomes of two cultures of *Prochlorococcus* MIT0604 at different points in time that were maintained in liquid culture for 10 years: *Prochlorococcus* MIT0604 (non-axenic, $NH_4^+$ ), the parent culture, was kept non-axenic, growing with ammonium as the sole nitrogen source, while *Prochlorococcus* MIT0604 (axenic, $NO_3^-$ ) was kept axenic, with nitrate as the sole nitrogen source. The parent culture was only sequenced in 2019, the culture growing in $NO_3^-$ in 2011 and 2019. The coloured bars represent syntenic regions of the genomes with the same arrangements. The yellow region showing an inversion in the reference is flanked by the integration sites of the duplicated *narB* cluster containing tycheposon unique to this strain[75]. The visible white gap in the third assembly at ~1.65 Mbp corresponds to an additional copy of the *narB* element that integrated at a tertiary location. The gap in the first assembly at 0.37 Mbp corresponds to another element lost in the culture grown on $NO_3^-$. All genomic rearrangements observed in these three genomes are, thus, tied to the presence and potentially also the activity of tycheposon elements.

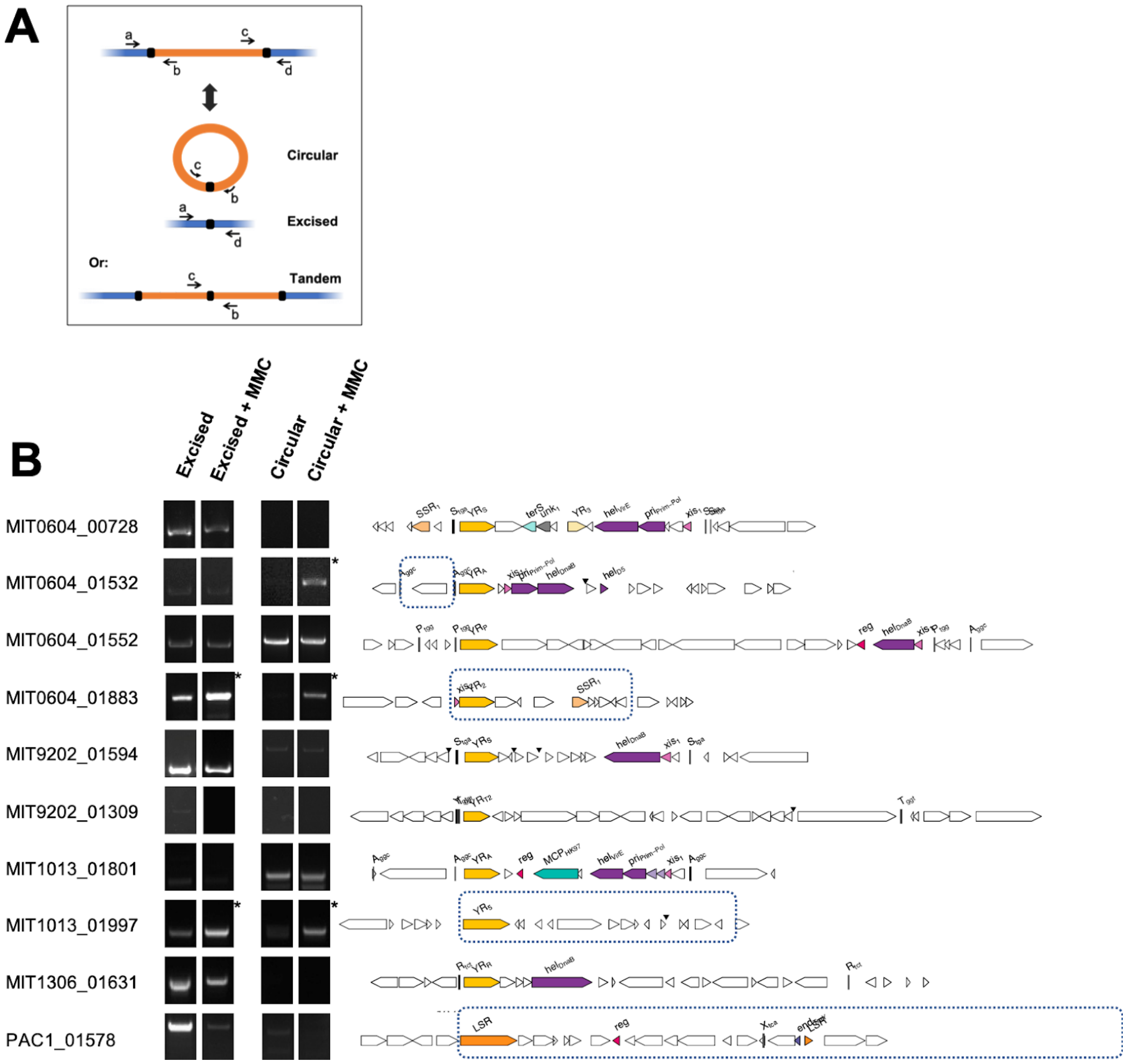


**Supplementary Figure 11 | Detection of integration and excision of elements in cultures.**

**a)** Cartoon of the PCR strategy to detect element excision, showing a, b, c, d primer design listed in supplementary table 11. **b)** PCR products run on an agarose gel. The gel exposure varies from one PCR amplicon to another but is identical for the same amplicon +/- mitomycin C. When the direct tRNA repeat borders of the element are not apparent, a blue dotted box indicates the predicted borders. Most elements show that the starting population is heterogeneous (excision or tandem repeats of elements amplify from a subpopulation of cells). Only a few elements show mobilisation by mitomycin C (indicated by a star), producing a likely circular intermediate. Interestingly, primers for the element MIT0604_01532 amplify a segment of DNA that does not contain the integrase, demonstrating the ability of elements to move segments of DNA *in trans*, as long as attachment sites are present.

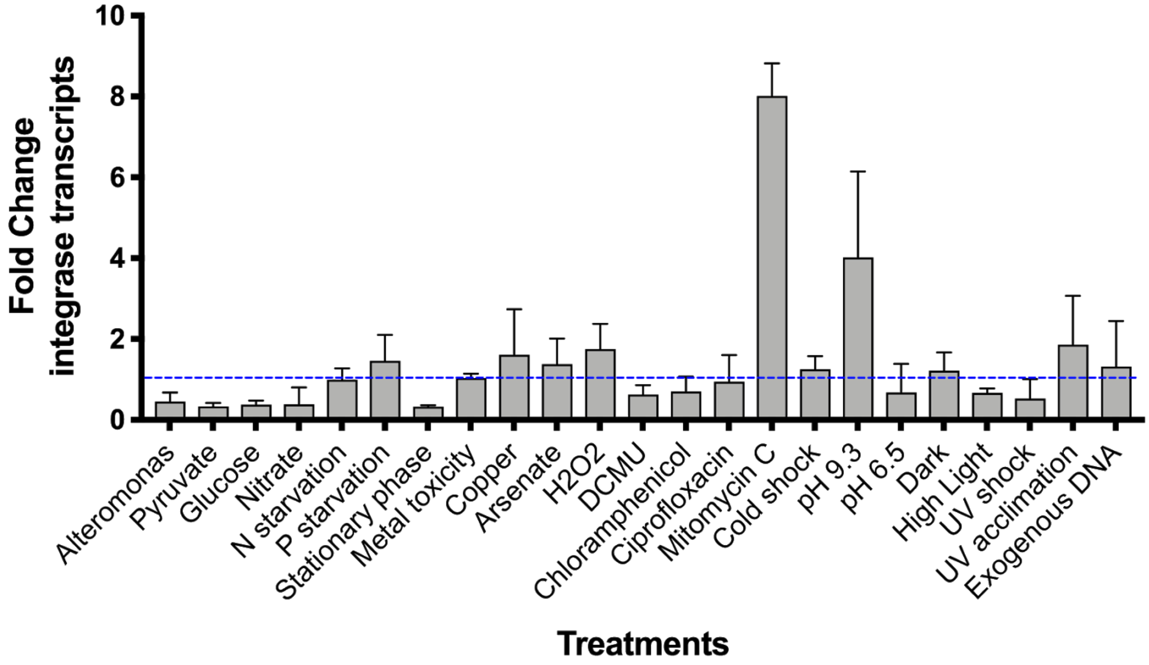


**Supplementary Figure 12 | Transcriptional response of integrase genes associated with cargo-carrying elements in *Prochlorococcus* MIT0604 in response to a wide range of shock treatments.**

Reverse-Transcription qPCR analysis of MIT0604_01303 integrase was performed on biological triplicates exposed to the treatment, compared to untreated controls. The dotted blue line indicates no change from treatment to control. See methods for details about treatments.

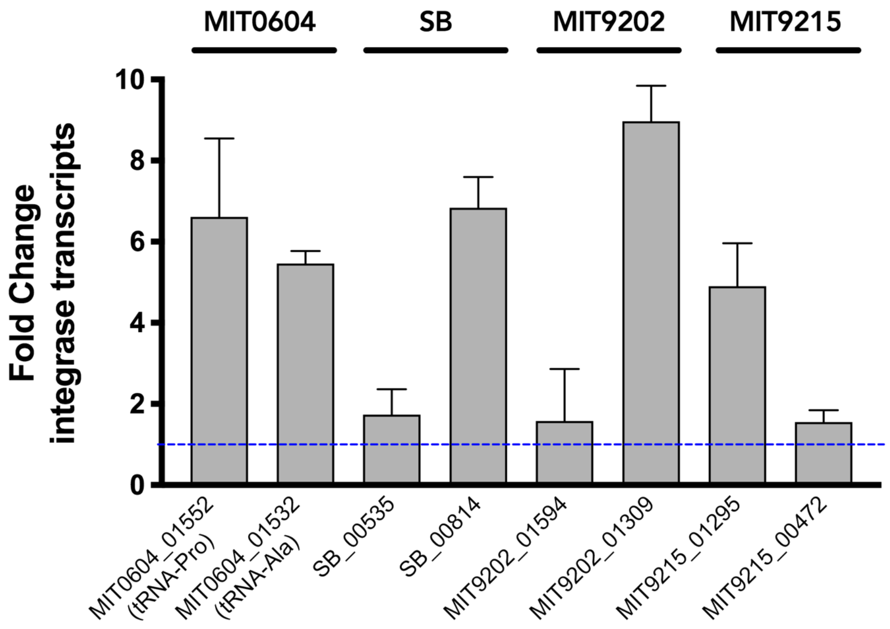


**Supplementary Figure 13 | Effect of mitomycin C treatment on various tycheposon integrases.**
Reverse-Transcription qPCR analysis of 8 integrase genes from 4 different *Prochlorococcus* strains. The dotted blue line indicates no change from treatment to control. Mitomycin C treatment was performed as mentioned in Supplementary Fig. 10. Five integrases, distantly related to each other, were upregulated in response to mitomycin C, suggesting that this type of DNA damage is a general induction cue for tycheposon elements.

### References


1. Parks DH, Imelfort M, Skennerton CT, Hugenholtz P, Tyson GW. CheckM: assessing the quality of microbial genomes recovered from isolates, single cells, and metagenomes. Genome Res. 2015;25: 1043–1055. doi:10.1101/gr.186072.114
2. Hackl T, Ankenbrand MJ. Supplementary code and data for "Novel integrative elements and genomic plasticity in ocean ecosystems." Zenodo. 2020. doi:10.5281/zenodo.4383240
3. Pachiadaki MG, Brown JM, Brown J, Bezuidt O, Berube PM, Biller SJ, et al. Charting the Complexity of the Marine Microbiome through Single-Cell Genomics. Cell. 2019;179: 1623–1635.e11. doi:10.1016/j.cell.2019.11.017
4. Biller SJ, Berube PM, Dooley K, Williams M, Satinsky BM, Hackl T, et al. Marine microbial metagenomes sampled across space and time. Scientific Data. 2018;5: 180176. doi:10.1038/sdata.2018.176
5. Biller SJ, Schubotz F, Roggensack SE, Thompson AW, Summons RE, Chisholm SW. Bacterial Vesicles in Marine Ecosystems. Science. 2014. pp. 183–186. doi:10.1126/science.1243457
6. Beaulaurier J, Luo E, Eppley JM, Uyl PD, Dai X, Burger A, et al. Assembly-free single-molecule sequencing recovers complete virus genomes from natural microbial communities. Genome Res. 2020;30: 437–446. doi:10.1101/gr.251686.119
7. Seemann T. Prokka: Rapid prokaryotic genome annotation. Bioinformatics. 2014;30: 2068–2069. doi:10.1093/bioinformatics/btu153
8. Ding W, Baumdicker F, Neher RA. panX: pan-genome analysis and exploration. Nucleic Acids Res. 2017. doi:10.1093/nar/gkx977
9. Eddy SR. Accelerated Profile HMM Searches. PLoS Comput Biol. 2011;7: e1002195. doi:10.1371/journal.pcbi.1002195
10. Haft DH, Loftus BJ, Richardson DL, Yang F, Eisen JA, Paulsen IT, et al. TIGRFAMs: a protein family resource for the functional identification of proteins. Nucleic Acids Res. 2001;29: 41–43. doi:10.1093/nar/29.1.41
11. Parks DH, Chuvochina M, Waite DW, Rinke C, Skarshewski A, Chaumeil P-A, et al. A standardized bacterial taxonomy based on genome phylogeny substantially revises the tree of life. Nat Biotechnol. 2018;36: 996–1004. doi:10.1038/nbt.4229
12. Shen W, Le S, Li Y, Hu F. SeqKit: A Cross-Platform and Ultrafast Toolkit for FASTA/Q File Manipulation. PLoS One. 2016;11: e0163962. doi:10.1371/journal.pone.0163962
13. Nakamura T, Yamada KD, Tomii K, Katoh K. Parallelization of MAFFT for large-scale multiple sequence alignments. Bioinformatics. 2018;34: 2490–2492. doi:10.1093/bioinformatics/bty121
14. Capella-Gutiérrez S, Silla-Martínez JM, Gabaldón T. trimAl: a tool for automated alignment trimming in large-scale phylogenetic analyses. Bioinformatics. 2009;25: 1972–1973. doi:10.1093/bioinformatics/btp348
15. Price MN, Dehal PS, Arkin AP. FastTree 2--approximately maximum-likelihood trees for large alignments. PLoS One. 2010;5: e9490. doi:10.1371/journal.pone.0009490
16. Revell LJ. phytools: an R package for phylogenetic comparative biology (and other things): phytools: R package. Methods Ecol Evol. 2012;3: 217–223. doi:10.1111/j.2041-210X.2011.00169.x
17. Yu G, Lam TT-Y, Zhu H, Guan Y. Two Methods for Mapping and Visualizing Associated Data on Phylogeny Using Ggtree. Mol Biol Evol. 2018;35: 3041–3043. doi:10.1093/molbev/msy194
18. Gao F, Zhang C-T. Ori-Finder: A web-based system for finding oriC s in unannotated bacterial genomes. BMC Bioinformatics. 2008;9: 79. doi:10.1186/1471-2105-9-79
19. Gao F, Luo H, Zhang C-T. DoriC 5.0: an updated database of oriC regions in both bacterial and archaeal genomes. Nucleic Acids Res. 2013;41: D90–3. doi:10.1093/nar/gks990
20. Tang H, Zhang X, Miao C, Zhang J, Ming R, Schnable JC, et al. ALLMAPS: robust scaffold ordering based on multiple maps. Genome Biol. 2015;16: 3. doi:10.1186/s13059-014-0573-1
21. Team RC, Others. R: A language and environment for statistical computing. 2013. Available: ftp://ftp.uvigo.es/CRAN/web/packages/dplR/vignettes/intro-dplR.pdf

22. Coleman ML, Sullivan MB, Martiny AC, Steglich C, Barry K, Delong EF, et al. Genomic islands and the ecology and evolution of Prochlorococcus. Science. 2006;311: 1768–1770. doi:10.1126/science.1122050
23. Dufresne A, Ostrowski M, Scanlan DJ, Garczarek L, Mazard S, Palenik BP, et al. Unraveling the genomic mosaic of a ubiquitous genus of marine cyanobacteria. Genome Biol. 2008;9: R90. doi:10.1186/gb-2008-9-5-r90
24. Avrani S, Wurtzel O, Sharon I, Sorek R, Lindell D. Genomic island variability facilitates Prochlorococcus-virus coexistence. Nature. 2011;474: 604–608. doi:10.1038/nature10172
25. Himmelmann L. HMM: HMM-Hidden Markov Models. R package version 1.0. 2016.
26. Rodriguez-Valera F, Martin-Cuadrado A-B, López-Pérez M. Flexible genomic islands as drivers of genome evolution. Curr Opin Microbiol. 2016;31: 154–160. doi:10.1016/j.mib.2016.03.014
27. Oliveira PH, Touchon M, Cury J, Rocha EPC. The chromosomal organization of horizontal gene transfer in bacteria. Nat Commun. 2017;8: 841. doi:10.1038/s41467-017-00808-w
28. Lin M, Kussell E. Inferring bacterial recombination rates from large-scale sequencing datasets. Nat Methods. 2019;16: 199–204. doi:10.1038/s41592-018-0293-7
29. Larsson A. AliView: a fast and lightweight alignment viewer and editor for large datasets. Bioinformatics. 2014;30: 3276–3278. doi:10.1093/bioinformatics/btu531
30. Rambaut A. FigTree v1. 4. 2012.
31. El-Gebali S, Mistry J, Bateman A, Eddy SR, Luciani A, Potter SC, et al. The Pfam protein families database in 2019. Nucleic Acids Res. 2019;47: D427–D432. doi:10.1093/nar/gky995
32. Hiramatsu K, Ito T, Tsubakishita S, Sasaki T, Takeuchi F, Morimoto Y, et al. Genomic Basis for Methicillin Resistance in Staphylococcus aureus. Infect Chemother. 2013;45: 117–136. doi:10.3947/ic.2013.45.2.117
33. Fillol-Salom A, Martínez-Rubio R, Abdulrahman RF, Chen J, Davies R, Penadés JR. Phage-inducible chromosomal islands are ubiquitous within the bacterial universe. ISME J. 2018;12: 2114–2128. doi:10.1038/s41396-018-0156-3
34. Martínez-Rubio R, Quiles-Puchalt N, Martí M, Humphrey S, Ram G, Smyth D, et al. Phage-inducible islands in the Gram-positive cocci. ISME J. 2017;11: 1029–1042. doi:10.1038/ismej.2016.163
35. O'Hara BJ, Barth ZK, McKitterick AC, Seed KD. A highly specific phage defense system is a conserved feature of the Vibrio cholerae mobilome. PLoS Genet. 2017;13: e1006838. doi:10.1371/journal.pgen.1006838
36. Hyatt D, Chen G-L, Locascio PF, Land ML, Larimer FW, Hauser LJ. Prodigal: prokaryotic gene recognition and translation initiation site identification. BMC Bioinformatics. 2010;11: 119. doi:10.1186/1471-2105-11-119
37. Laslett D, Canback B. ARAGORN, a program to detect tRNA genes and tmRNA genes in nucleotide sequences. Nucleic Acids Res. 2004;32: 11–16. doi:10.1093/nar/gkh152
38. Roux S, Enault F, Hurwitz BL, Sullivan MB. VirSorter: mining viral signal from microbial genomic data. PeerJ. 2015;3: e985. doi:10.7717/peerj.985
39. Altschul SF, Gish W, Miller W, Myers EW, Lipman DJ. Basic local alignment search tool. J Mol Biol. 1990;215: 403–410. doi:10.1016/S0022-2836(05)80360-2
40. Buchfink B, Xie C, Huson DH. Fast and sensitive protein alignment using DIAMOND. Nat Methods. 2015;12: 59–60. doi:10.1038/nmeth.3176
41. Alexa A, Rahnenfuhrer J. topGO: enrichment analysis for gene ontology. R package version. 2010;2: 2010. Available: http://bioconductor.uib.no/2.7/bioc/html/topGO.html
42. Liao Y, Wang J, Jaehnig EJ, Shi Z, Zhang B. WebGestalt 2019: gene set analysis toolkit with revamped UIs and APIs. Nucleic Acids Research. 2019. pp. W199–W205. doi:10.1093/nar/gkz401
43. Smyshlyaev G, Barabas O, Bateman A. Sequence analysis allows functional annotation of tyrosine recombinases in prokaryotic genomes. bioRxiv. 2019. p. 542381. doi:10.1101/542381
44. Letunic I, Bork P. Interactive Tree Of Life (iTOL) v4: recent updates and new developments. Nucleic Acids Res. 2019;47: W256–W259. doi:10.1093/nar/gkz239
45. Woodcroft BJ, Boyd JA, Tyson GW. OrfM: a fast open reading frame predictor for metagenomic data. Bioinformatics. 2016;32: 2702–2703. doi:10.1093/bioinformatics/btw241
46. Robinson MD, McCarthy DJ, Smyth GK. edgeR: a Bioconductor package for differential expression analysis of digital gene expression data. Bioinformatics. 2010;26: 139–140. doi:10.1093/bioinformatics/btp616
47. Bushnell BJR. BBMap short read aligner, and other bioinformatic tools. 2014. Available: http://sourceforge.net/projects/bbmap/
48. Wickham H. ggplot2. WIREs Comp Stat. 2011;3: 180–185. doi:10.1002/wics.147
49. Kearse M, Moir R, Wilson A, Stones-Havas S, Cheung M, Sturrock S, et al. Geneious Basic: an integrated and

extendable desktop software platform for the organization and analysis of sequence data. Bioinformatics. 2012;28: 1647–1649. doi:10.1093/bioinformatics/bts199
50. Darling AE, Mau B, Perna NT. Progressivemauve: Multiple genome alignment with gene gain, loss and rearrangement. PLoS One. 2010;5. doi:10.1371/journal.pone.0011147
51. Li H. Minimap2: pairwise alignment for nucleotide sequences. Bioinformatics. 2018;34: 3094–3100. doi:10.1093/bioinformatics/bty191
52. Nattestad M, Chin C-S, Schatz MC. Ribbon: Visualizing complex genome alignments and structural variation. bioRxiv. 2016. p. 082123. doi:10.1101/082123
53. Quick J. Ultra-long read sequencing protocol for RAD004 v3 (protocols.io.mrxc57n). 2018. doi:10.17504/protocols.io.mrxc57n
54. Wilson K. Preparation of genomic DNA from bacteria. Curr Protoc Mol Biol. 2001;Chapter 2: Unit 2.4. doi:10.1002/0471142727.mb0204s56
55. Chen I-MA, Markowitz VM, Chu K, Palaniappan K, Szeto E, Pillay M, et al. IMG/M: integrated genome and metagenome comparative data analysis system. Nucleic Acids Res. 2016. doi:10.1093/nar/gkw929
56. Markowitz VM, Chen I-MA, Palaniappan K, Chu K, Szeto E, Pillay M, et al. IMG 4 version of the integrated microbial genomes comparative analysis system. Nucleic Acids Research. 2014. pp. D560–D567. doi:10.1093/nar/gkt963
57. Berube PM, Biller SJ, Hackl T, Hogle SL, Satinsky BM, Becker JW, et al. Single cell genomes of Prochlorococcus, Synechococcus, and sympatric microbes from diverse marine environments. Scientific Data. 2018;5: 180154. doi:10.1038/sdata.2018.154
58. Moore LR, Coe A, Zinser ER, Saito MA, Sullivan MB, Lindell D, et al. Culturing the marine cyanobacterium Prochlorococcus. Limnol Oceanogr Methods. 2007;5: 353–362. Available: https://aslopubs.onlinelibrary.wiley.com/doi/abs/10.4319/lom.2007.5.353
59. Biller SJ, Berube PM, Berta-Thompson JW, Kelly L, Roggensack SE, Awad L, et al. Genomes of diverse isolates of the marine cyanobacterium Prochlorococcus. Scientific Data. 2014;1: 140034. Available: http://www.nature.com/articles/sdata201434
60. Thompson AW, Huang K, Saito MA, Chisholm SW. Transcriptome response of high- and low-light-adapted Prochlorococcus strains to changing iron availability. ISME J. 2011;5: 1580–1594. doi:10.1038/ismej.2011.49
61. Coleman ML, Sullivan MB, Martiny AC, Steglich C, Barry K, DeLong EF, et al. Genomic islands and the ecology and evolution of Prochlorococcus. Science. 2006;311: 1768–1770. Available: http://science.sciencemag.org/content/311/5768/1768.abstract
62. Kettler GC, Martiny AC, Huang K, Zucker J, Coleman ML, Rodrigue S, et al. Patterns and implications of gene gain and loss in the evolution of Prochlorococcus. PLoS Genet. 2007;3: e231. Available: http://journals.plos.org/plosgenetics/article?id=10.1371/journal.pgen.0030231
63. Shimada A, Kanai S, Maruyama T. Partial sequence of ribulose-1,5-bisphosphate carboxylase/oxygenase and the phylogeny of Prochloron and Prochlorococcus (Prochlorales). J Mol Evol. 1995;40: 671–677. doi:10.1007/bf00160516
64. Penno S, Campbell L, Hess WR. PRESENCE OF PHYCOERYTHRIN IN TWO STRAINS OF PROCHLOROCOCCUS (CYANOBACTERIA) ISOLATED FROM THE SUBTROPICAL NORTH PACIFIC OCEAN. J Phycol. 2000;36: 723–729. doi:10.1046/j.1529-8817.2000.99203.x
65. Cubillos-Ruiz A, Berta-Thompson JW, Becker JW, van der Donk WA, Chisholm SW. Evolutionary radiation of lanthipeptides in marine cyanobacteria. Proc Natl Acad Sci U S A. 2017;114: E5424–E5433. Available: http://www.pnas.org/lookup/doi/10.1073/pnas.1700990114
66. Pfaffl MW. A new mathematical model for relative quantification in real-time RT-PCR. Nucleic Acids Res. 2001;29: e45. Available: http://eutils.ncbi.nlm.nih.gov/entrez/eutils/elink.fcgi?dbfrom=pubmed&id=11328886&retmode=ref&cmd=prlinks
67. Laurenceau R, Bliem C, Osburne MS, Becker JW, Biller SJ, Cubillos-Ruiz A, et al. Toward a genetic system in the marine cyanobacterium Prochlorococcus. bioRxiv. 2019. p. 820027. doi:10.1101/820027
68. Osburne MS, Holmbeck BM, Frias-Lopez J, Steen R, Huang K, Kelly L, et al. UV hyper-resistance in Prochlorococcus MED4 results from a single base pair deletion just upstream of an operon encoding nudix hydrolase and photolyase. Environ Microbiol. 2010;12: 1978–1988. Available: http://doi.wiley.com/10.1111/j.1462-2920.2010.02203.x
69. Kolowrat C, Partensky F, Mella-Flores D, Le Corguillé G, Boutte C, Blot N, et al. Ultraviolet stress delays

chromosome replication in light/dark synchronized cells of the marine cyanobacterium Prochlorococcus marinus PCC9511. BMC Microbiol. 2010;10: 204. Available: http://www.biomedcentral.com/1471-2180/10/204
70. Bushnell B. BBTools software package. URL http://sourceforge net/projects/bbmap. 2014.
71. Li H, Durbin R. Fast and accurate short read alignment with Burrows-Wheeler transform. Bioinformatics. 2009. pp. 1754–1760. doi:10.1093/bioinformatics/btp324
72. Anders S, Pyl PT, Huber W. HTSeq—a Python framework to work with high-throughput sequencing data. Bioinformatics. 2015;31: 166–169. doi:10.1093/bioinformatics/btu638
73. Love MI, Huber W, Anders S. Moderated estimation of fold change and dispersion for RNA-seq data with DESeq2. Genome Biol. 2014;15: 550. doi:10.1186/s13059-014-0550-8
74. Berube PM, Biller SJ, Kent AG, Berta-Thompson JW, Kelly L, Roggensack SE, et al. Physiology and evolution of nitrogen acquisition in Prochlorococcus. 2014;9: 1195–1207. doi:10.1038/ismej.2014.211
75. Berube PM, Coe A, Roggensack SE, Chisholm SW. Temporal dynamics of Prochlorococcuscells with the potential for nitrate assimilation in the subtropical Atlantic and Pacific oceans. Limnol Oceanogr. 2015. Available: http://doi.wiley.com/10.1002/lno.10226
