## Supplement-2 for "Novel integrative elements and genomic plasticity in ocean ecosystems"

### YR_IRVE−P ( 261 ) 1

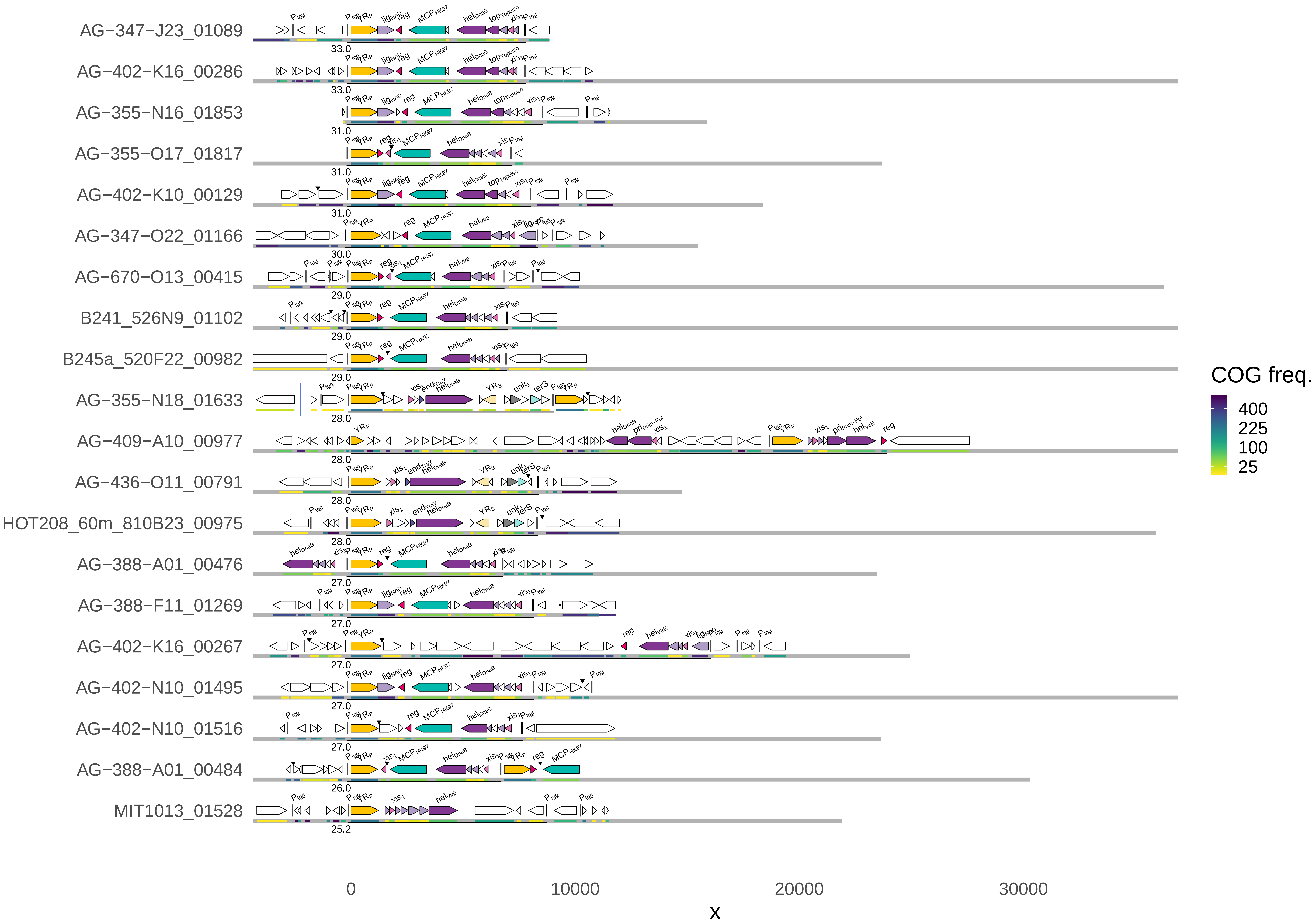

YR_IRVE−P ( 261 ) 2

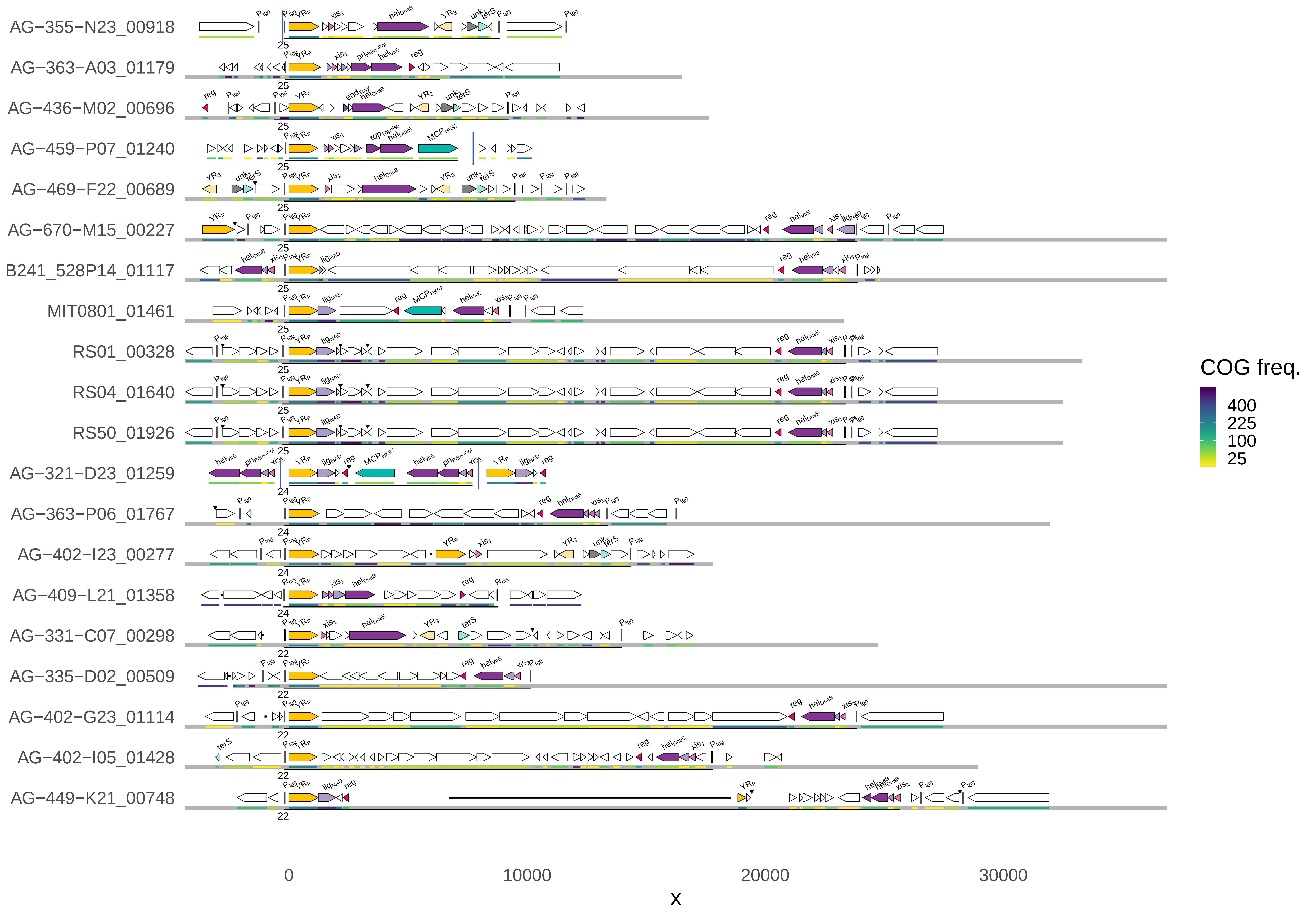

### YR_IRVE−P ( 261 ) 3

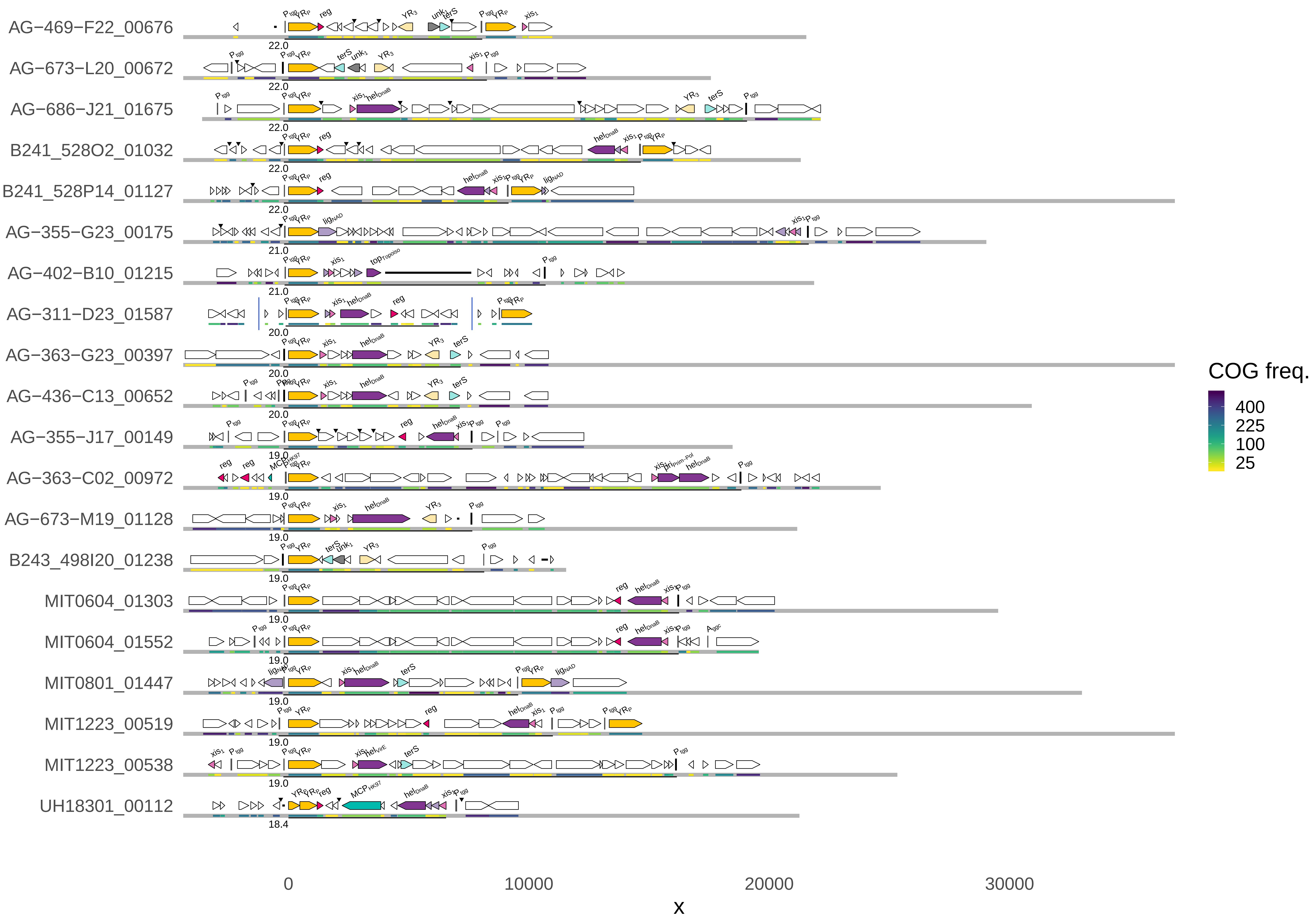

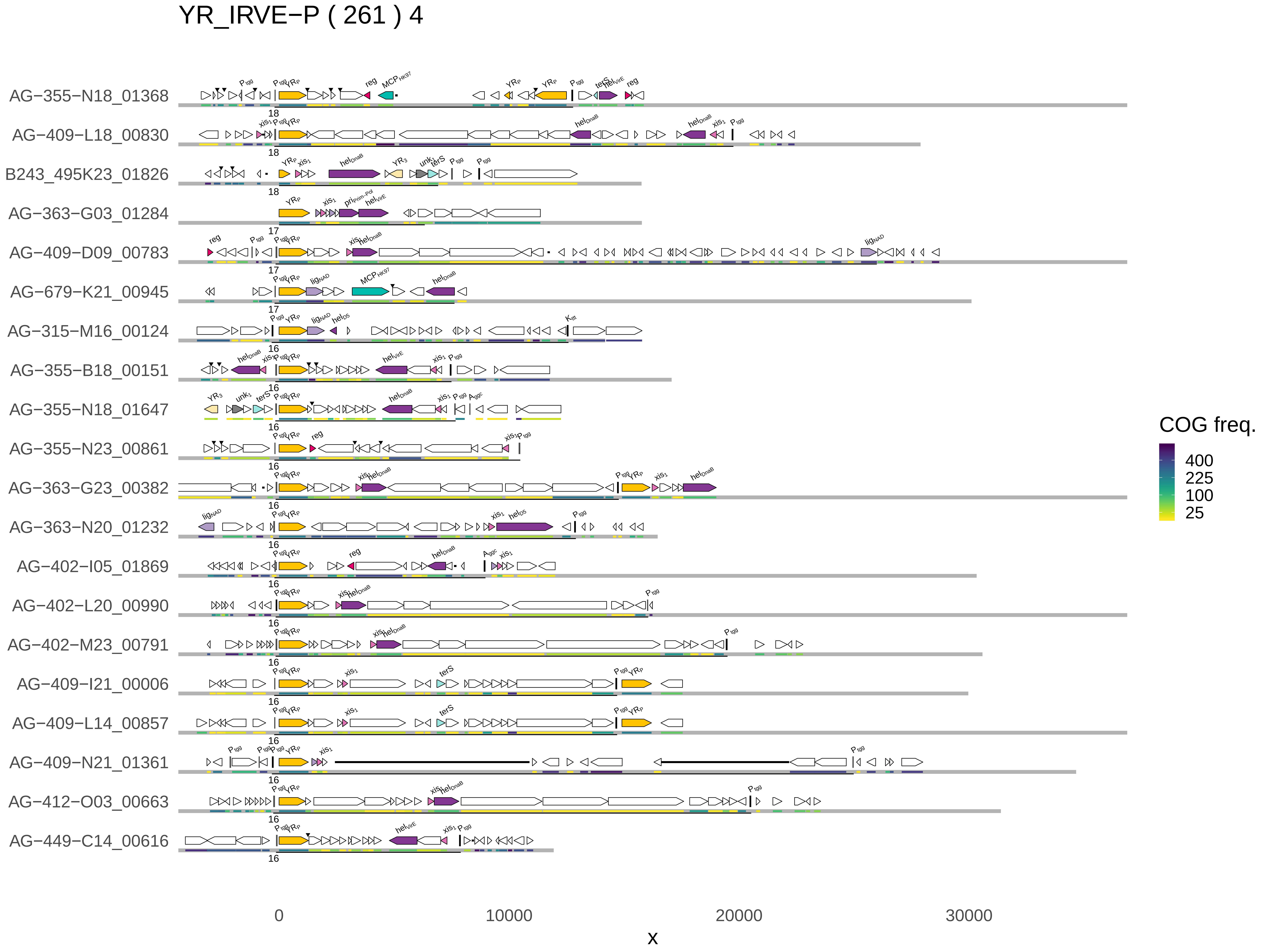

YR_IRVE−P ( 261 ) 4
AG−355−N18_01368
AG−409−L18_00830
B243_495K23_01826
AG−363−G03_01284
AG−409−D09_00783
AG−679−K21_00945
AG−315−M16_00124
AG−355−B18_00151
AG−355−N18_01647
AG−355−N23_00861
AG−363−G23_00382
AG−363−N20_01232
AG−402−I05_01869
AG−402−L20_00990
AG−402−M23_00791
AG−409−I21_00006
AG−409−L14_00857
AG−409−N21_01361
AG−412−O03_00663
AG−449−C14_00616
0
10000
20000
30000
x
COG freq.
400
225
100
25

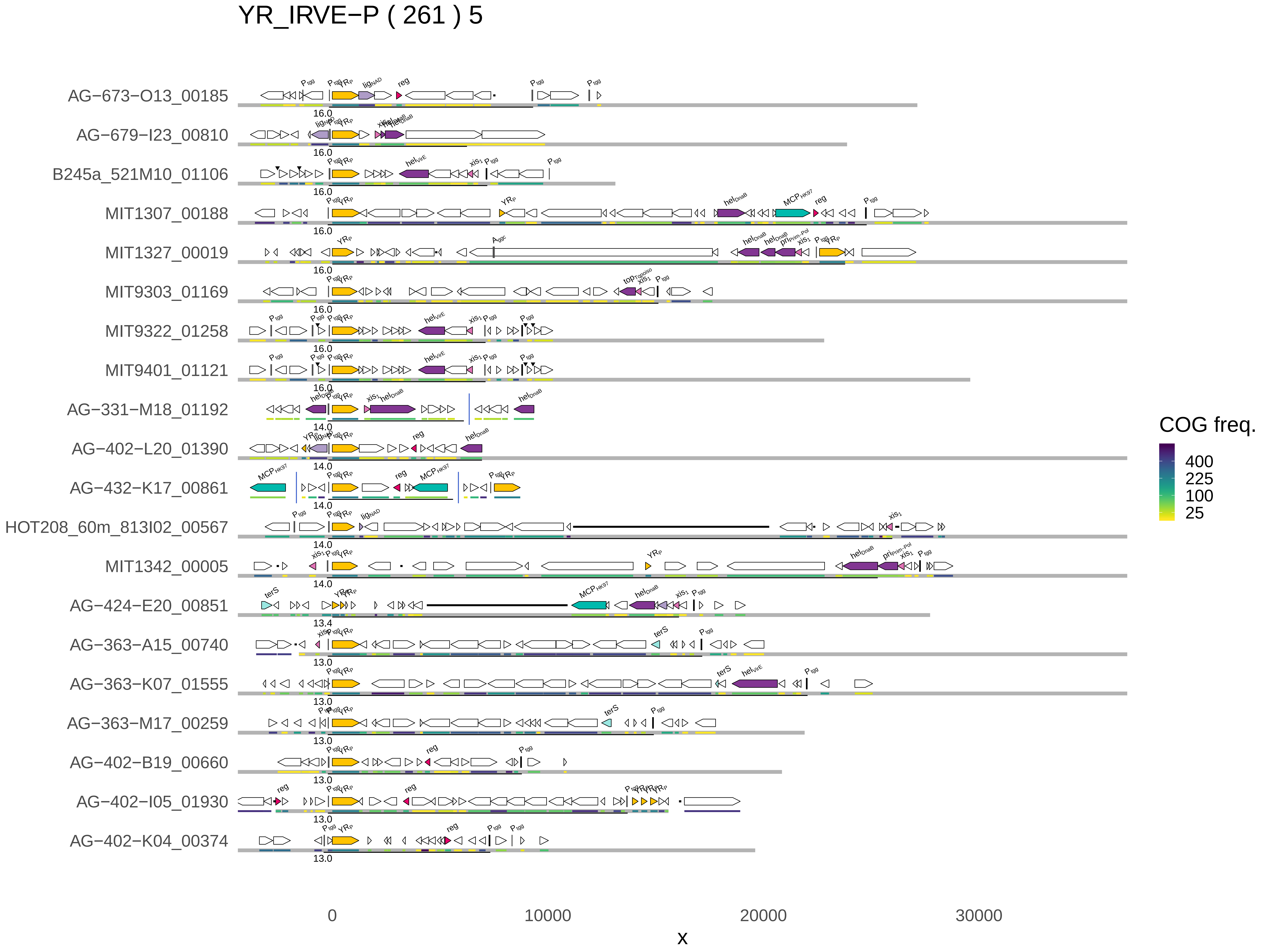

YR_IRVE−P ( 261 ) 5
AG−673−O13_00185
AG−679−I23_00810
B245a_521M10_01106
MIT1307_00188
MIT1327_00019
MIT9303_01169
MIT9322_01258
MIT9401_01121
AG−331−M18_01192
AG−402−L20_01390
AG−432−K17_00861
HOT208_60m_813I02_00567
MIT1342_00005
AG−424−E20_00851
AG−363−A15_00740
AG−363−K07_01555
AG−363−M17_00259
AG−402−B19_00660
AG−402−I05_01930
AG−402−K04_00374
0
10000
20000
30000
x
COG freq.
400
225
100
25

### YR_IRVE−P ( 261 ) 6

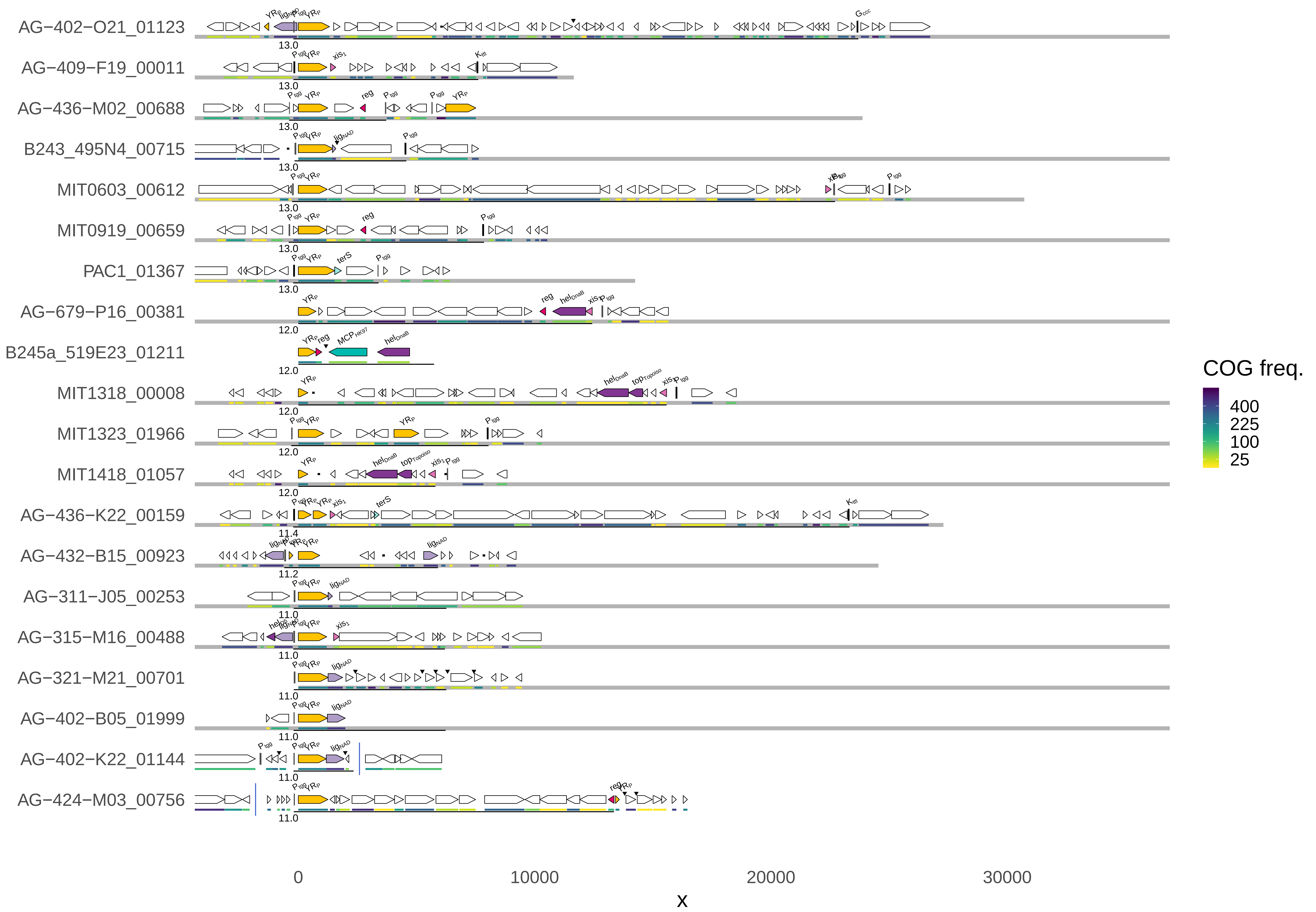

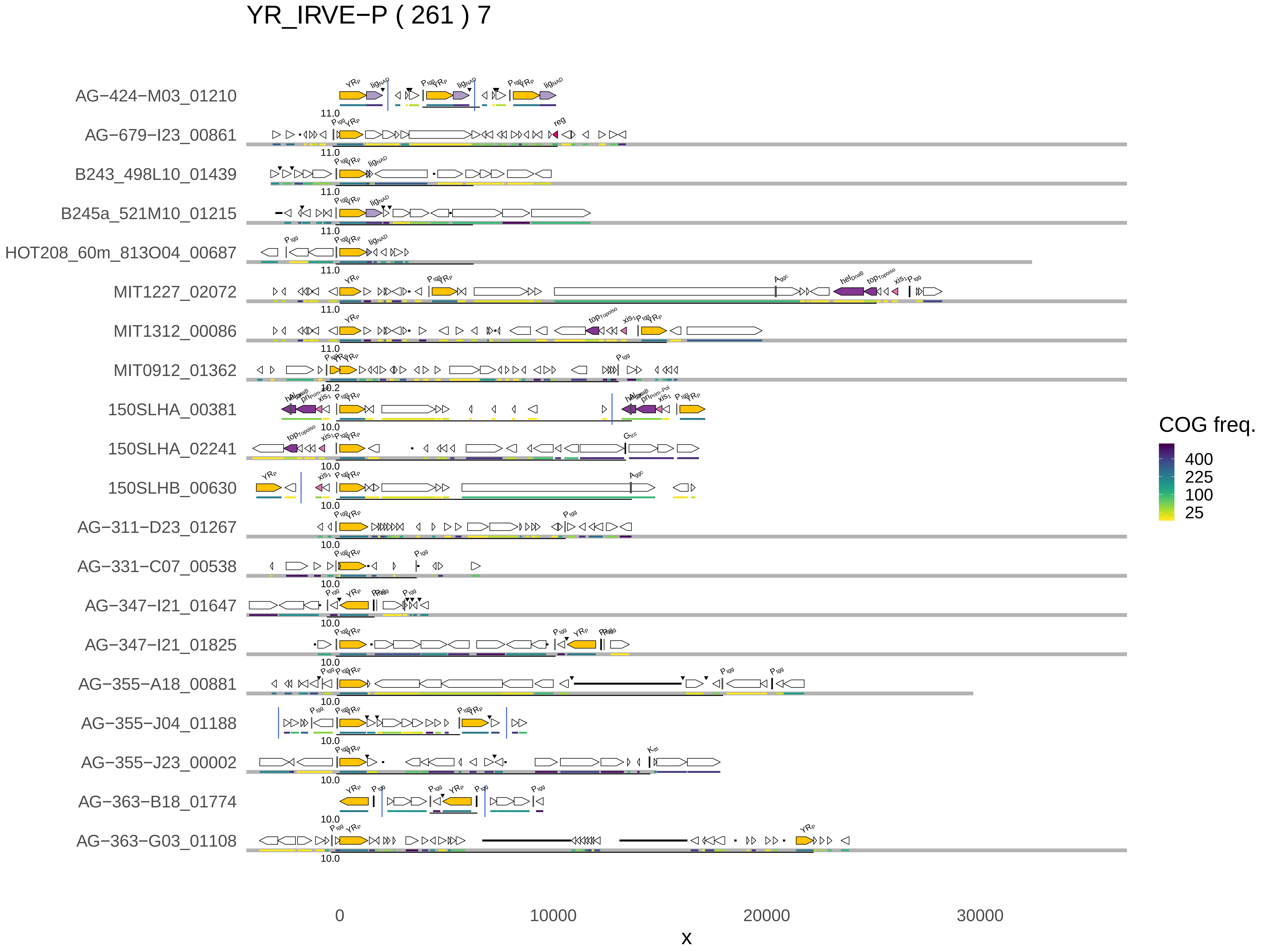
YR_IRVE−P ( 261 ) 7
AG−424−M03_01210
AG−679−I23_00861
B243_498L10_01439
B245a_521M10_01215
HOT208_60m_813O04_00687
MIT1227_02072
MIT1312_00086
MIT0912_01362
150SLHA_00381
150SLHA_02241
150SLHB_00630
AG−311−D23_01267
AG−331−C07_00538
AG−347−I21_01647
AG−347−I21_01825
AG−355−A18_00881
AG−355−J04_01188
AG−355−J23_00002
AG−363−B18_01774
AG−363−G03_01108
0
10000
20000
30000
x
COG freq.
400
225
100
25

YR_IRVE−P ( 261 ) 8

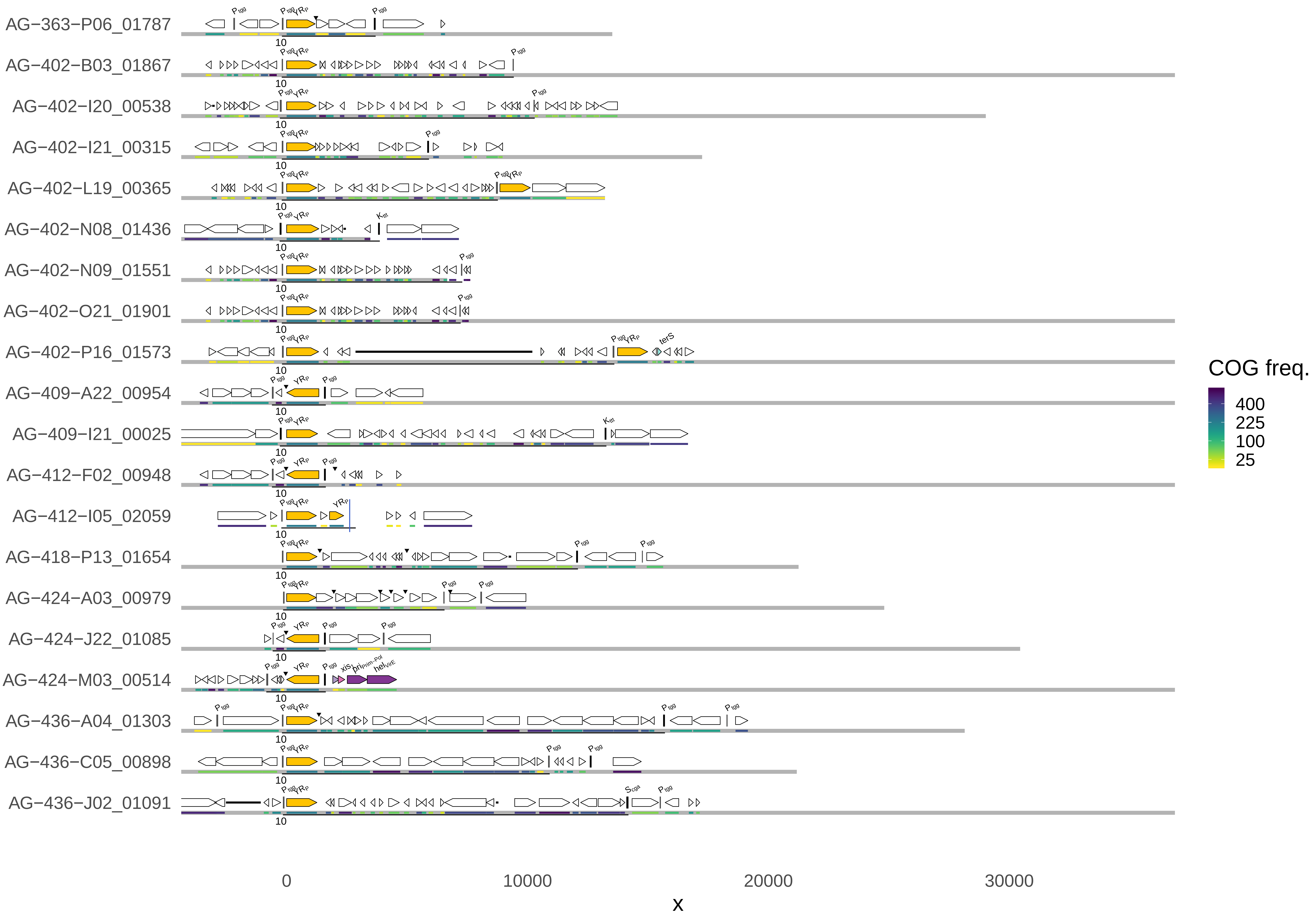

YR_IRVE−P ( 261 ) 9

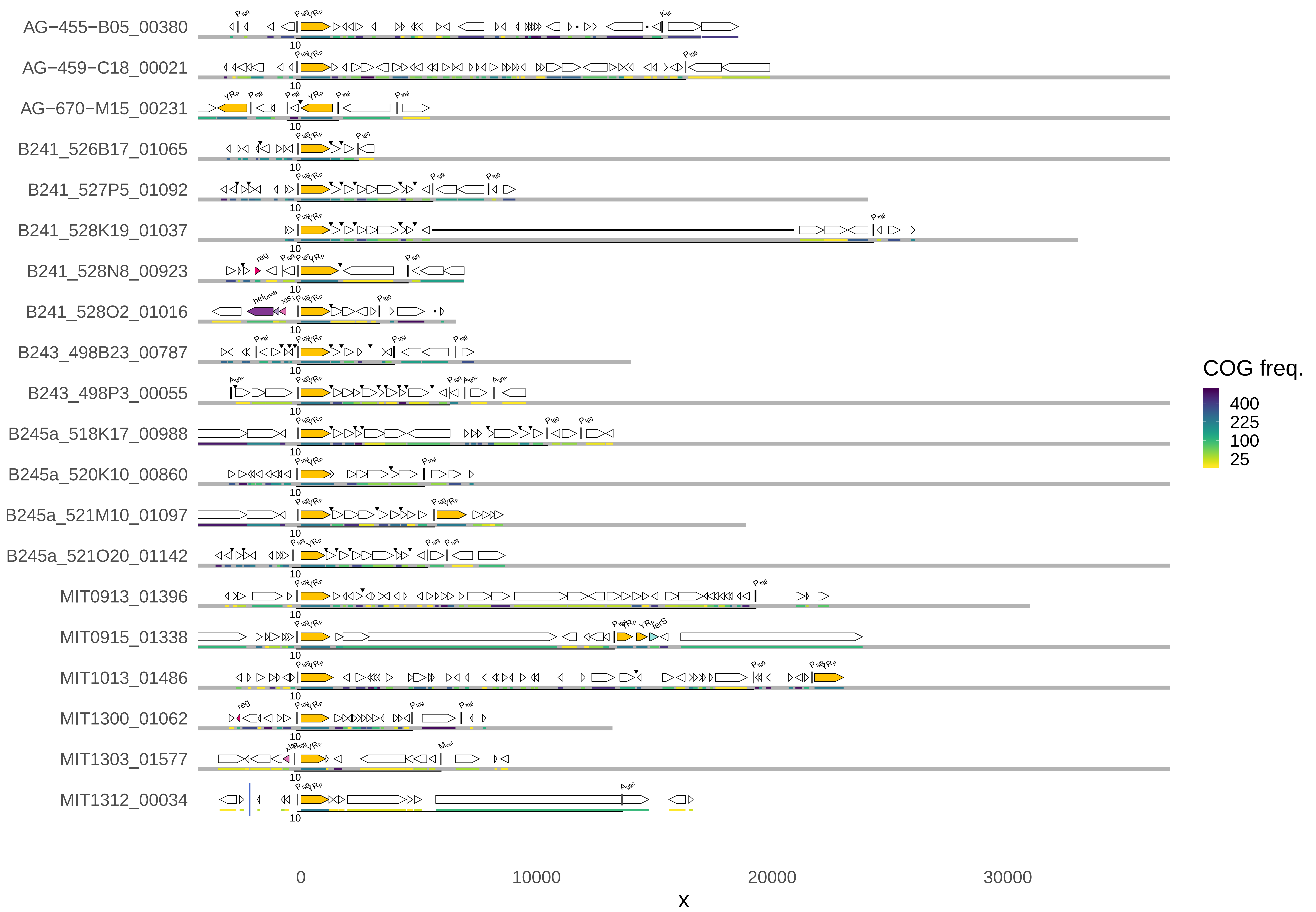

YR_IRVE−P ( 261 ) 10

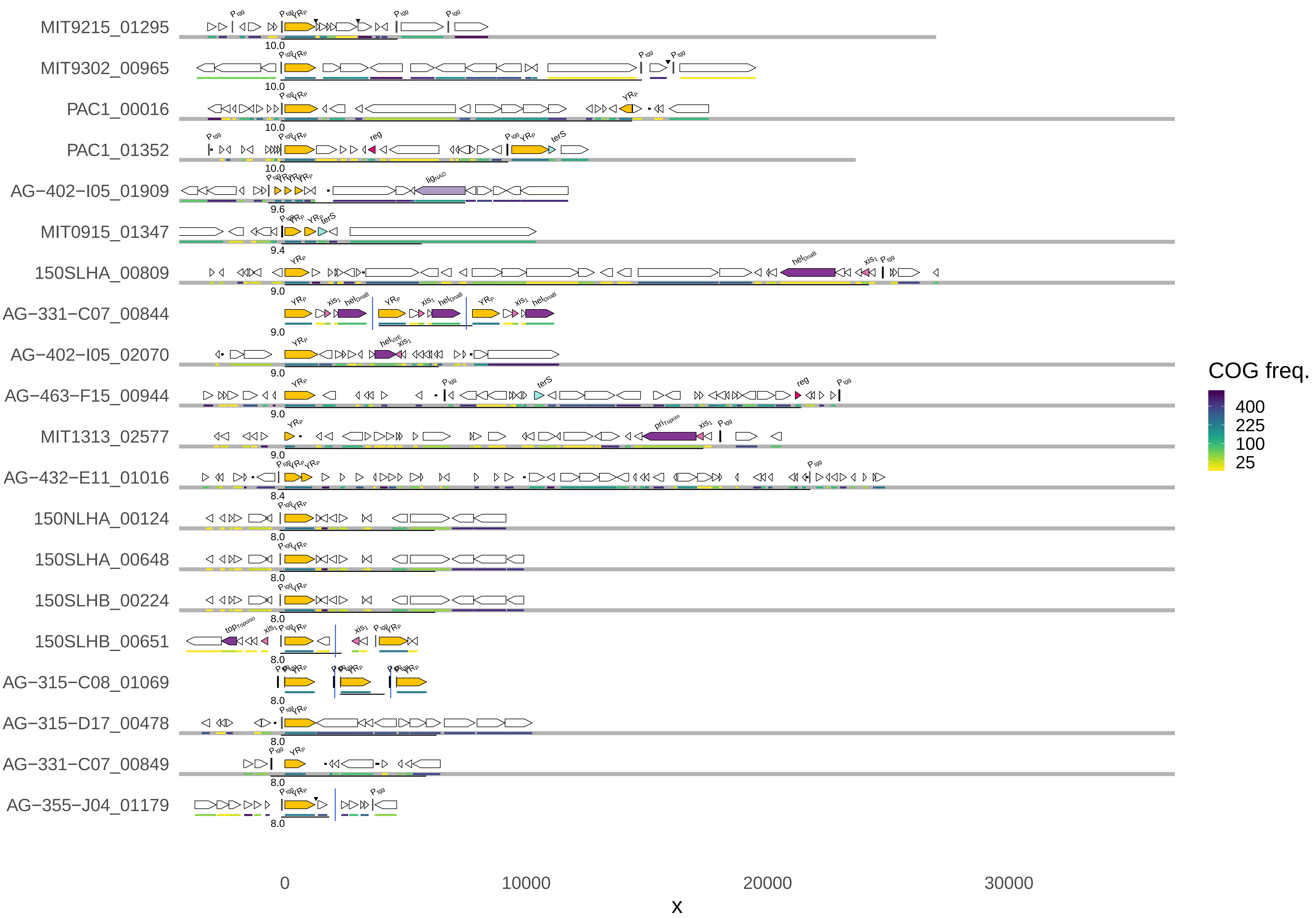

YR_IRVE−P ( 261 ) 11

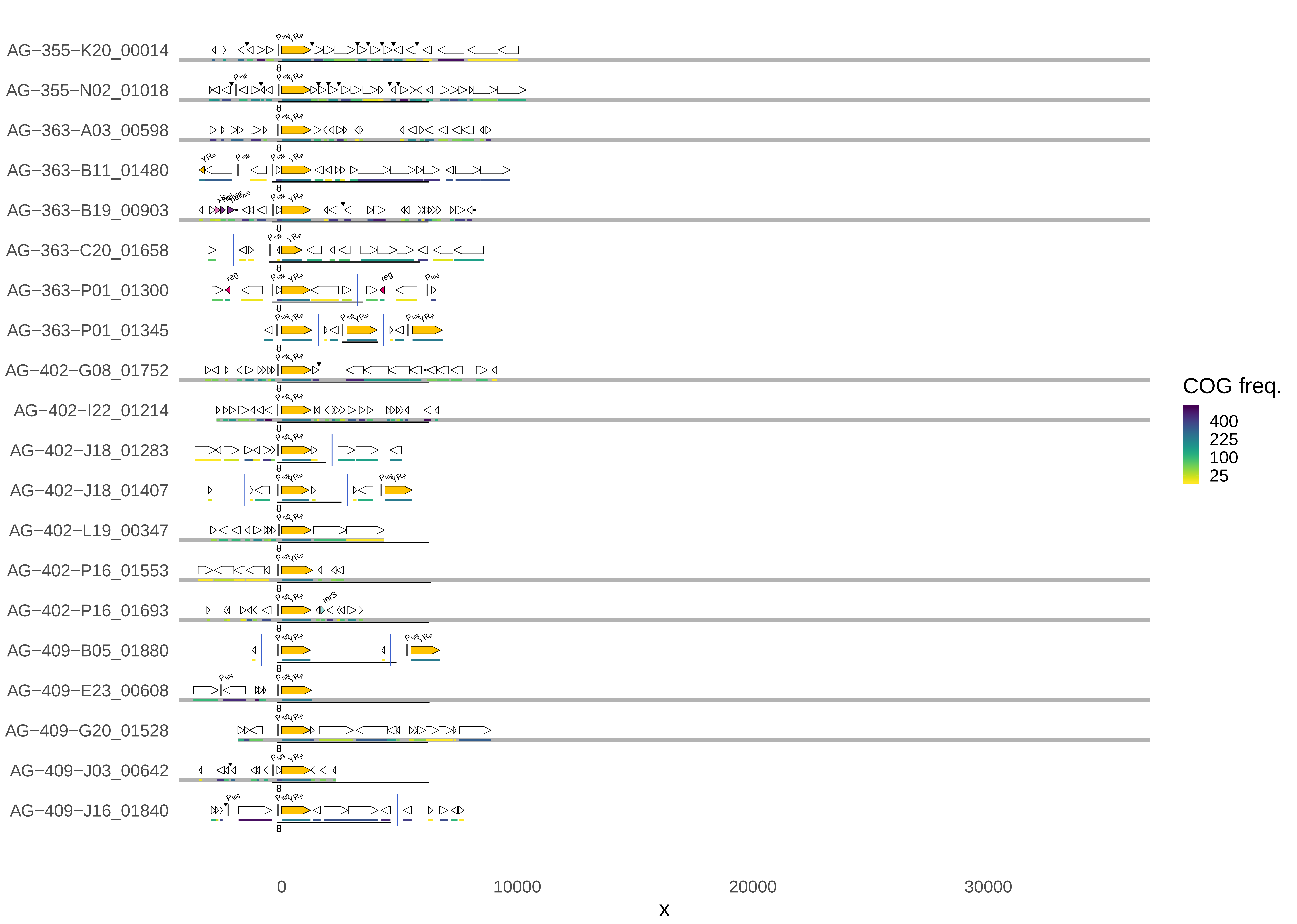

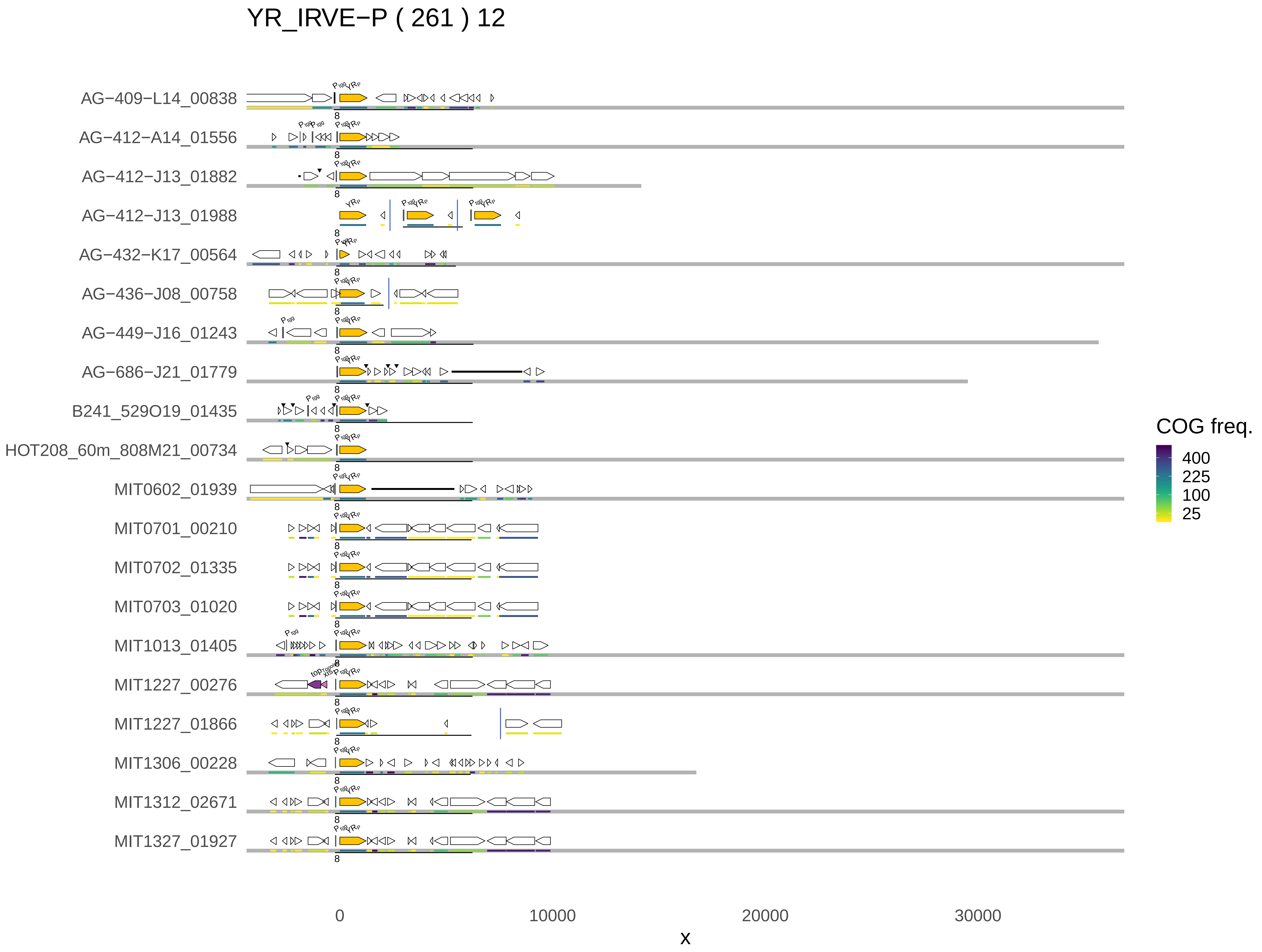

YR_IRVE−P ( 261 ) 12
AG−409−L14_00838
AG−412−A14_01556
AG−412−J13_01882
AG−412−J13_01988
AG−432−K17_00564
AG−436−J08_00758
AG−449−J16_01243
AG−686−J21_01779
B241_529O19_01435
HOT208_60m_808M21_00734
MIT0602_01939
MIT0701_00210
MIT0702_01335
MIT0703_01020
MIT1013_01405
MIT1227_00276
MIT1227_01866
MIT1306_00228
MIT1312_02671
MIT1327_01927
COG freq.
400
225
100
25
0
10000
20000
30000
x

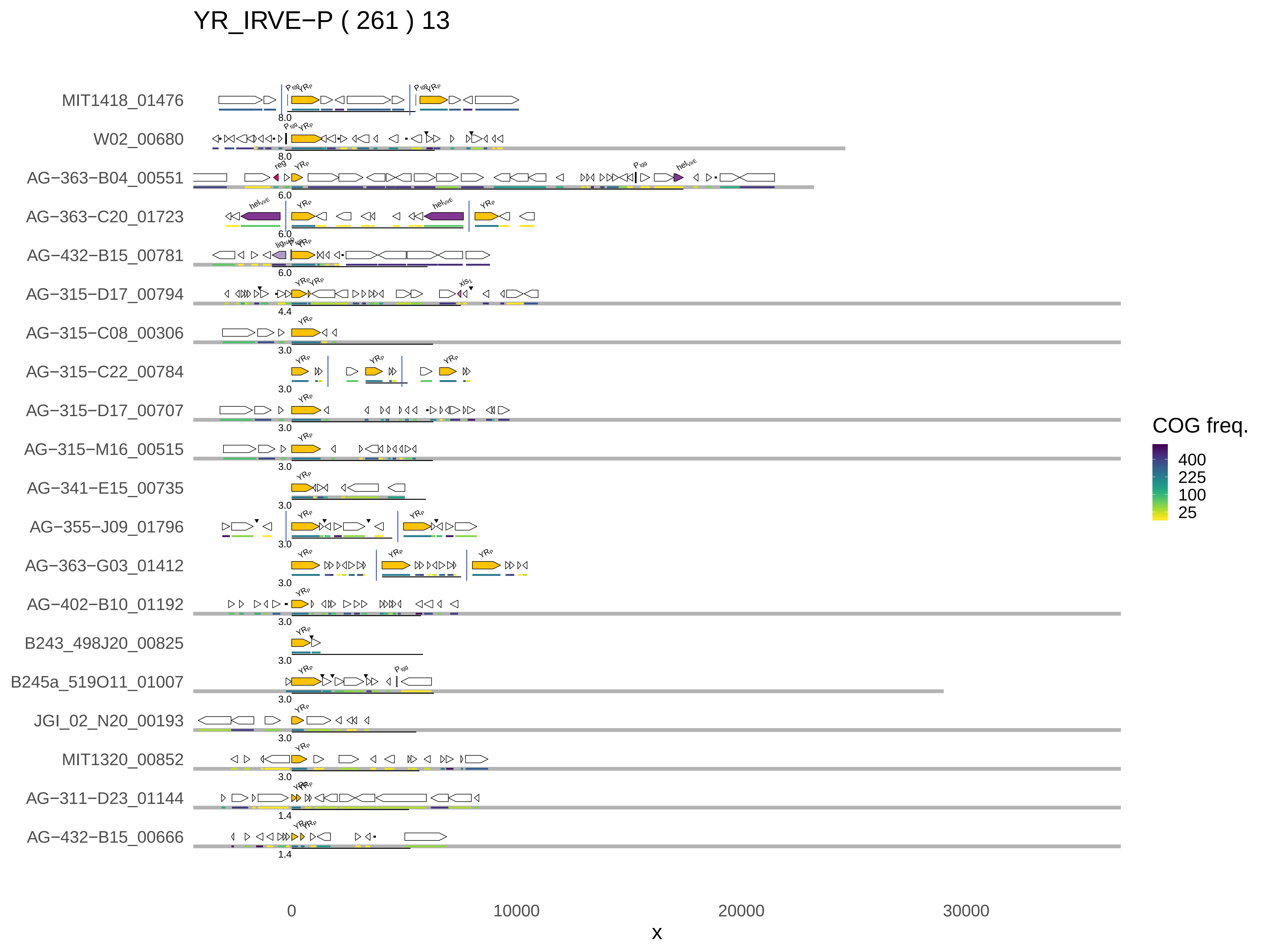
YR_IRVE−P ( 261 ) 13
MIT1418_01476
W02_00680
AG−363−B04_00551
AG−363−C20_01723
AG−432−B15_00781
AG−315−D17_00794
AG−315−C08_00306
AG−315−C22_00784
AG−315−D17_00707
AG−315−M16_00515
AG−341−E15_00735
AG−355−J09_01796
AG−363−G03_01412
AG−402−B10_01192
B243_498J20_00825
B245a_519O11_01007
JGI_02_N20_00193
MIT1320_00852
AG−311−D23_01144
AG−432−B15_00666
0
10000
20000
30000
x
COG freq.
400
225
100
25

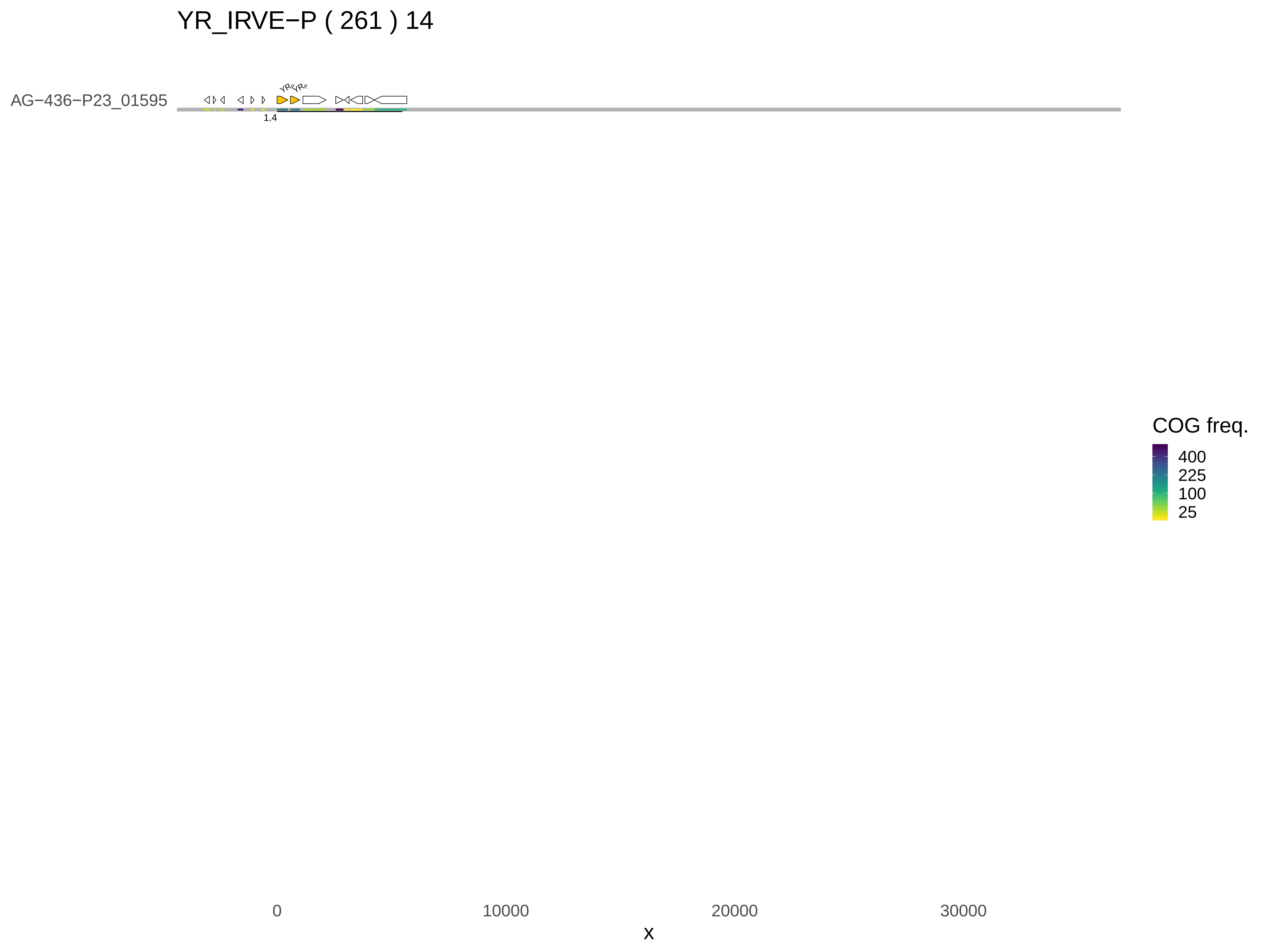

YR_IRVE−P ( 261 ) 14
AG−436−P23_01595
YR
YR
1.4
COG freq.
400
225
100
25
0
10000
20000
30000
x

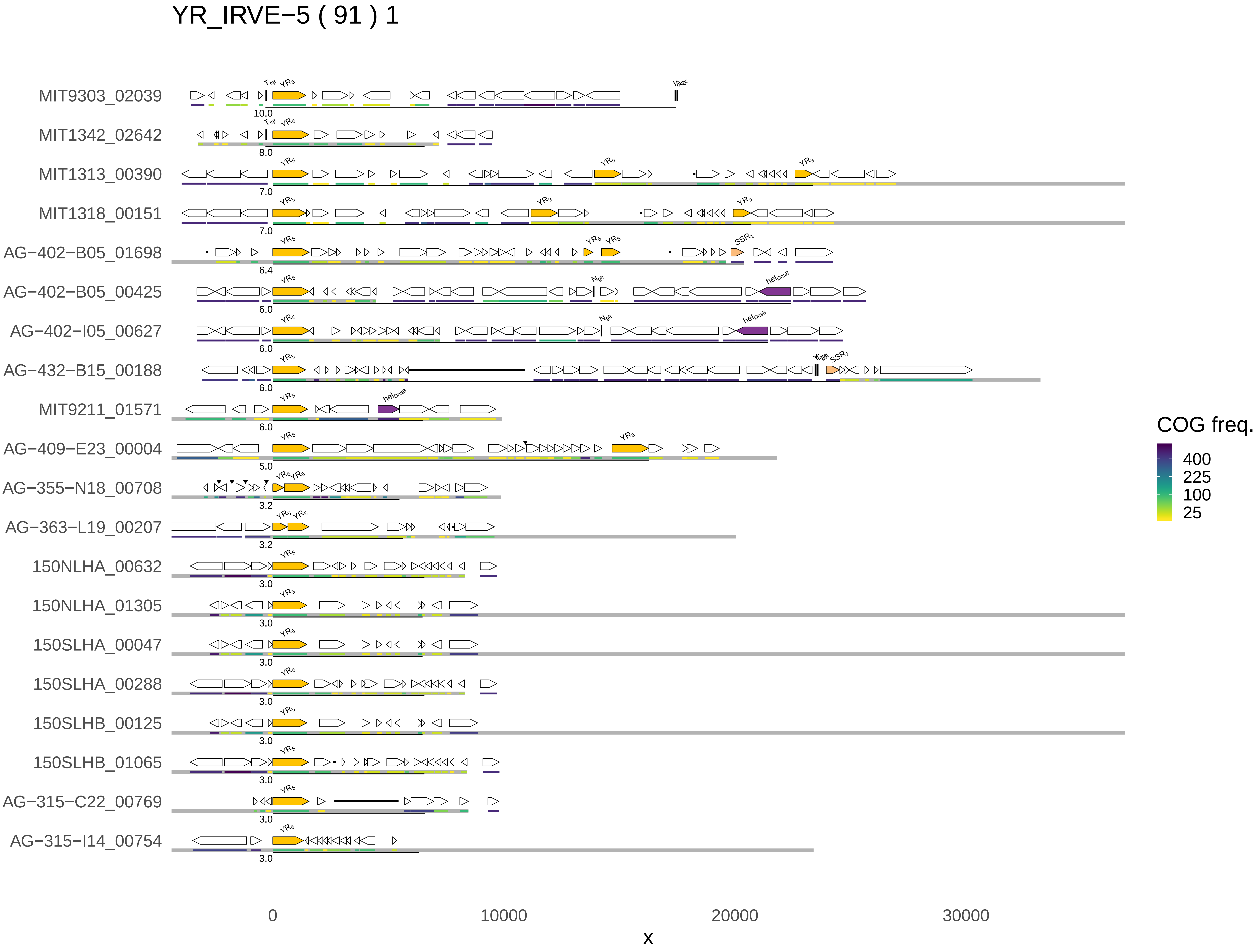

YR_IRVE−5 ( 91 ) 1
MIT9303_02039
MIT1342_02642
MIT1313_00390
MIT1318_00151
AG−402−B05_01698
AG−402−B05_00425
AG−402−I05_00627
AG−432−B15_00188
MIT9211_01571
AG−409−E23_00004
AG−355−N18_00708
AG−363−L19_00207
150NLHA_00632
150NLHA_01305
150SLHA_00047
150SLHA_00288
150SLHB_00125
150SLHB_01065
AG−315−C22_00769
AG−315−I14_00754
10.0
8.0
7.0
7.0
6.4
6.0
6.0
6.0
6.0
5.0
3.2
3.2
3.0
3.0
3.0
3.0
3.0
3.0
3.0
3.0
0
10000
20000
30000
x
COG freq.
400
225
100
25

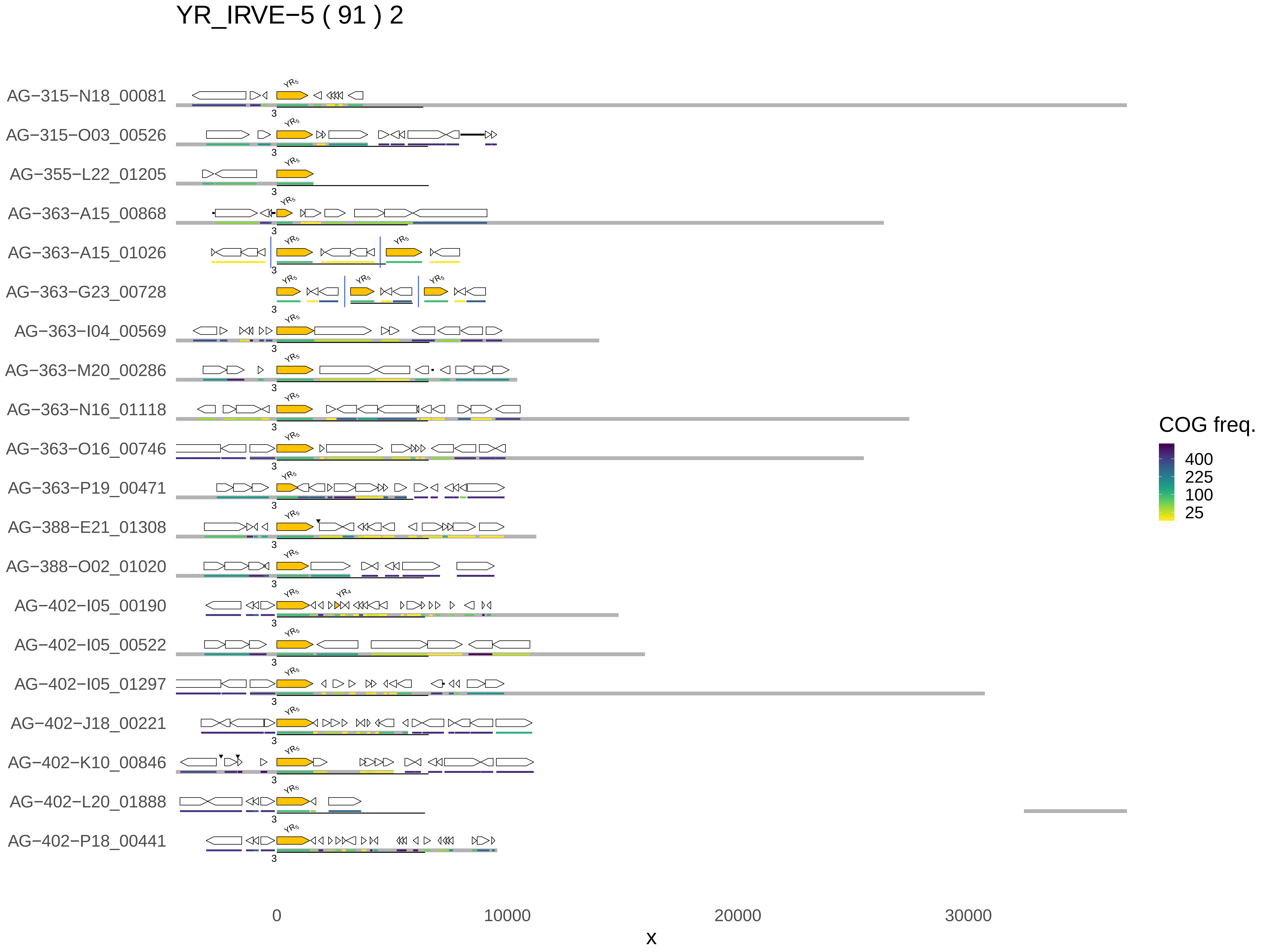
YR_IRVE−5 ( 91 ) 2
AG−315−N18_00081
AG−315−O03_00526
AG−355−L22_01205
AG−363−A15_00868
AG−363−A15_01026
AG−363−G23_00728
AG−363−I04_00569
AG−363−M20_00286
AG−363−N16_01118
AG−363−O16_00746
AG−363−P19_00471
AG−388−E21_01308
AG−388−O02_01020
AG−402−I05_00190
AG−402−I05_00522
AG−402−I05_01297
AG−402−J18_00221
AG−402−K10_00846
AG−402−L20_01888
AG−402−P18_00441
COG freq.
400
225
100
25
0
10000
20000
30000
x

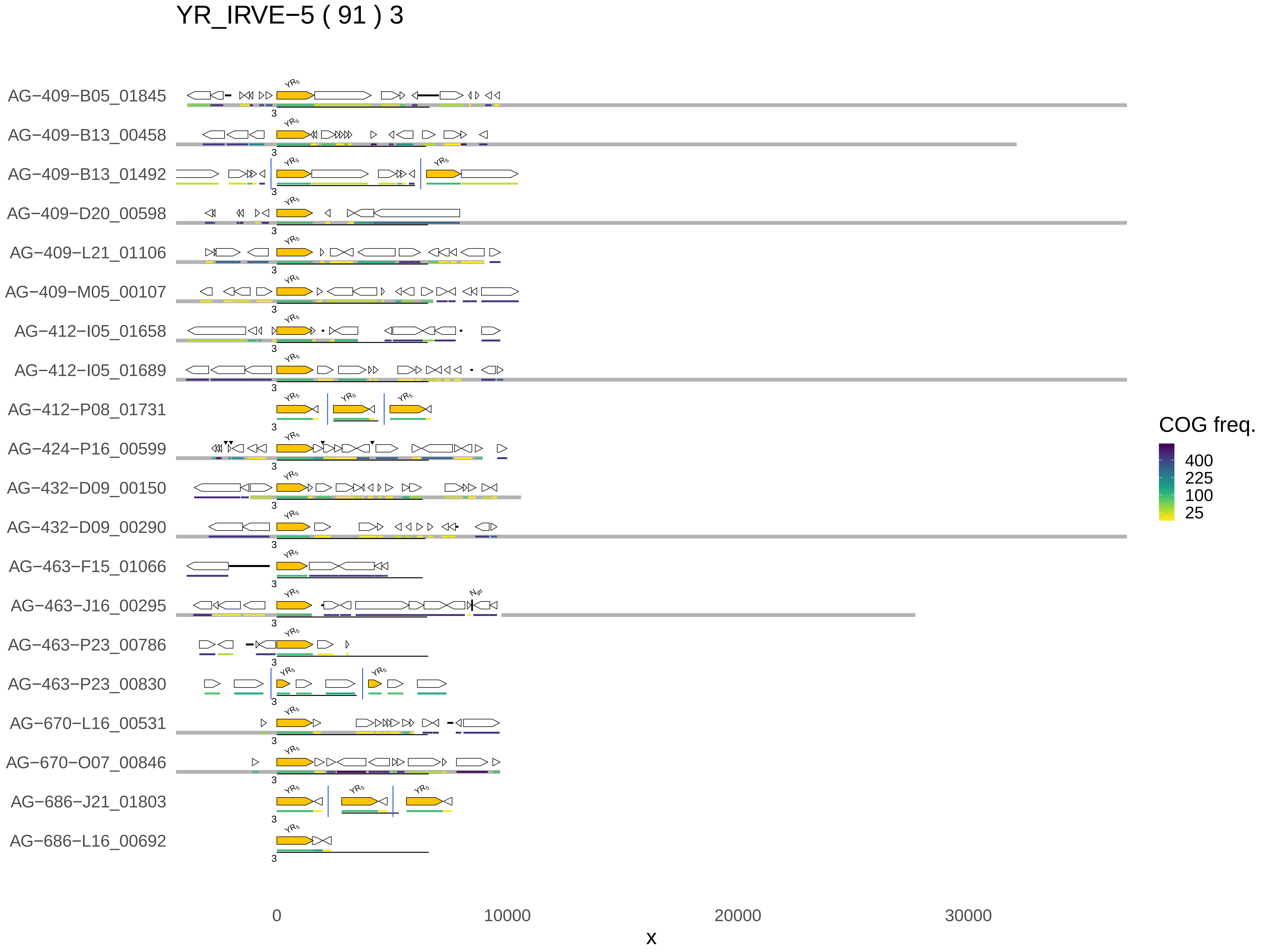

YR_IRVE−5 ( 91 ) 3
AG−409−B05_01845
AG−409−B13_00458
AG−409−B13_01492
AG−409−D20_00598
AG−409−L21_01106
AG−409−M05_00107
AG−412−I05_01658
AG−412−I05_01689
AG−412−P08_01731
AG−424−P16_00599
AG−432−D09_00150
AG−432−D09_00290
AG−463−F15_01066
AG−463−J16_00295
AG−463−P23_00786
AG−463−P23_00830
AG−670−L16_00531
AG−670−O07_00846
AG−686−J21_01803
AG−686−L16_00692
YR5
3
COG freq.
400
225
100
25
0
10000
20000
30000
x

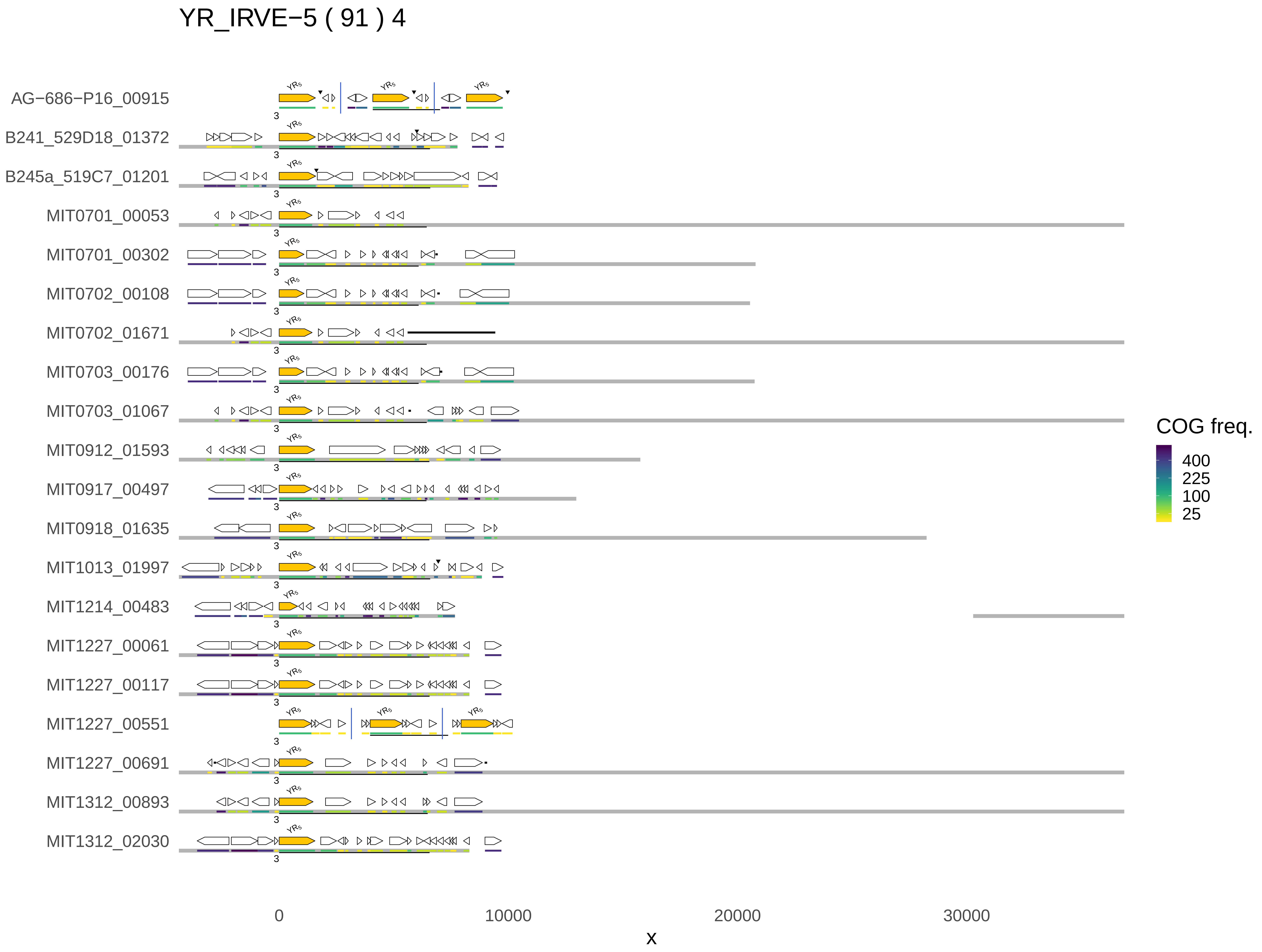
YR_IRVE−5 ( 91 ) 4
AG−686−P16_00915
B241_529D18_01372
B245a_519C7_01201
MIT0701_00053
MIT0701_00302
MIT0702_00108
MIT0702_01671
MIT0703_00176
MIT0703_01067
MIT0912_01593
MIT0917_00497
MIT0918_01635
MIT1013_01997
MIT1214_00483
MIT1227_00061
MIT1227_00117
MIT1227_00551
MIT1227_00691
MIT1312_00893
MIT1312_02030
YRs
COG freq.
400
225
100
25
0
10000
20000
30000
x

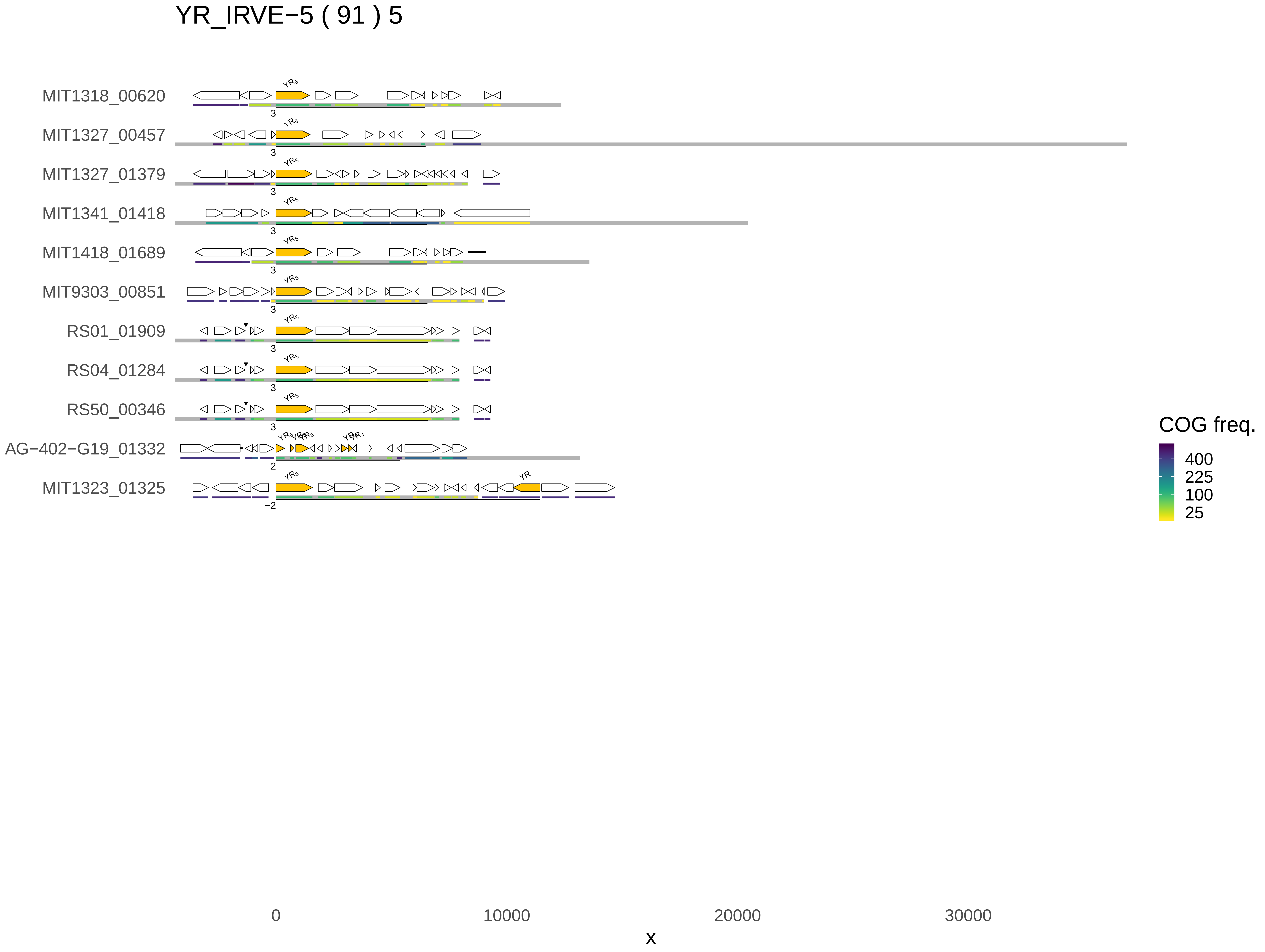
YR_IRVE−5 ( 91 ) 5
MIT1318_00620
MIT1327_00457
MIT1327_01379
MIT1341_01418
MIT1418_01689
MIT9303_00851
RS01_01909
RS04_01284
RS50_00346
AG−402−G19_01332
MIT1323_01325
COG freq.
400
225
100
25
0
10000
20000
30000
x

YR_IRVE−4 ( 74 ) 1

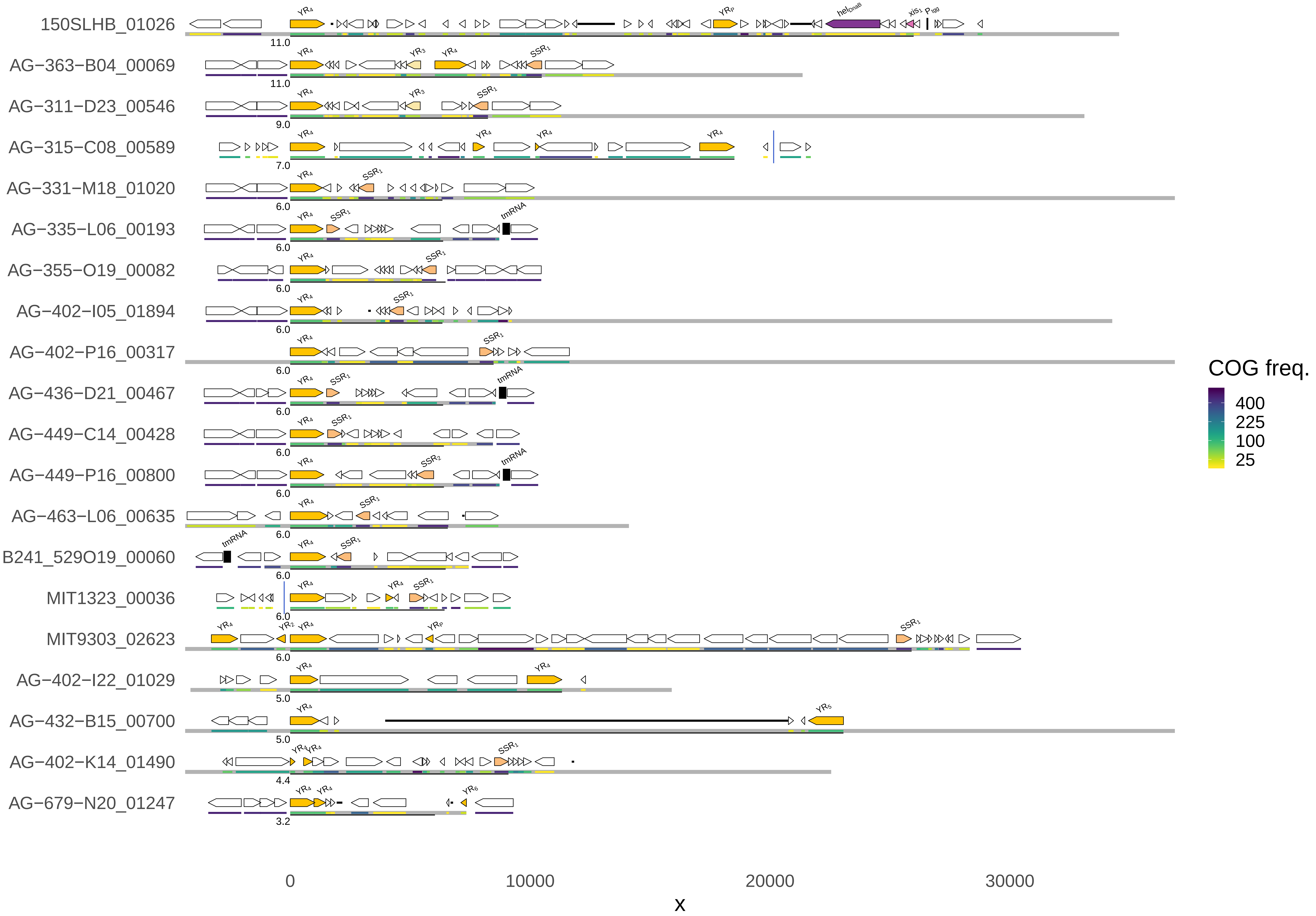
150SLHB_01026
AG−363−B04_00069
AG−311−D23_00546
AG−315−C08_00589
AG−331−M18_01020
AG−335−L06_00193
AG−355−O19_00082
AG−402−I05_01894
AG−402−P16_00317
AG−436−D21_00467
AG−449−C14_00428
AG−449−P16_00800
AG−463−L06_00635
B241_529O19_00060
MIT1323_00036
MIT9303_02623
AG−402−I22_01029
AG−432−B15_00700
AG−402−K14_01490
AG−679−N20_01247
11.0
11.0
9.0
7.0
6.0
6.0
6.0
6.0
6.0
6.0
6.0
6.0
6.0
6.0
6.0
6.0
5.0
5.0
4.4
3.2
0
10000
20000
30000
x
COG freq.
400
225
100
25

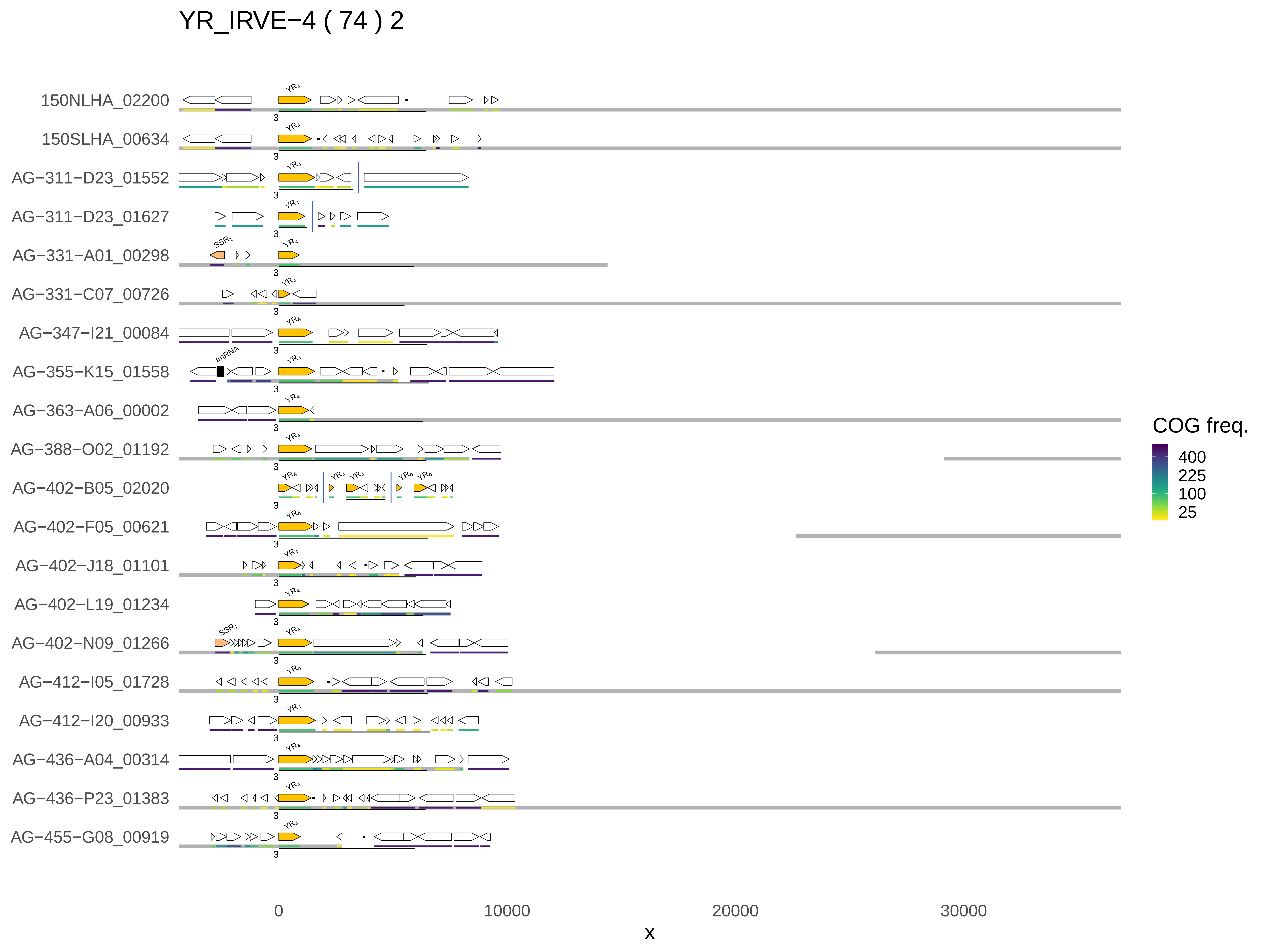

YR_IRVE−4 ( 74 ) 2
150NLHA_02200
150SLHA_00634
AG−311−D23_01552
AG−311−D23_01627
AG−331−A01_00298
AG−331−C07_00726
AG−347−I21_00084
AG−355−K15_01558
AG−363−A06_00002
AG−388−O02_01192
AG−402−B05_02020
AG−402−F05_00621
AG−402−J18_01101
AG−402−L19_01234
AG−402−N09_01266
AG−412−I05_01728
AG−412−I20_00933
AG−436−A04_00314
AG−436−P23_01383
AG−455−G08_00919
YR4
SSR1
tmRNA
3
0
10000
20000
30000
x
COG freq.
400
225
100
25

YR_IRVE−4 ( 74 ) 3

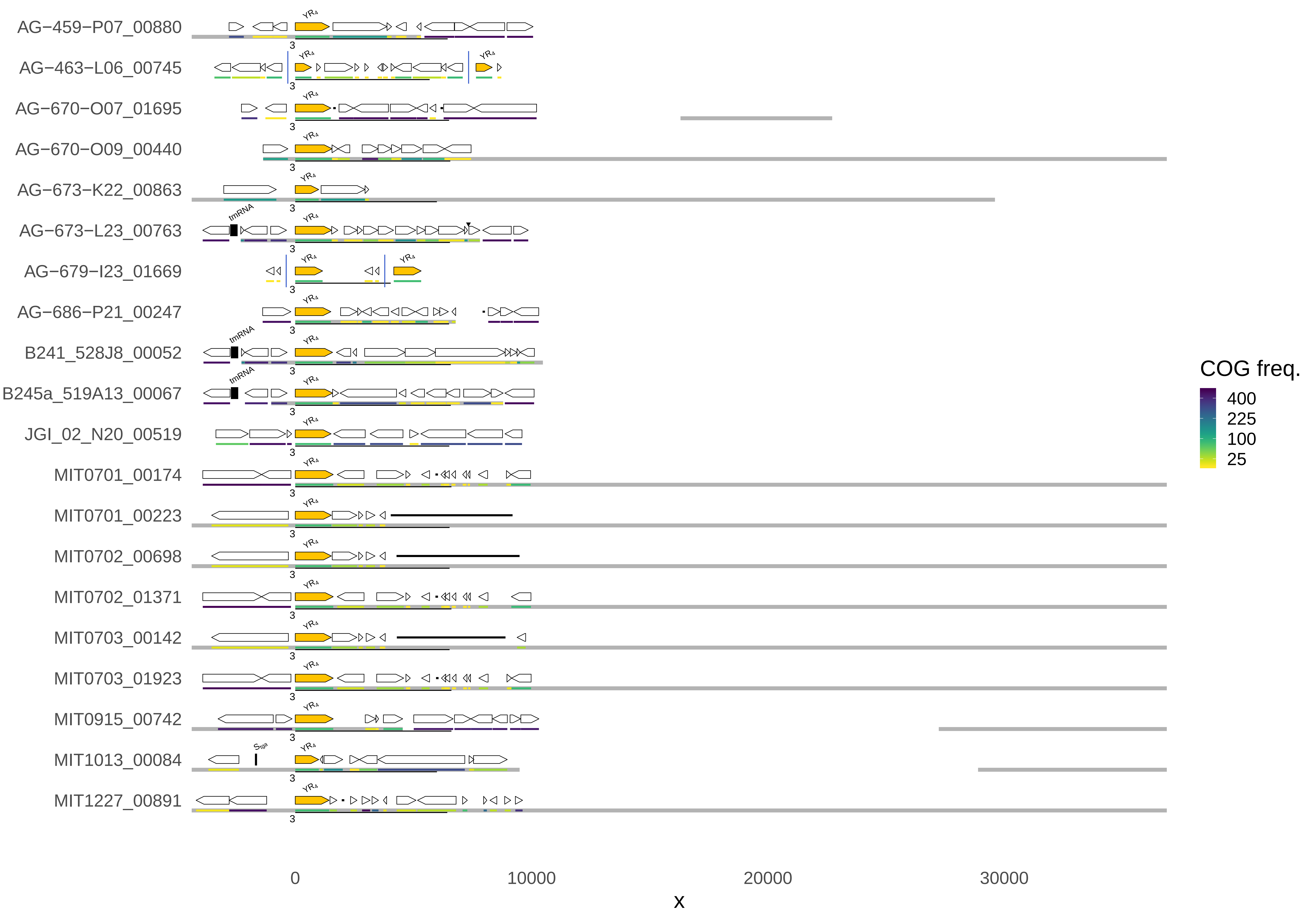
AG−459−P07_00880
AG−463−L06_00745
AG−670−O07_01695
AG−670−O09_00440
AG−673−K22_00863
AG−673−L23_00763
AG−679−I23_01669
AG−686−P21_00247
B241_528J8_00052
B245a_519A13_00067
JGI_02_N20_00519
MIT0701_00174
MIT0701_00223
MIT0702_00698
MIT0702_01371
MIT0703_00142
MIT0703_01923
MIT0915_00742
MIT1013_00084
MIT1227_00891
tmRNA
0
10000
20000
30000
x
COG freq.
400
225
100
25

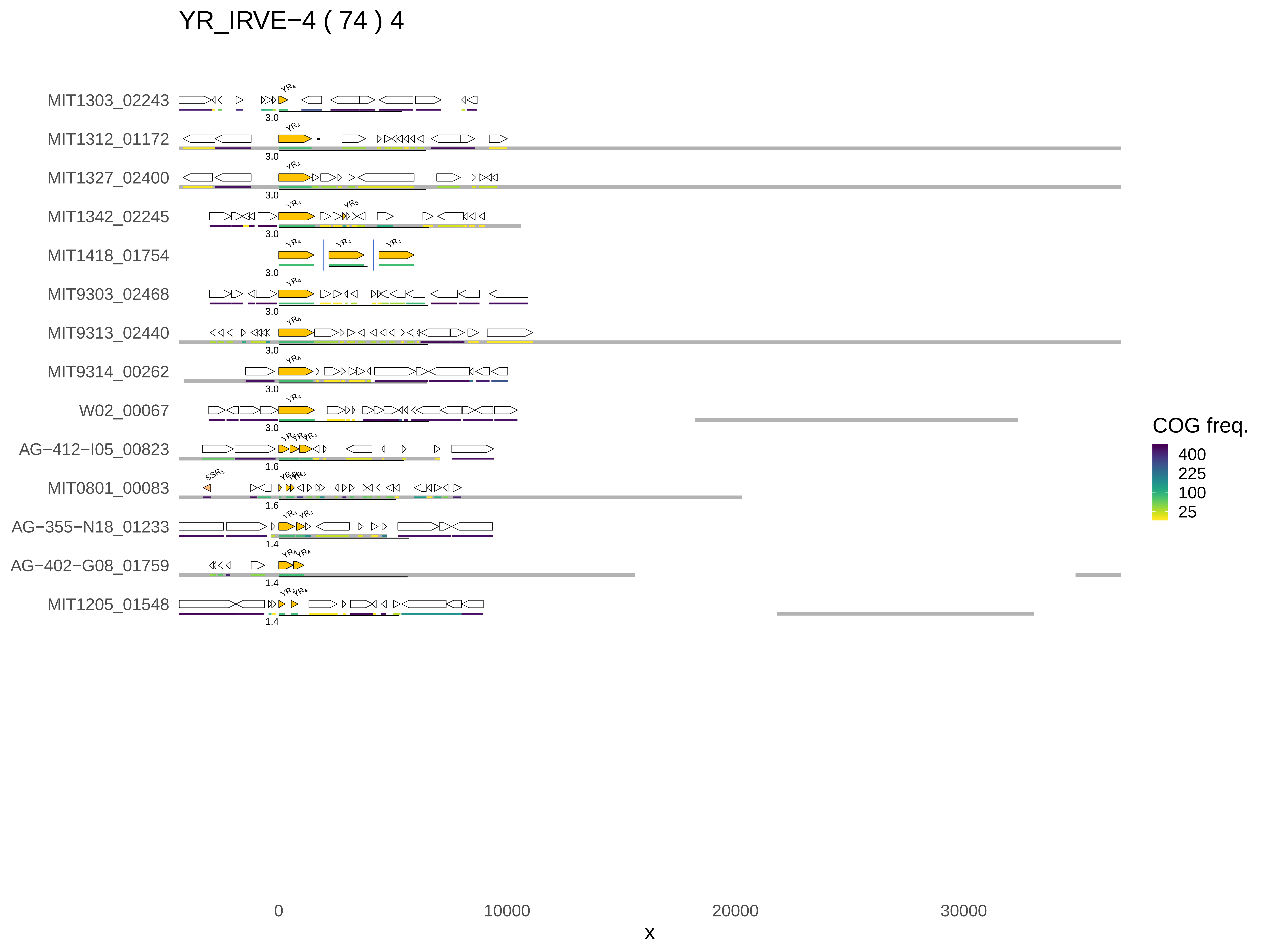
YR_IRVE−4 ( 74 ) 4
MIT1303_02243
MIT1312_01172
MIT1327_02400
MIT1342_02245
MIT1418_01754
MIT9303_02468
MIT9313_02440
MIT9314_00262
W02_00067
AG−412−I05_00823
MIT0801_00083
AG−355−N18_01233
AG−402−G08_01759
MIT1205_01548
3.0
1.6
1.4
SSR1
0
10000
20000
30000
x
COG freq.
400
225
100
25

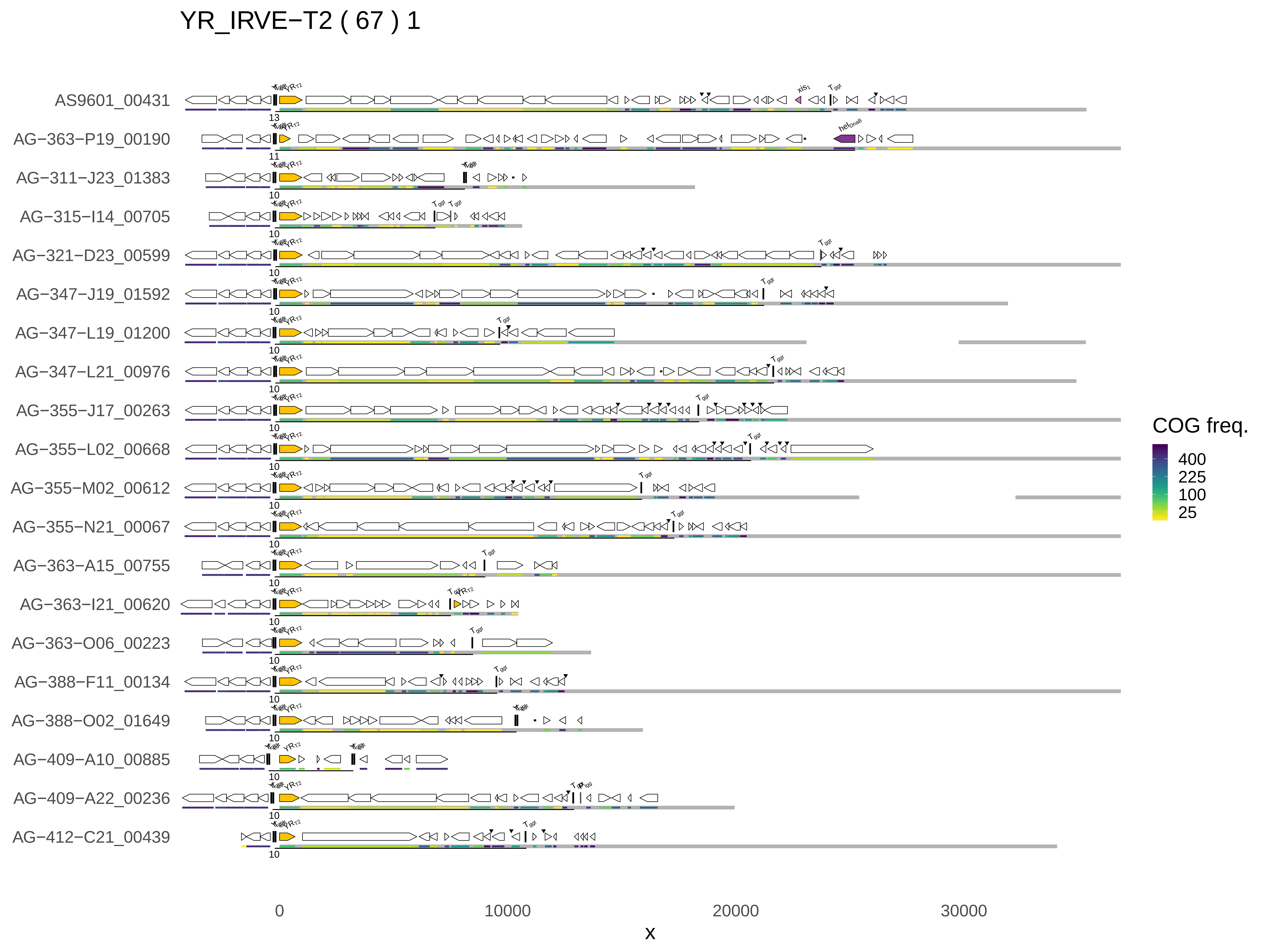
YR_IRVE−T2 ( 67 ) 1
AS9601_00431
AG−363−P19_00190
AG−311−J23_01383
AG−315−I14_00705
AG−321−D23_00599
AG−347−J19_01592
AG−347−L19_01200
AG−347−L21_00976
AG−355−J17_00263
AG−355−L02_00668
AG−355−M02_00612
AG−355−N21_00067
AG−363−A15_00755
AG−363−I21_00620
AG−363−O06_00223
AG−388−F11_00134
AG−388−O02_01649
AG−409−A10_00885
AG−409−A22_00236
AG−412−C21_00439
COG freq.
400
225
100
25
0
10000
20000
30000
x

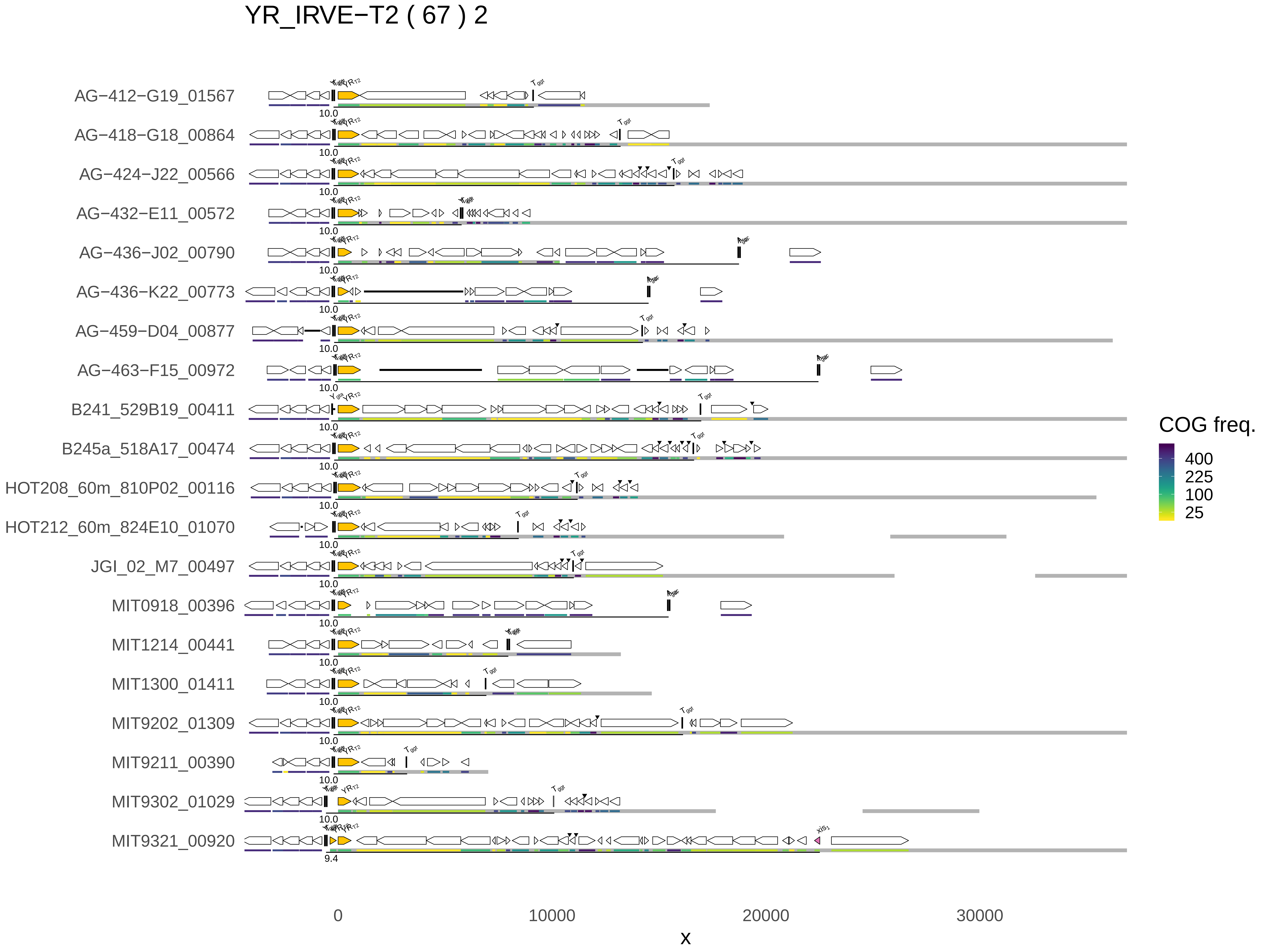
YR_IRVE−T2 ( 67 ) 2
AG−412−G19_01567
AG−418−G18_00864
AG−424−J22_00566
AG−432−E11_00572
AG−436−J02_00790
AG−436−K22_00773
AG−459−D04_00877
AG−463−F15_00972
B241_529B19_00411
B245a_518A17_00474
HOT208_60m_810P02_00116
HOT212_60m_824E10_01070
JGI_02_M7_00497
MIT0918_00396
MIT1214_00441
MIT1300_01411
MIT9202_01309
MIT9211_00390
MIT9302_01029
MIT9321_00920
10.0
9.4
0
10000
20000
30000
x
COG freq.
400
225
100
25

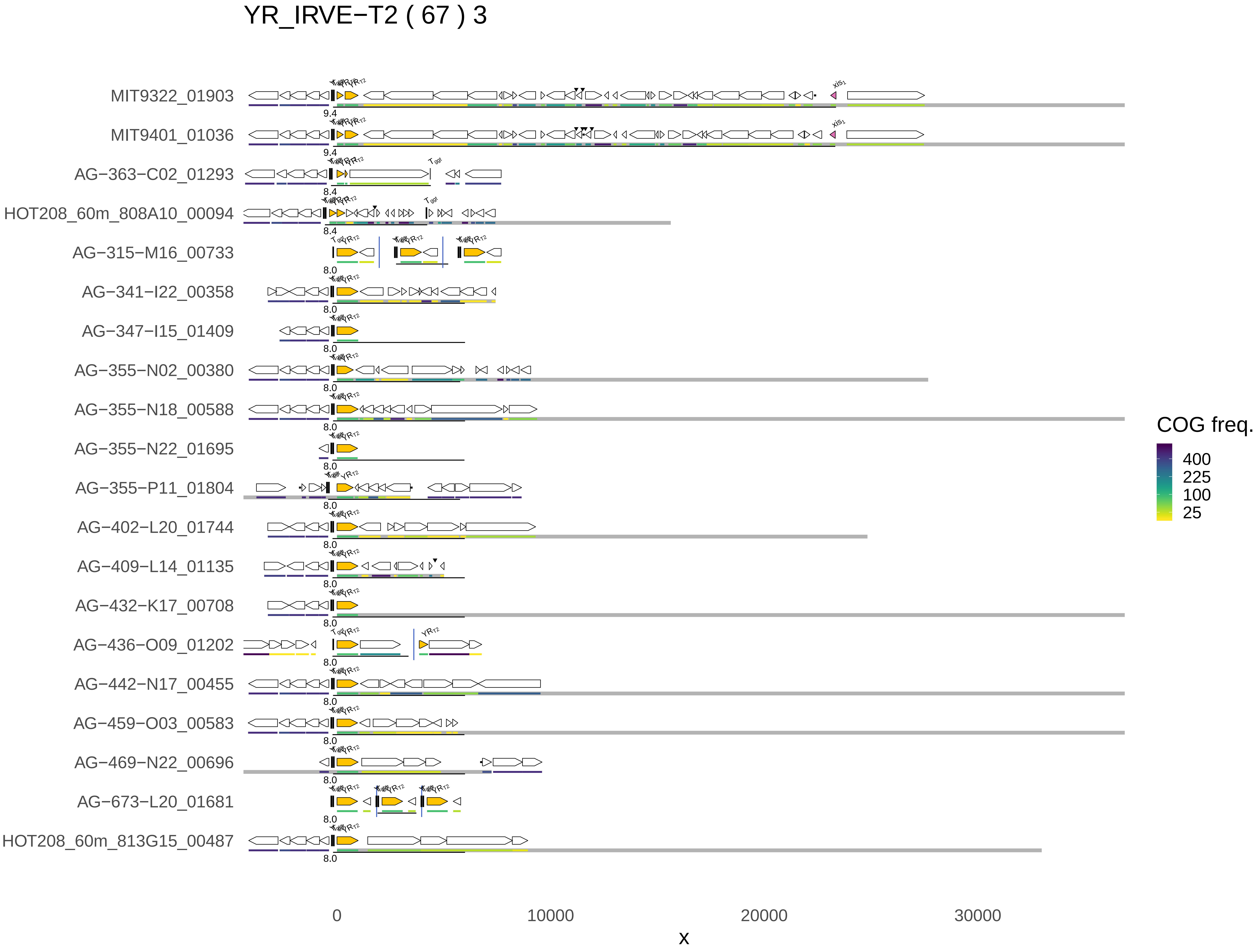
YR_IRVE−T2 ( 67 ) 3
MIT9322_01903
MIT9401_01036
AG−363−C02_01293
HOT208_60m_808A10_00094
AG−315−M16_00733
AG−341−I22_00358
AG−347−I15_01409
AG−355−N02_00380
AG−355−N18_00588
AG−355−N22_01695
AG−355−P11_01804
AG−402−L20_01744
AG−409−L14_01135
AG−432−K17_00708
AG−436−O09_01202
AG−442−N17_00455
AG−459−O03_00583
AG−469−N22_00696
AG−673−L20_01681
HOT208_60m_813G15_00487
9.4
8.4
8.0
0
10000
20000
30000
x
COG freq.
400
225
100
25

### YR_IRVE−T2 ( 67 ) 4

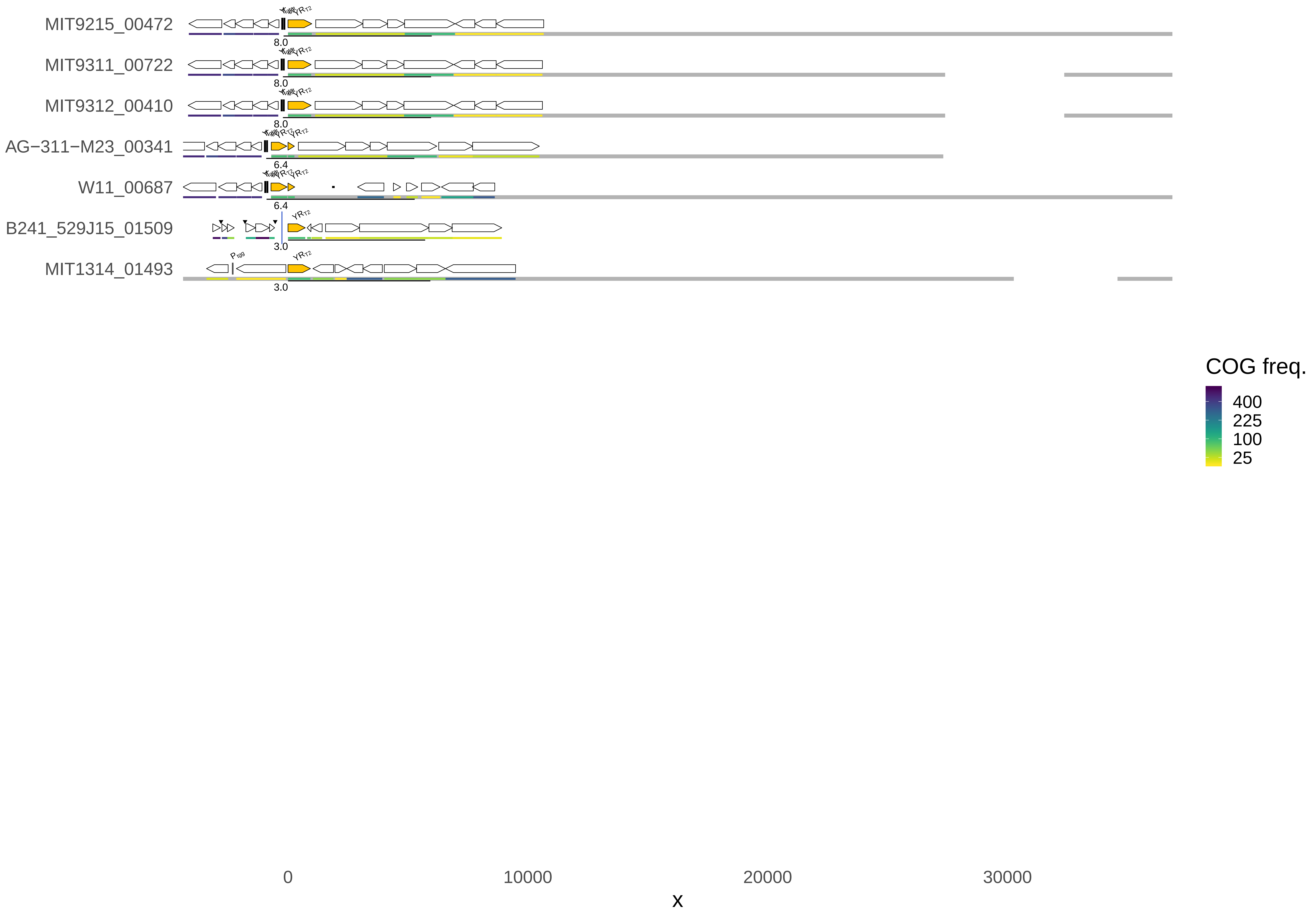

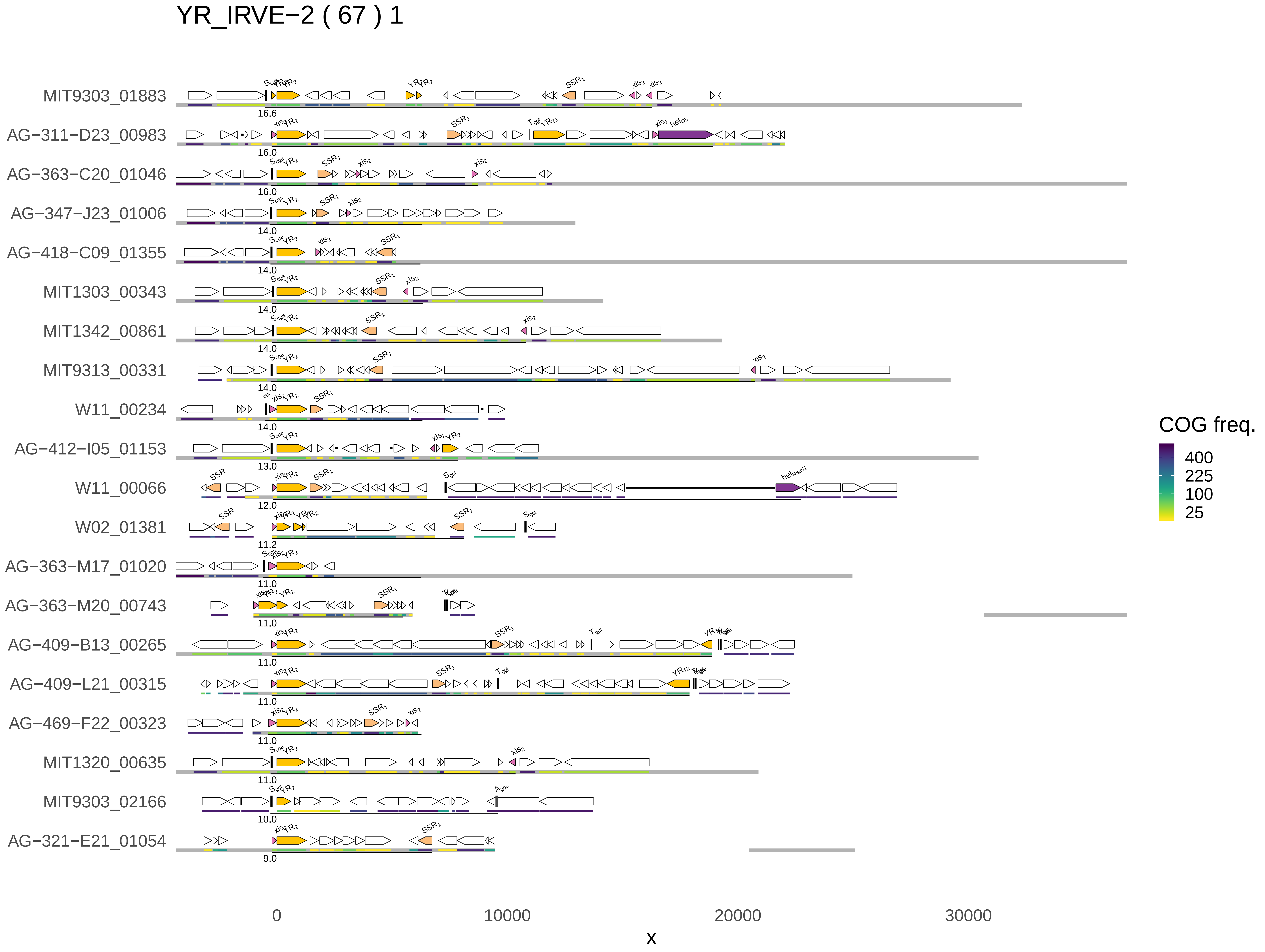

YR_IRVE−2 ( 67 ) 1
MIT9303_01883
AG−311−D23_00983
AG−363−C20_01046
AG−347−J23_01006
AG−418−C09_01355
MIT1303_00343
MIT1342_00861
MIT9313_00331
W11_00234
AG−412−I05_01153
W11_00066
W02_01381
AG−363−M17_01020
AG−363−M20_00743
AG−409−B13_00265
AG−409−L21_00315
AG−469−F22_00323
MIT1320_00635
MIT9303_02166
AG−321−E21_01054
COG freq.
400
225
100
25
0
10000
20000
30000
x

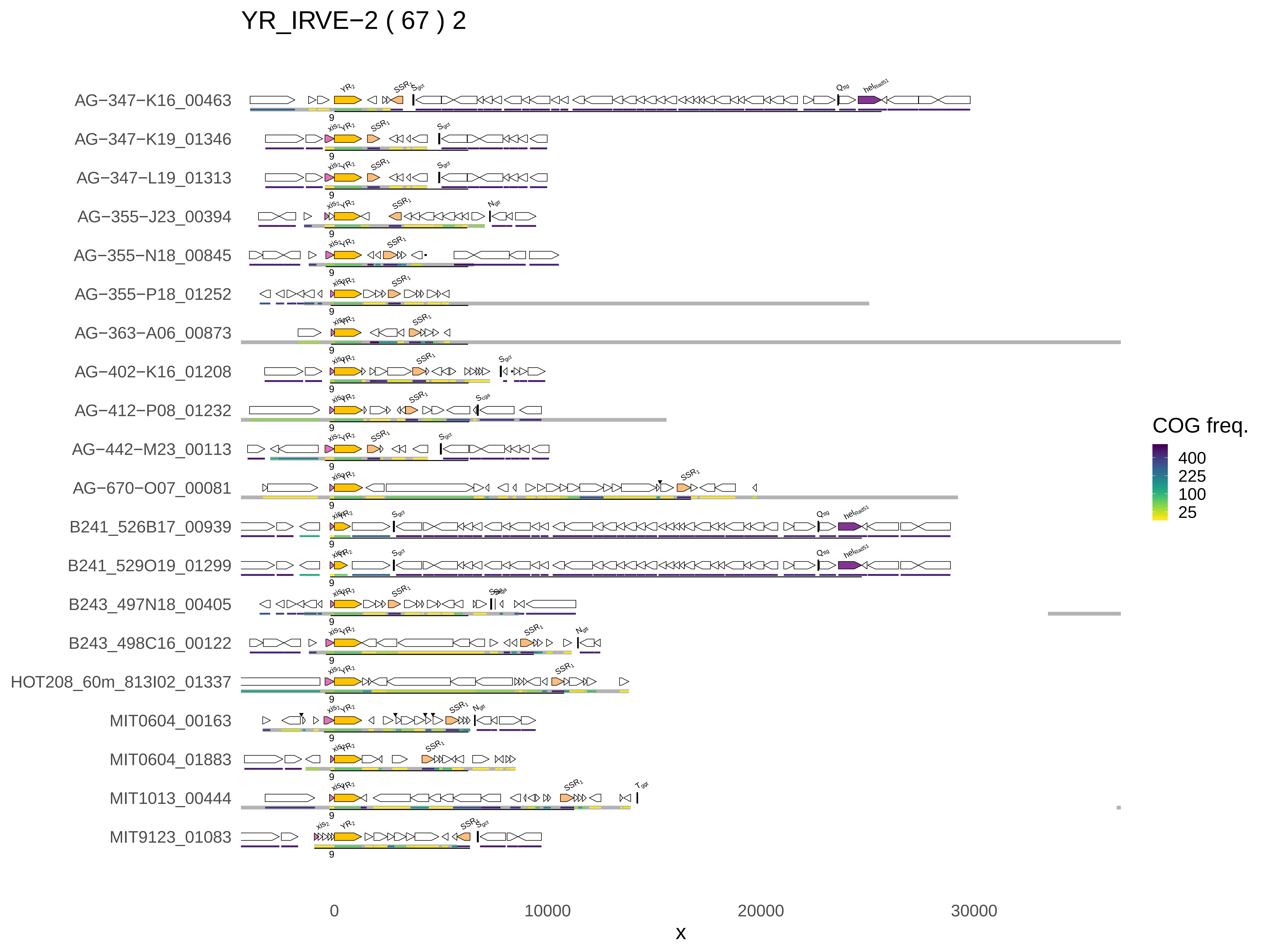
YR_IRVE−2 ( 67 ) 2
AG−347−K16_00463
AG−347−K19_01346
AG−347−L19_01313
AG−355−J23_00394
AG−355−N18_00845
AG−355−P18_01252
AG−363−A06_00873
AG−402−K16_01208
AG−412−P08_01232
AG−442−M23_00113
AG−670−O07_00081
B241_526B17_00939
B241_529O19_01299
B243_497N18_00405
B243_498C16_00122
HOT208_60m_813I02_01337
MIT0604_00163
MIT0604_01883
MIT1013_00444
MIT9123_01083
0
10000
20000
30000
x
COG freq.
400
225
100
25

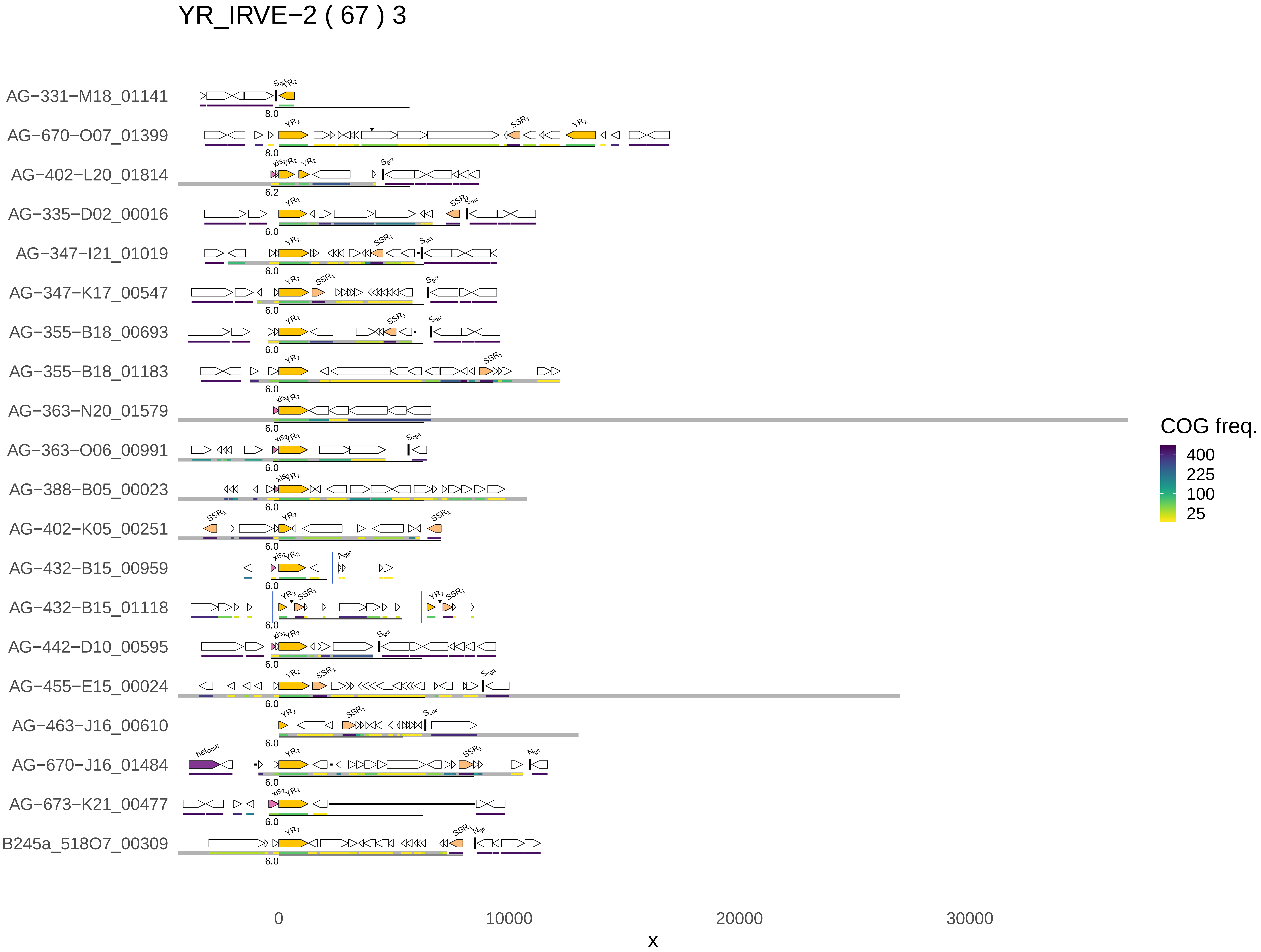
YR_IRVE−2 ( 67 ) 3
AG−331−M18_01141
AG−670−O07_01399
AG−402−L20_01814
AG−335−D02_00016
AG−347−I21_01019
AG−347−K17_00547
AG−355−B18_00693
AG−355−B18_01183
AG−363−N20_01579
AG−363−O06_00991
AG−388−B05_00023
AG−402−K05_00251
AG−432−B15_00959
AG−432−B15_01118
AG−442−D10_00595
AG−455−E15_00024
AG−463−J16_00610
AG−670−J16_01484
AG−673−K21_00477
B245a_518O7_00309
0
10000
20000
30000
x
COG freq.
400
225
100
25

YR_IRVE−2 ( 67 ) 4

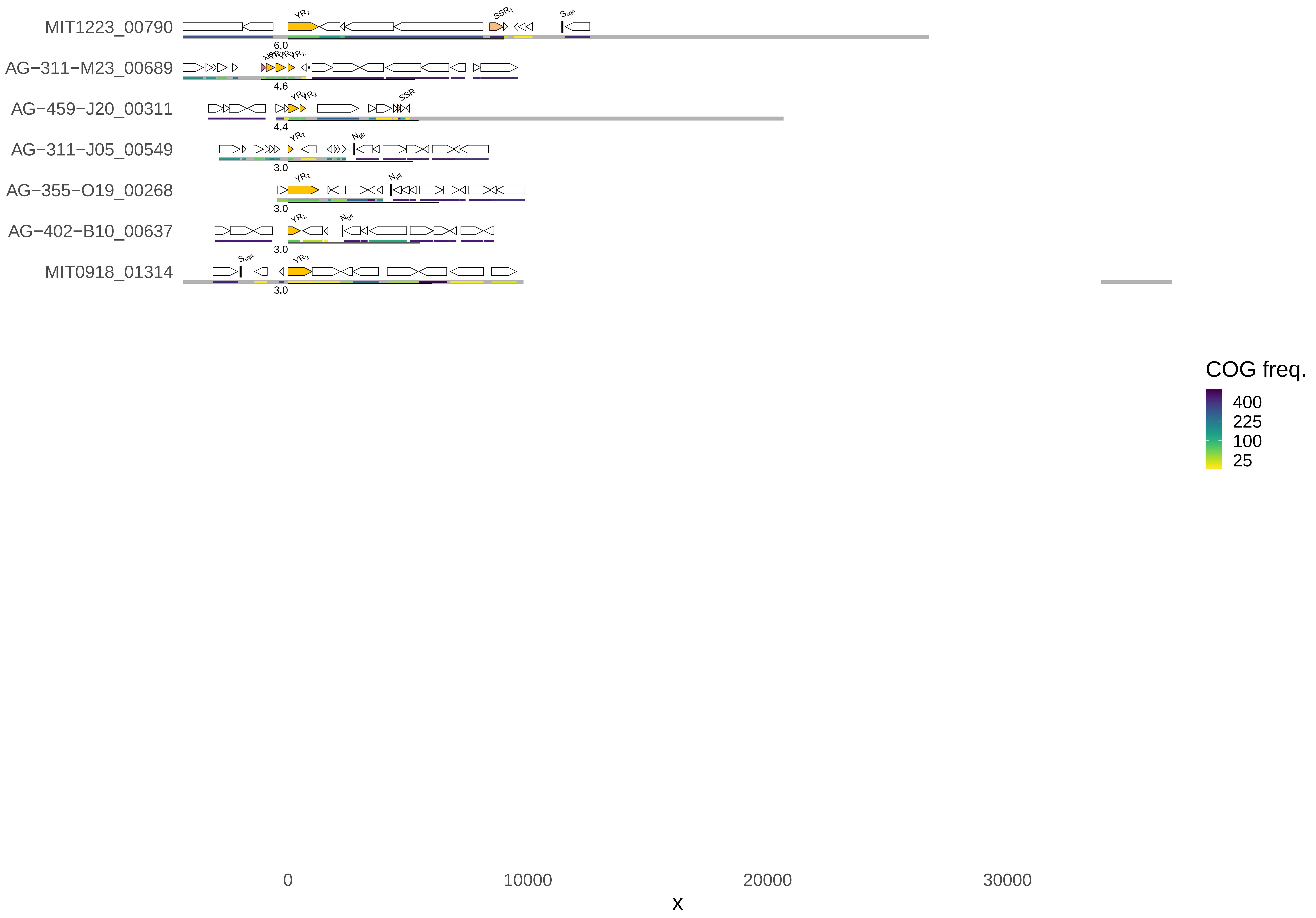
MIT1223_00790
AG−311−M23_00689
AG−459−J20_00311
AG−311−J05_00549
AG−355−O19_00268
AG−402−B10_00637
MIT0918_01314
6.0
4.6
4.4
3.0
3.0
3.0
3.0
COG freq.
400
225
100
25
0
10000
20000
30000
x

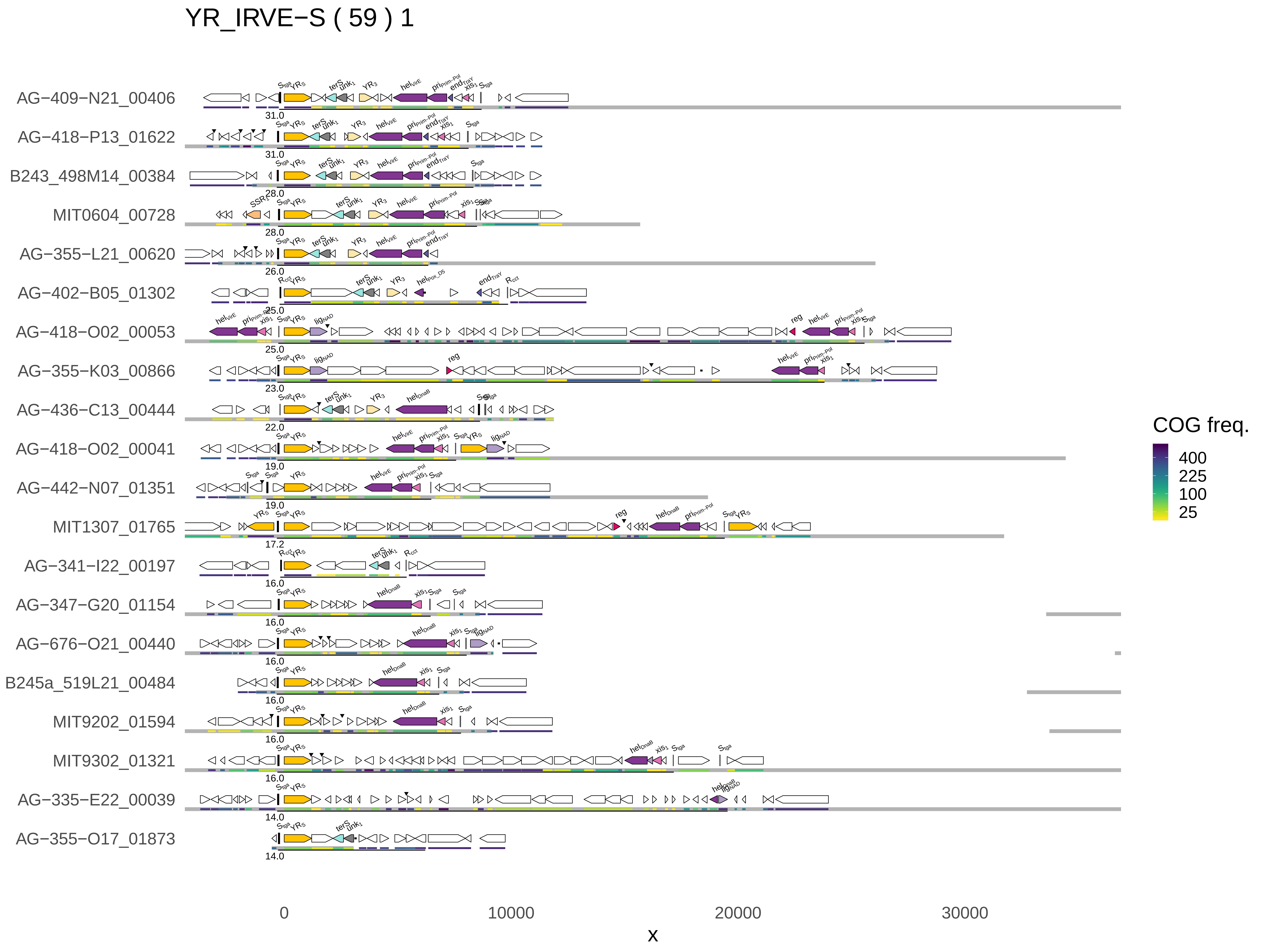

YR_IRVE−S ( 59 ) 1
AG−409−N21_00406
AG−418−P13_01622
B243_498M14_00384
MIT0604_00728
AG−355−L21_00620
AG−402−B05_01302
AG−418−O02_00053
AG−355−K03_00866
AG−436−C13_00444
AG−418−O02_00041
AG−442−N07_01351
MIT1307_01765
AG−341−I22_00197
AG−347−G20_01154
AG−676−O21_00440
B245a_519L21_00484
MIT9202_01594
MIT9302_01321
AG−335−E22_00039
AG−355−O17_01873
31.0
31.0
28.0
28.0
26.0
25.0
25.0
23.0
22.0
19.0
19.0
17.2
16.0
16.0
16.0
16.0
16.0
16.0
14.0
14.0
0
10000
20000
30000
x
COG freq.
400
225
100
25

YR_IRVE−S ( 59 ) 2

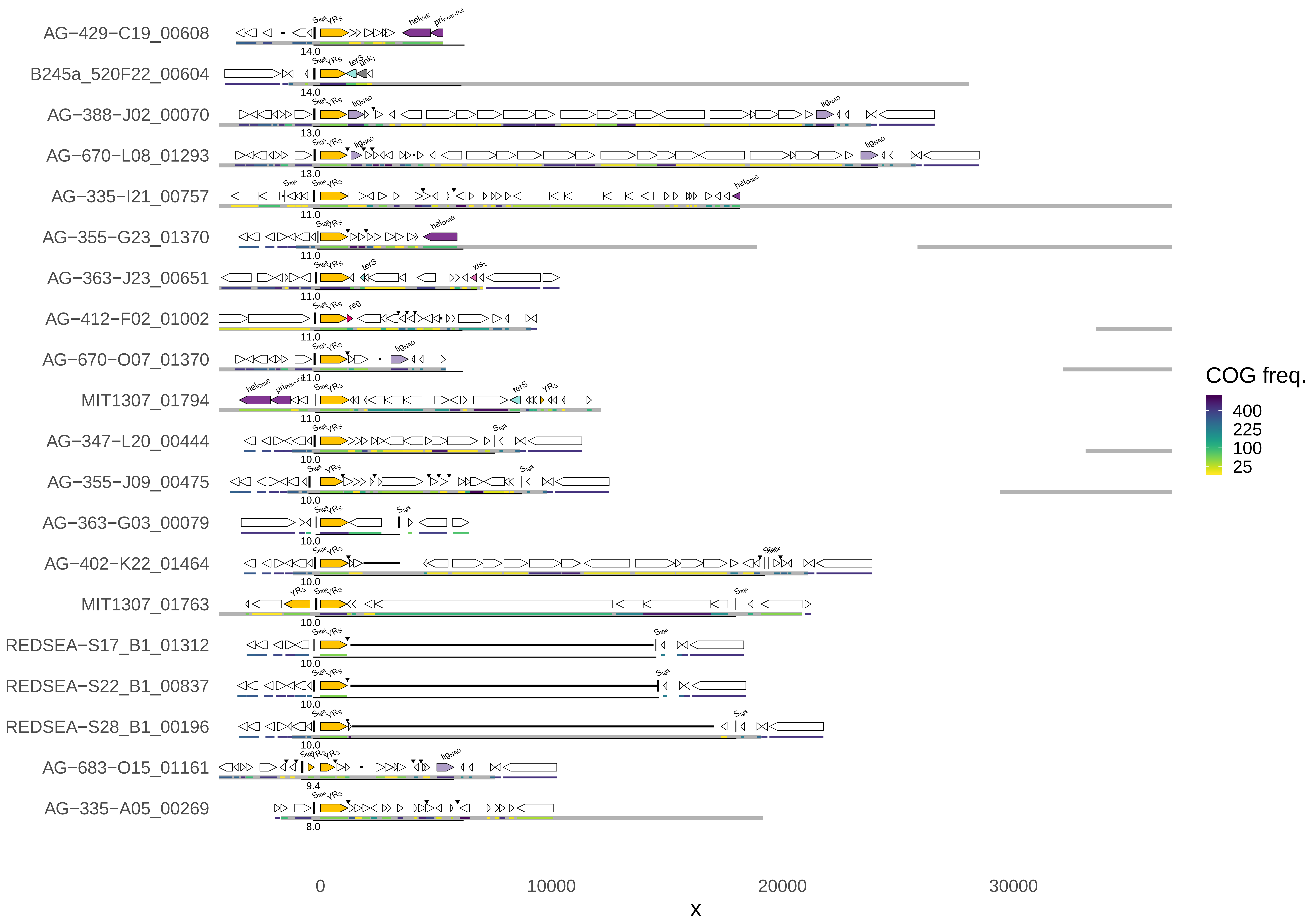
AG−429−C19_00608
B245a_520F22_00604
AG−388−J02_00070
AG−670−L08_01293
AG−335−I21_00757
AG−355−G23_01370
AG−363−J23_00651
AG−412−F02_01002
AG−670−O07_01370
MIT1307_01794
AG−347−L20_00444
AG−355−J09_00475
AG−363−G03_00079
AG−402−K22_01464
MIT1307_01763
REDSEA−S17_B1_01312
REDSEA−S22_B1_00837
REDSEA−S28_B1_00196
AG−683−O15_01161
AG−335−A05_00269
COG freq.
400
225
100
25
0
10000
20000
30000
x

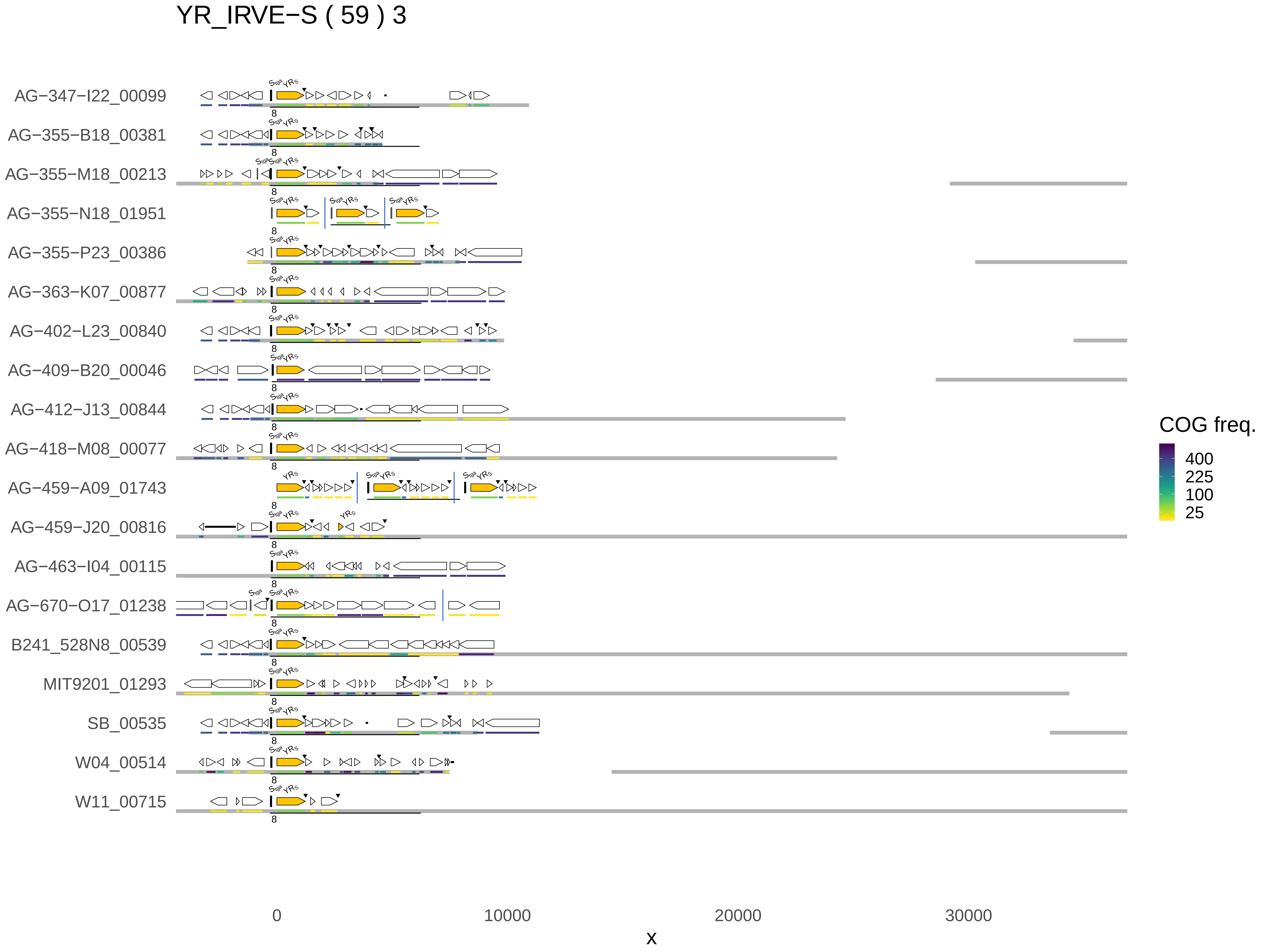

YR_IRVE−S ( 59 ) 3
AG−347−I22_00099
AG−355−B18_00381
AG−355−M18_00213
AG−355−N18_01951
AG−355−P23_00386
AG−363−K07_00877
AG−402−L23_00840
AG−409−B20_00046
AG−412−J13_00844
AG−418−M08_00077
AG−459−A09_01743
AG−459−J20_00816
AG−463−I04_00115
AG−670−O17_01238
B241_528N8_00539
MIT9201_01293
SB_00535
W04_00514
W11_00715
0
10000
20000
30000
x
COG freq.
400
225
100
25

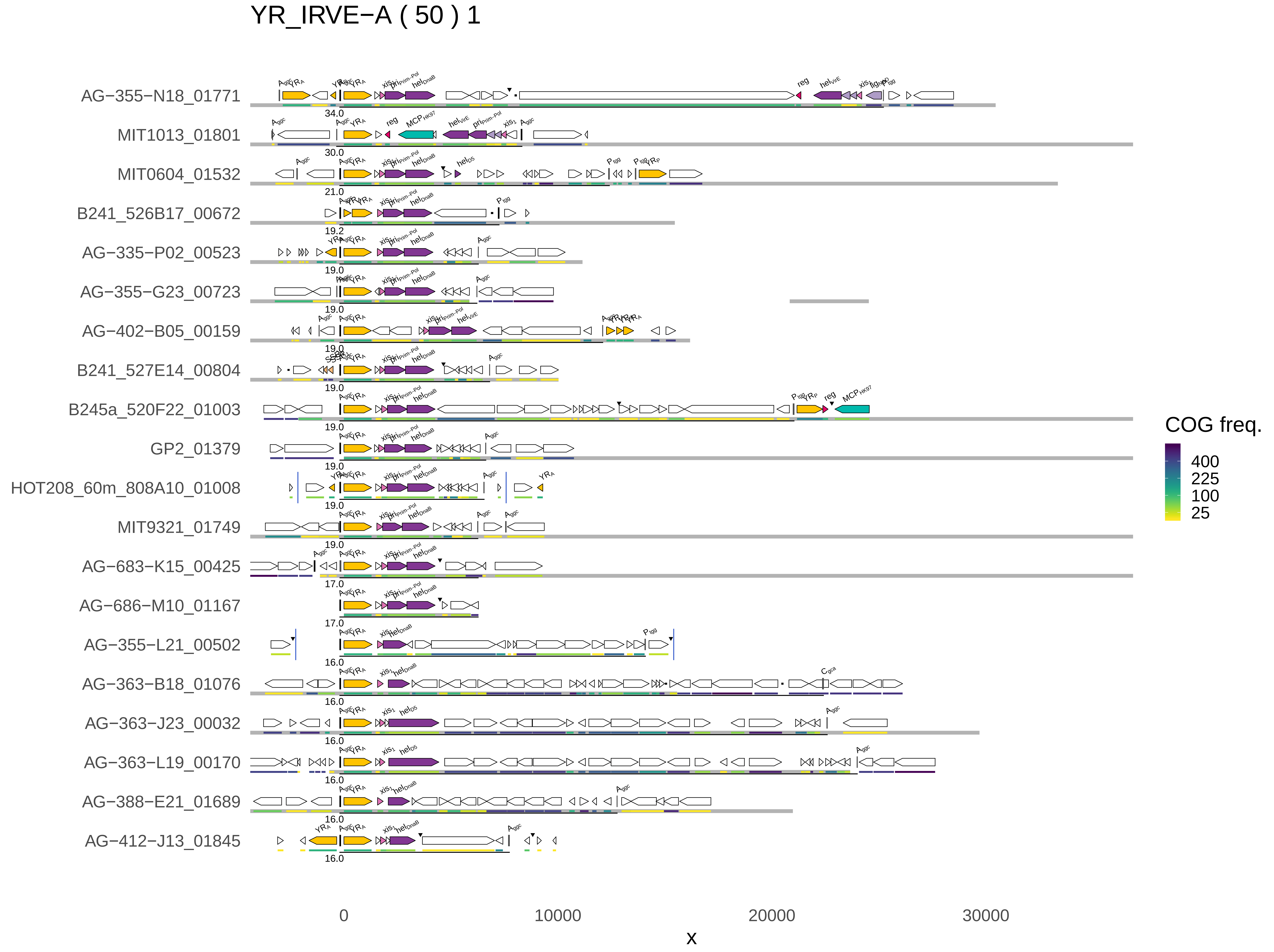
YR_IRVE−A ( 50 ) 1
AG−355−N18_01771
34.0
MIT1013_01801
30.0
MIT0604_01532
21.0
B241_526B17_00672
19.2
AG−335−P02_00523
19.0
AG−355−G23_00723
19.0
AG−402−B05_00159
19.0
B241_527E14_00804
19.0
B245a_520F22_01003
19.0
GP2_01379
19.0
HOT208_60m_808A10_01008
19.0
MIT9321_01749
19.0
AG−683−K15_00425
17.0
AG−686−M10_01167
17.0
AG−355−L21_00502
16.0
AG−363−B18_01076
16.0
AG−363−J23_00032
16.0
AG−363−L19_00170
16.0
AG−388−E21_01689
16.0
AG−412−J13_01845
16.0
0
10000
20000
30000
x
COG freq.
400
225
100
25

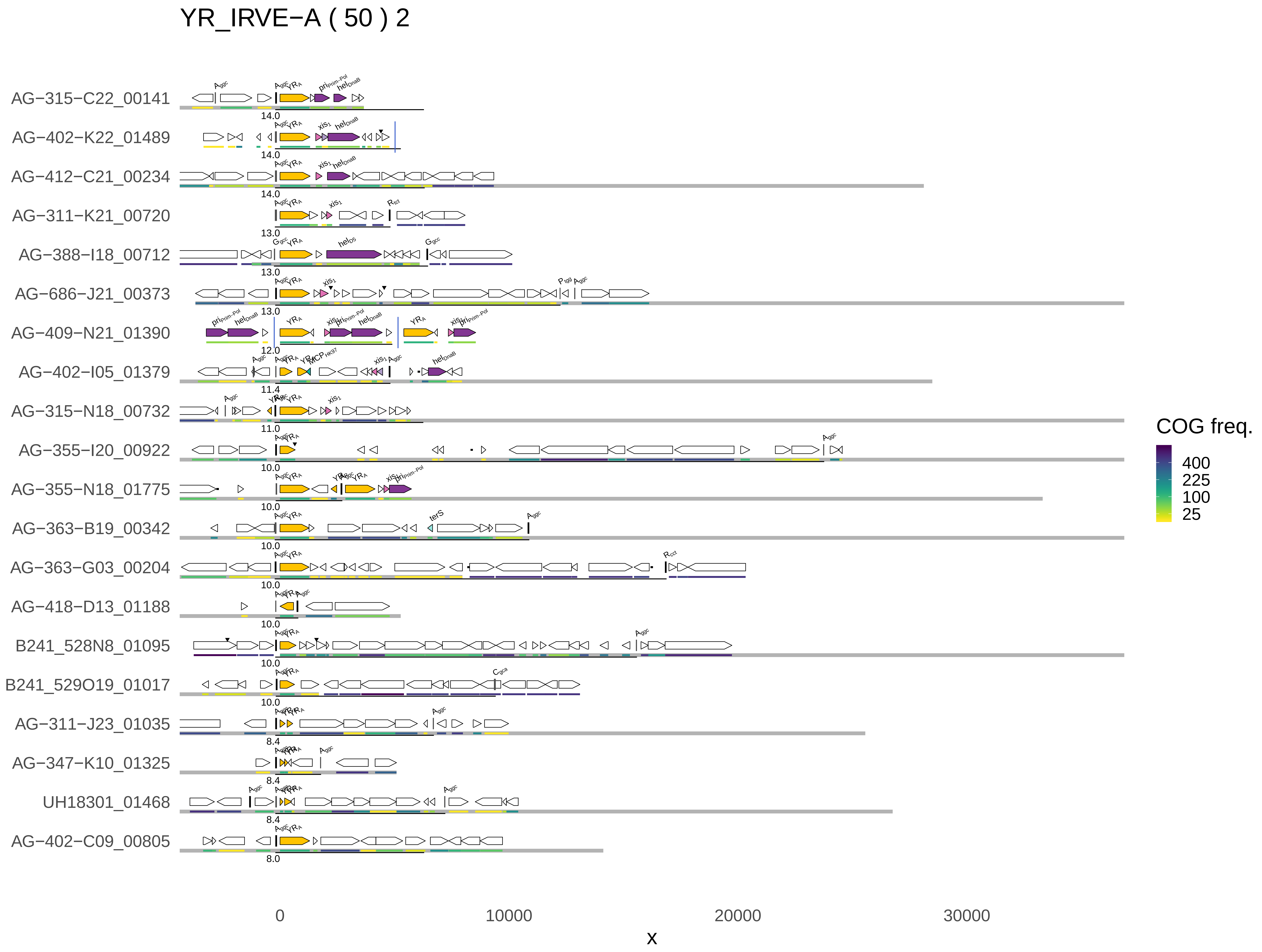
YR_IRVE−A ( 50 ) 2
AG−315−C22_00141
AG−402−K22_01489
AG−412−C21_00234
AG−311−K21_00720
AG−388−I18_00712
AG−686−J21_00373
AG−409−N21_01390
AG−402−I05_01379
AG−315−N18_00732
AG−355−I20_00922
AG−355−N18_01775
AG−363−B19_00342
AG−363−G03_00204
AG−418−D13_01188
B241_528N8_01095
B241_529O19_01017
AG−311−J23_01035
AG−347−K10_01325
UH18301_01468
AG−402−C09_00805
0
10000
20000
30000
x
COG freq.
400
225
100
25

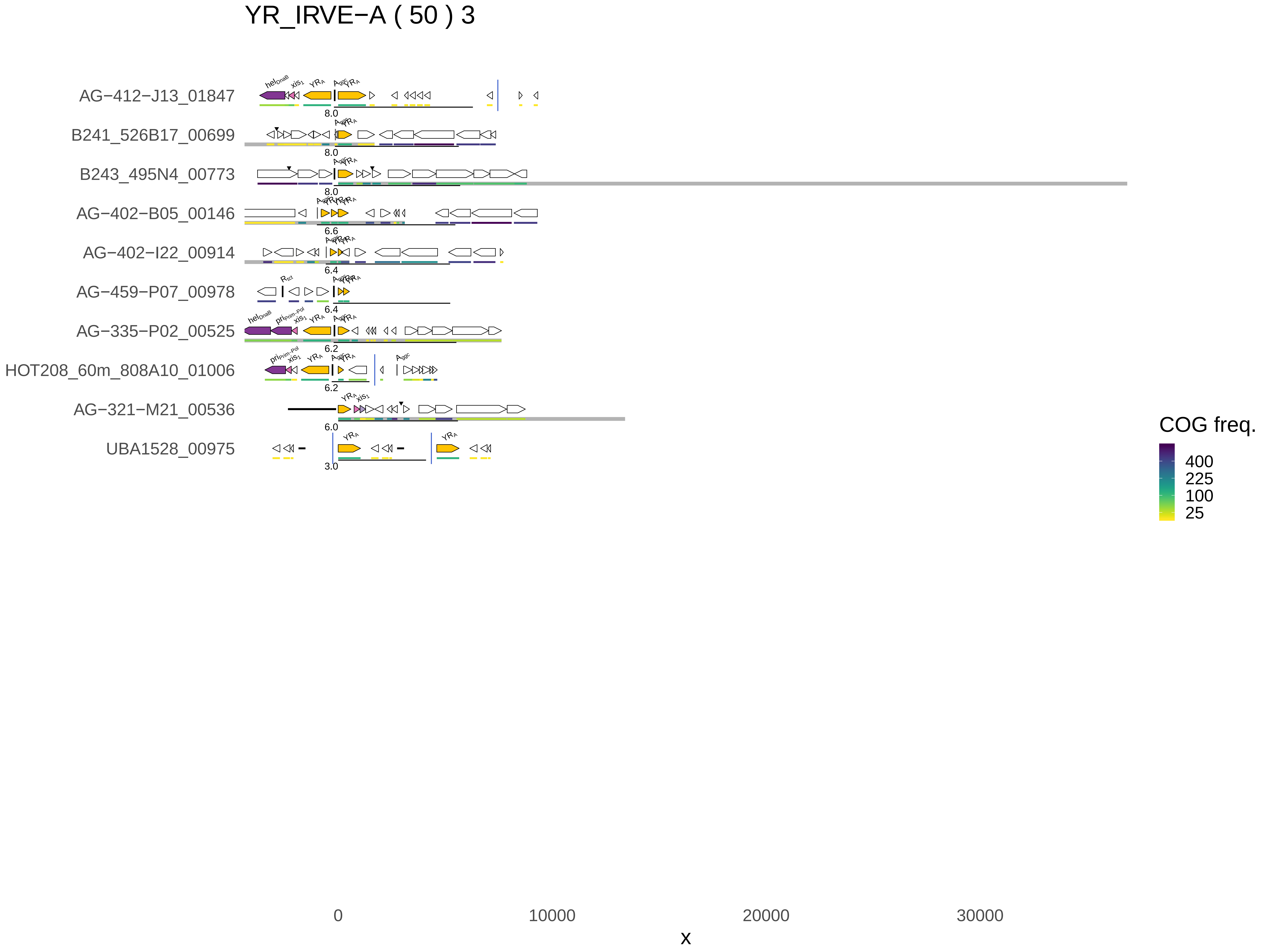
YR_IRVE−A ( 50 ) 3
AG−412−J13_01847
B241_526B17_00699
B243_495N4_00773
AG−402−B05_00146
AG−402−I22_00914
AG−459−P07_00978
AG−335−P02_00525
HOT208_60m_808A10_01006
AG−321−M21_00536
UBA1528_00975
8.0
8.0
8.0
6.6
6.4
6.4
6.2
6.2
6.0
3.0
COG freq.
400
225
100
25
0
10000
20000
30000
x

YR_IRVE−M ( 48 ) 1

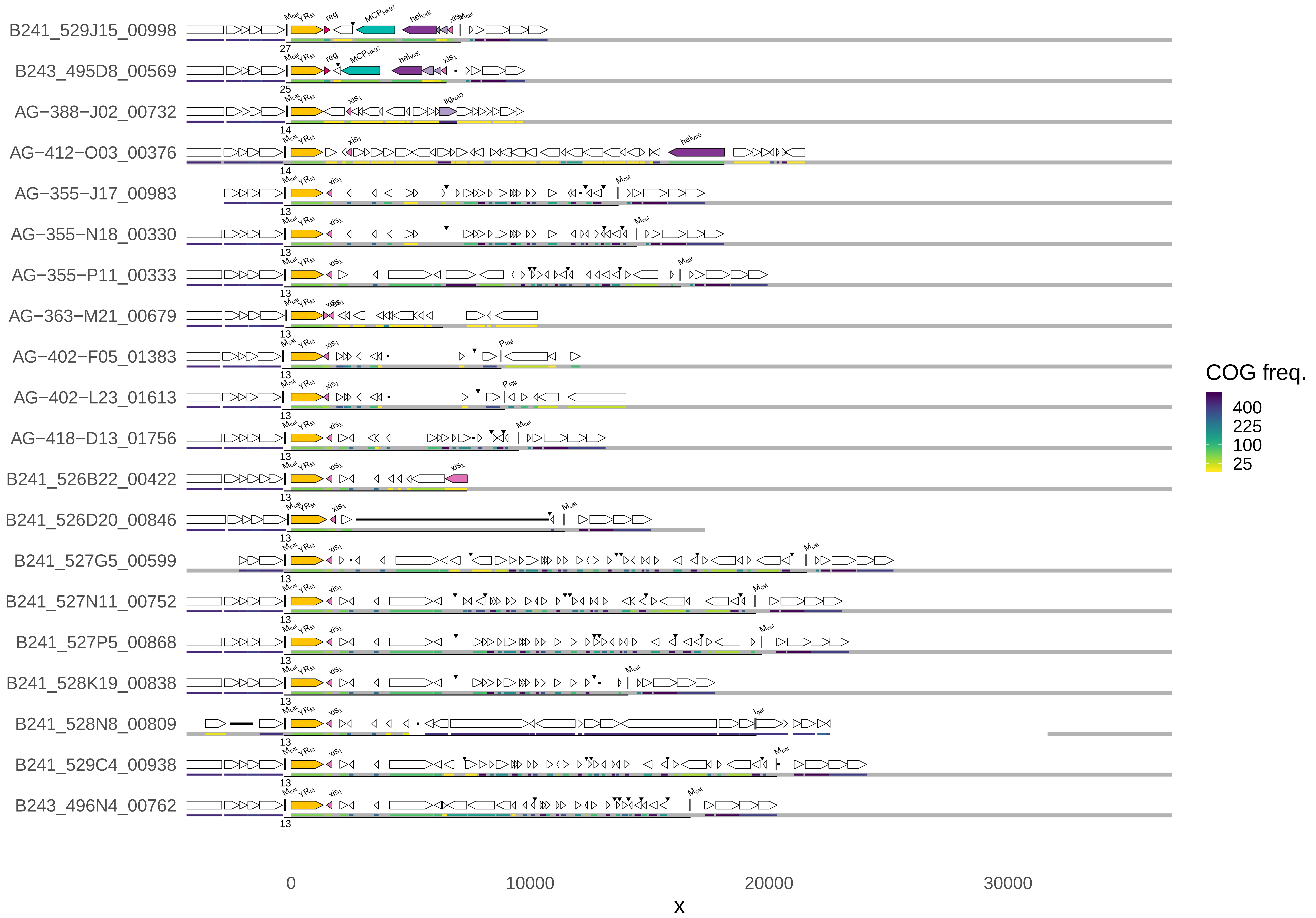
B241_529J15_00998
B243_495D8_00569
AG−388−J02_00732
AG−412−O03_00376
AG−355−J17_00983
AG−355−N18_00330
AG−355−P11_00333
AG−363−M21_00679
AG−402−F05_01383
AG−402−L23_01613
AG−418−D13_01756
B241_526B22_00422
B241_526D20_00846
B241_527G5_00599
B241_527N11_00752
B241_527P5_00868
B241_528K19_00838
B241_528N8_00809
B241_529C4_00938
B243_496N4_00762
0
10000
20000
30000
x
COG freq.
400
225
100
25

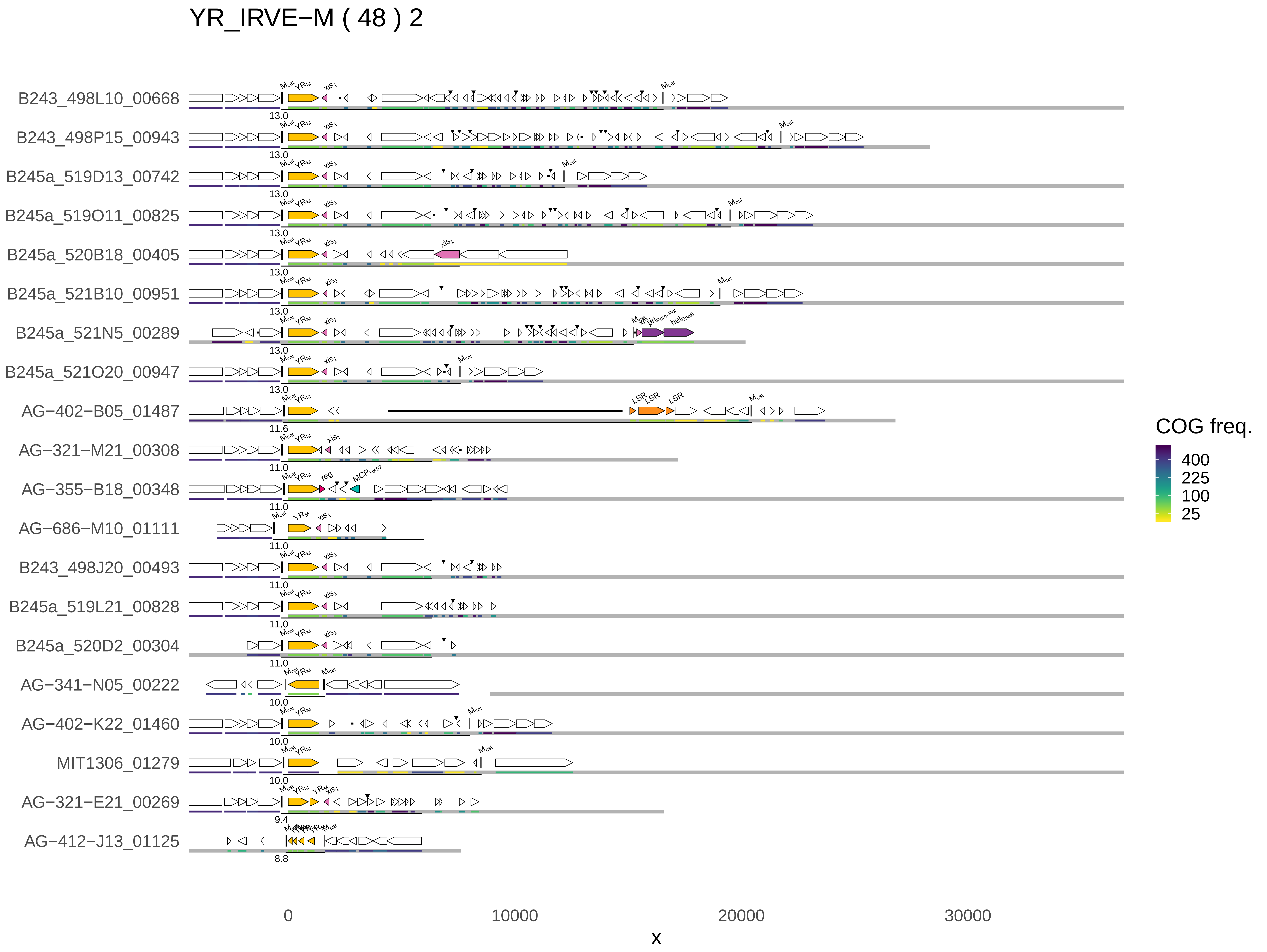

YR_IRVE−M ( 48 ) 2
B243_498L10_00668
B243_498P15_00943
B245a_519D13_00742
B245a_519O11_00825
B245a_520B18_00405
B245a_521B10_00951
B245a_521N5_00289
B245a_521O20_00947
AG−402−B05_01487
AG−321−M21_00308
AG−355−B18_00348
AG−686−M10_01111
B243_498J20_00493
B245a_519L21_00828
B245a_520D2_00304
AG−341−N05_00222
AG−402−K22_01460
MIT1306_01279
AG−321−E21_00269
AG−412−J13_01125
13.0
11.6
11.0
10.0
9.4
8.8
xis1
reg
LSR
0
10000
20000
30000
x
COG freq.
400
225
100
25

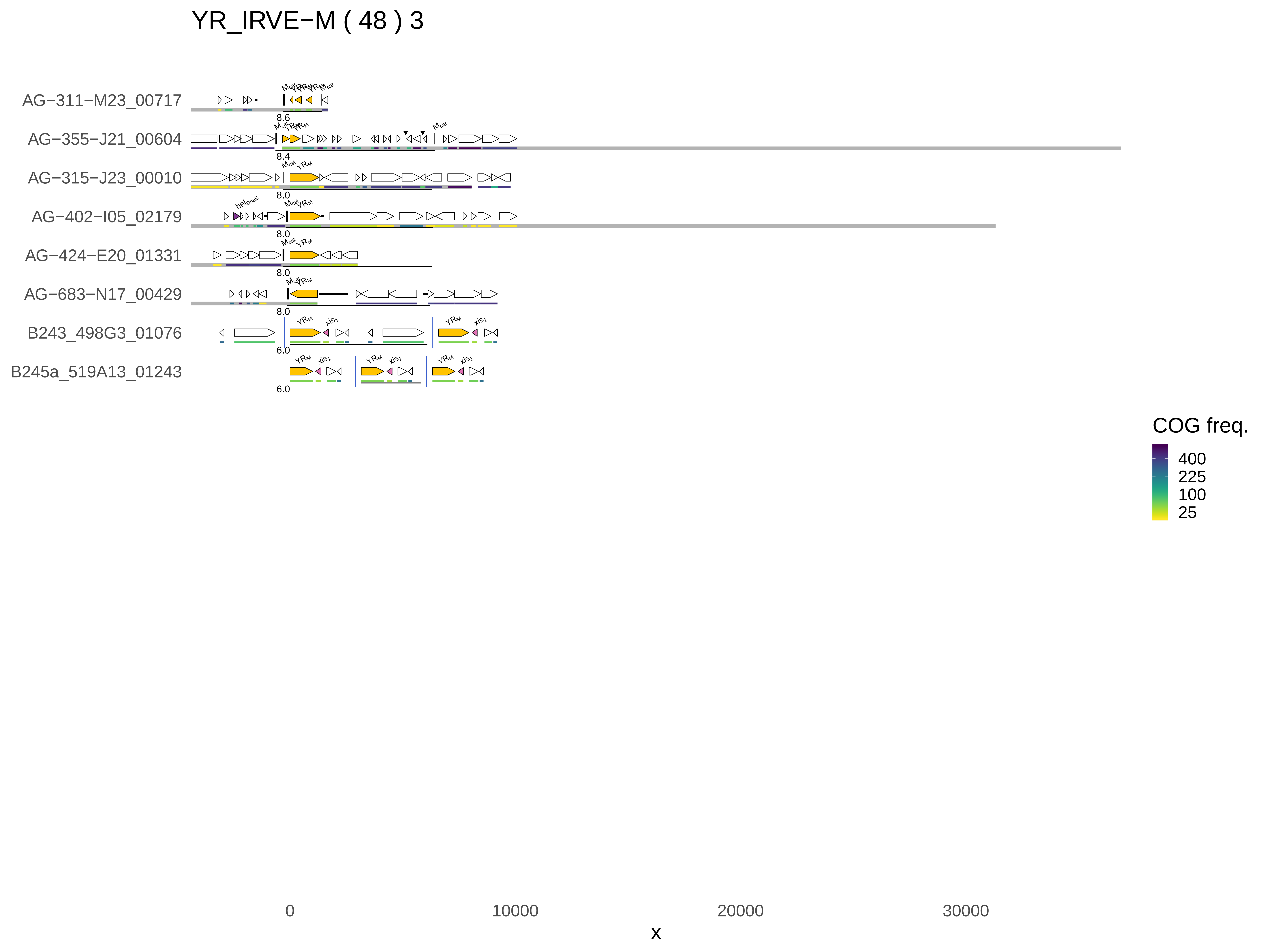

YR_IRVE−M ( 48 ) 3
AG−311−M23_00717
AG−355−J21_00604
AG−315−J23_00010
AG−402−I05_02179
AG−424−E20_01331
AG−683−N17_00429
B243_498G3_01076
B245a_519A13_01243
8.6
8.4
8.0
6.0
COG freq.
400
225
100
25
0
10000
20000
30000
x

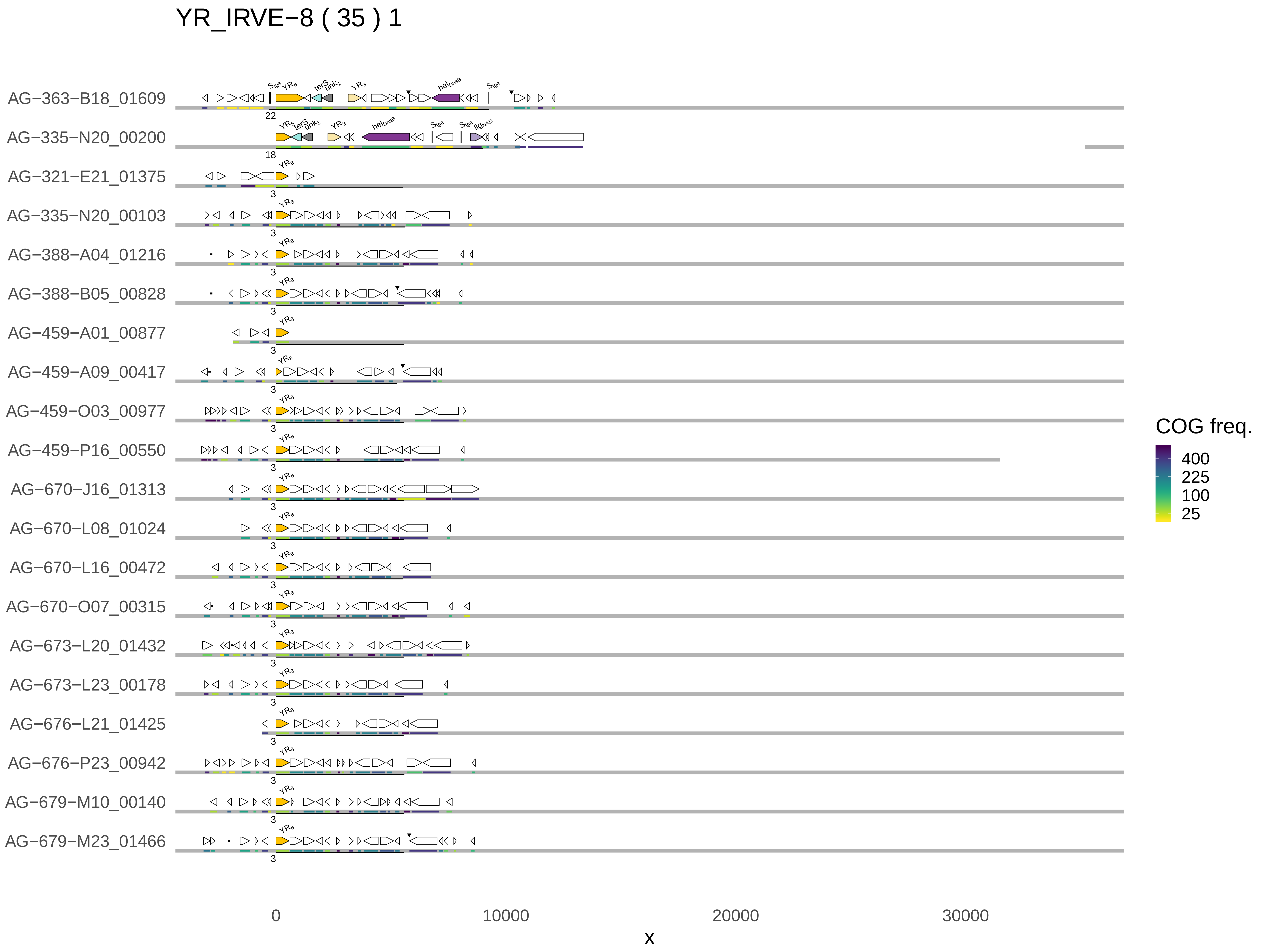

YR_IRVE−8 ( 35 ) 1
AG−363−B18_01609
AG−335−N20_00200
AG−321−E21_01375
AG−335−N20_00103
AG−388−A04_01216
AG−388−B05_00828
AG−459−A01_00877
AG−459−A09_00417
AG−459−O03_00977
AG−459−P16_00550
AG−670−J16_01313
AG−670−L08_01024
AG−670−L16_00472
AG−670−O07_00315
AG−673−L20_01432
AG−673−L23_00178
AG−676−L21_01425
AG−676−P23_00942
AG−679−M10_00140
AG−679−M23_01466
0
10000
20000
30000
x
COG freq.
400
225
100
25

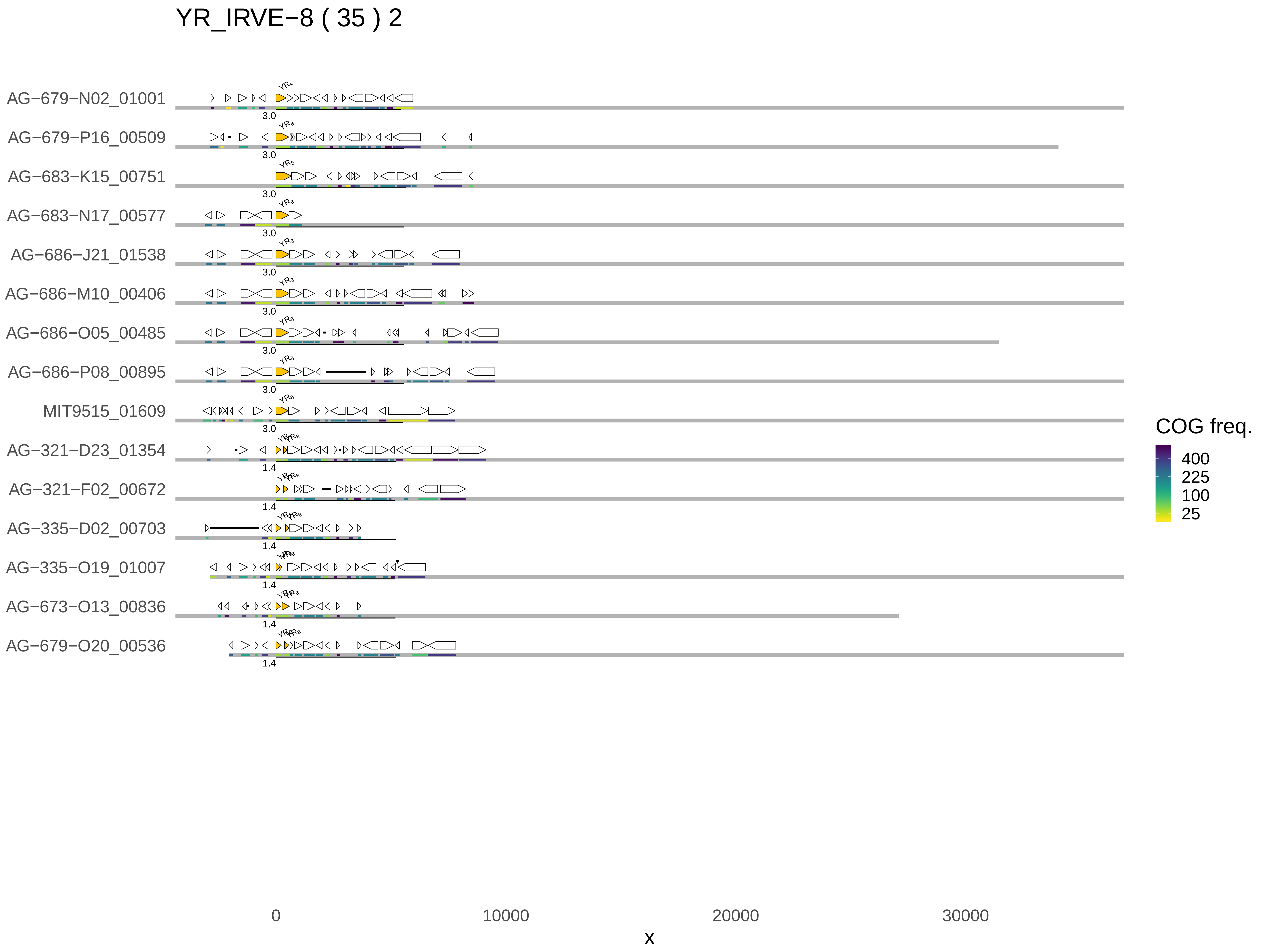

YR_IRVE−8 ( 35 ) 2
AG−679−N02_01001
AG−679−P16_00509
AG−683−K15_00751
AG−683−N17_00577
AG−686−J21_01538
AG−686−M10_00406
AG−686−O05_00485
AG−686−P08_00895
MIT9515_01609
AG−321−D23_01354
AG−321−F02_00672
AG−335−D02_00703
AG−335−O19_01007
AG−673−O13_00836
AG−679−O20_00536
3.0
3.0
3.0
3.0
3.0
3.0
3.0
3.0
3.0
1.4
1.4
1.4
1.4
1.4
1.4
0
10000
20000
30000
x
COG freq.
400
225
100
25

YR_IRVE−T1 ( 33 ) 1

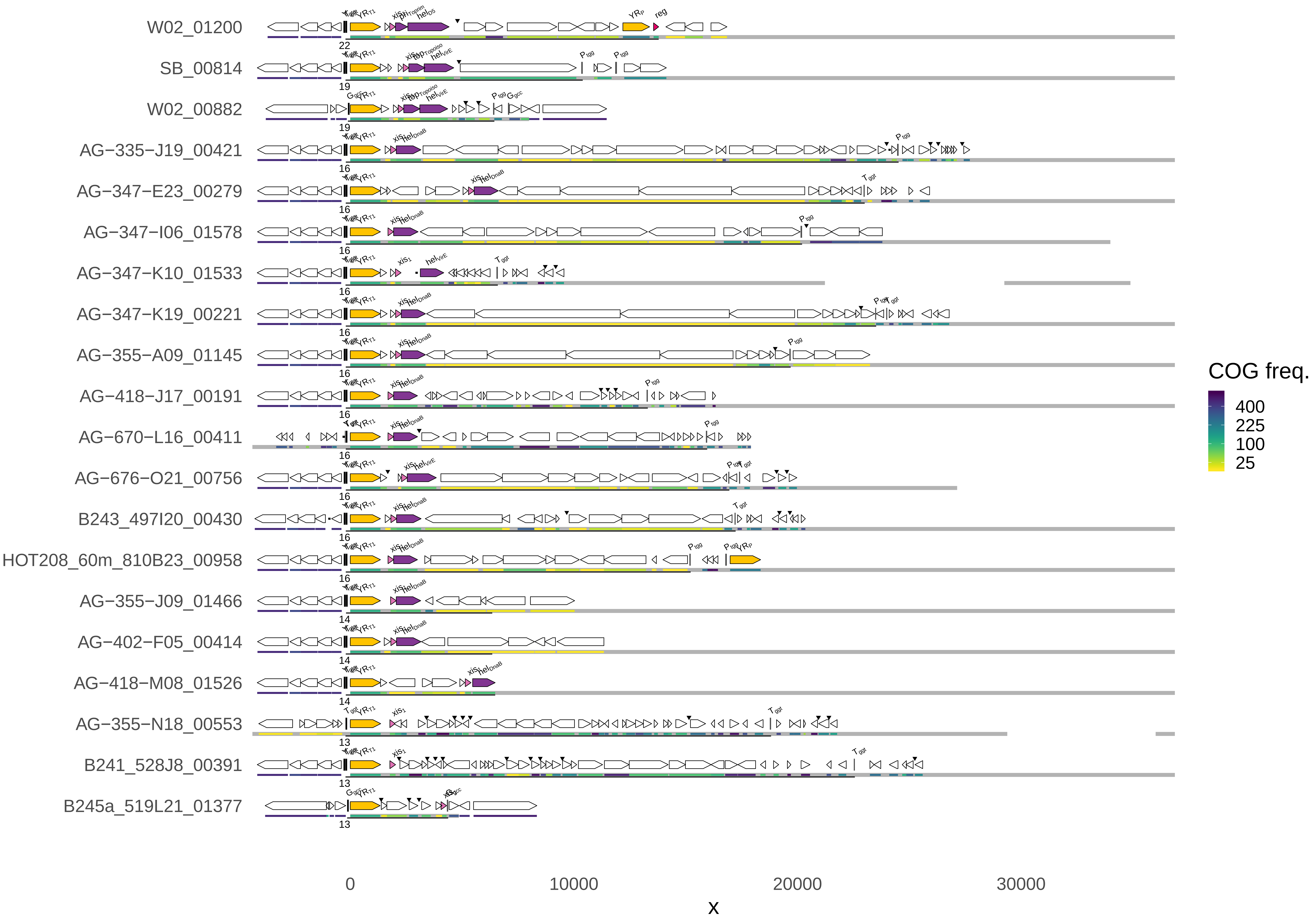

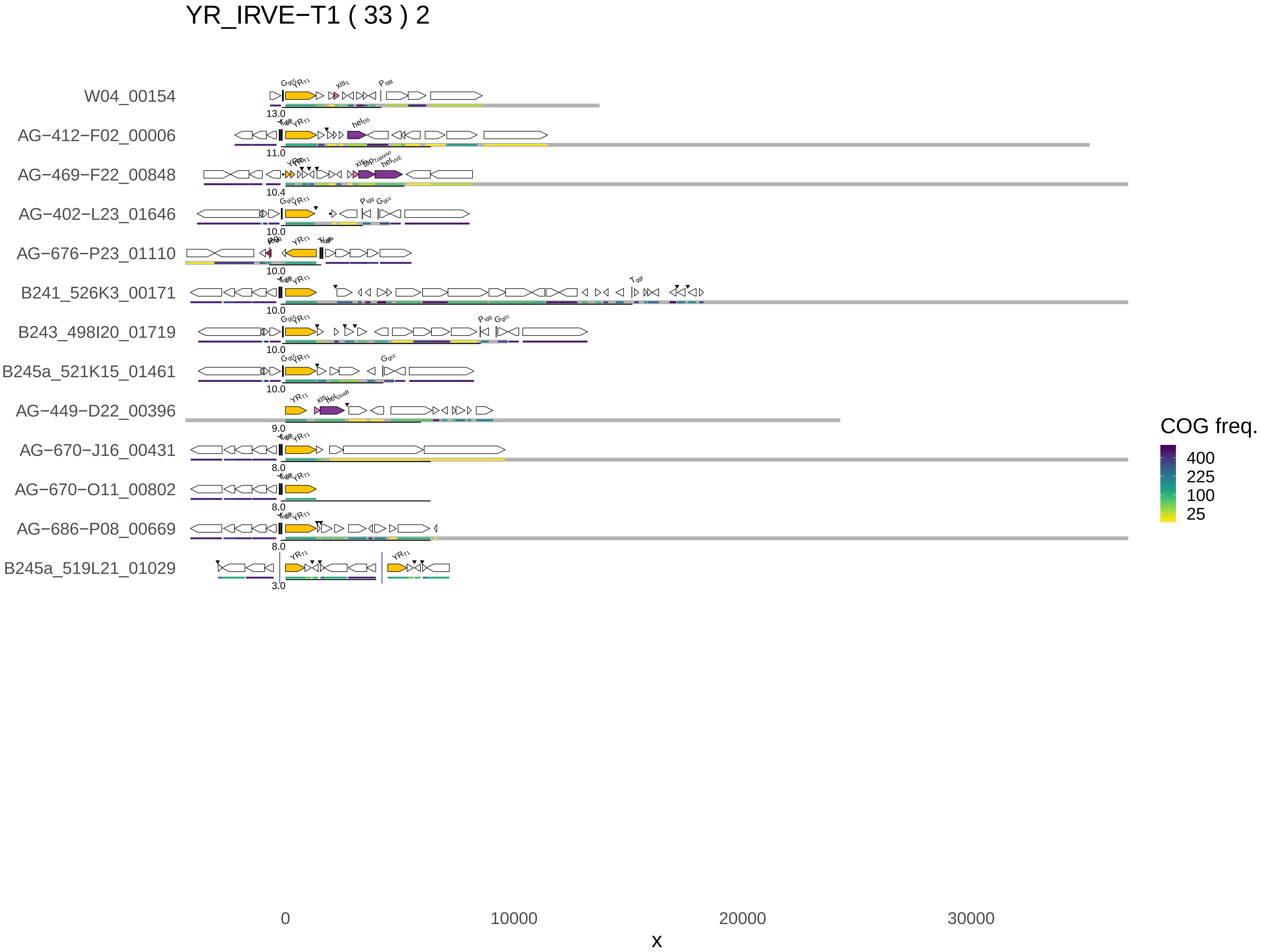
YR_IRVE−T1 ( 33 ) 2
W04_00154
AG−412−F02_00006
AG−469−F22_00848
AG−402−L23_01646
AG−676−P23_01110
B241_526K3_00171
B243_498I20_01719
B245a_521K15_01461
AG−449−D22_00396
AG−670−J16_00431
AG−670−O11_00802
AG−686−P08_00669
B245a_519L21_01029
0
10000
20000
30000
x
COG freq.
400
225
100
25

LSR_IRVE ( 27 ) 1

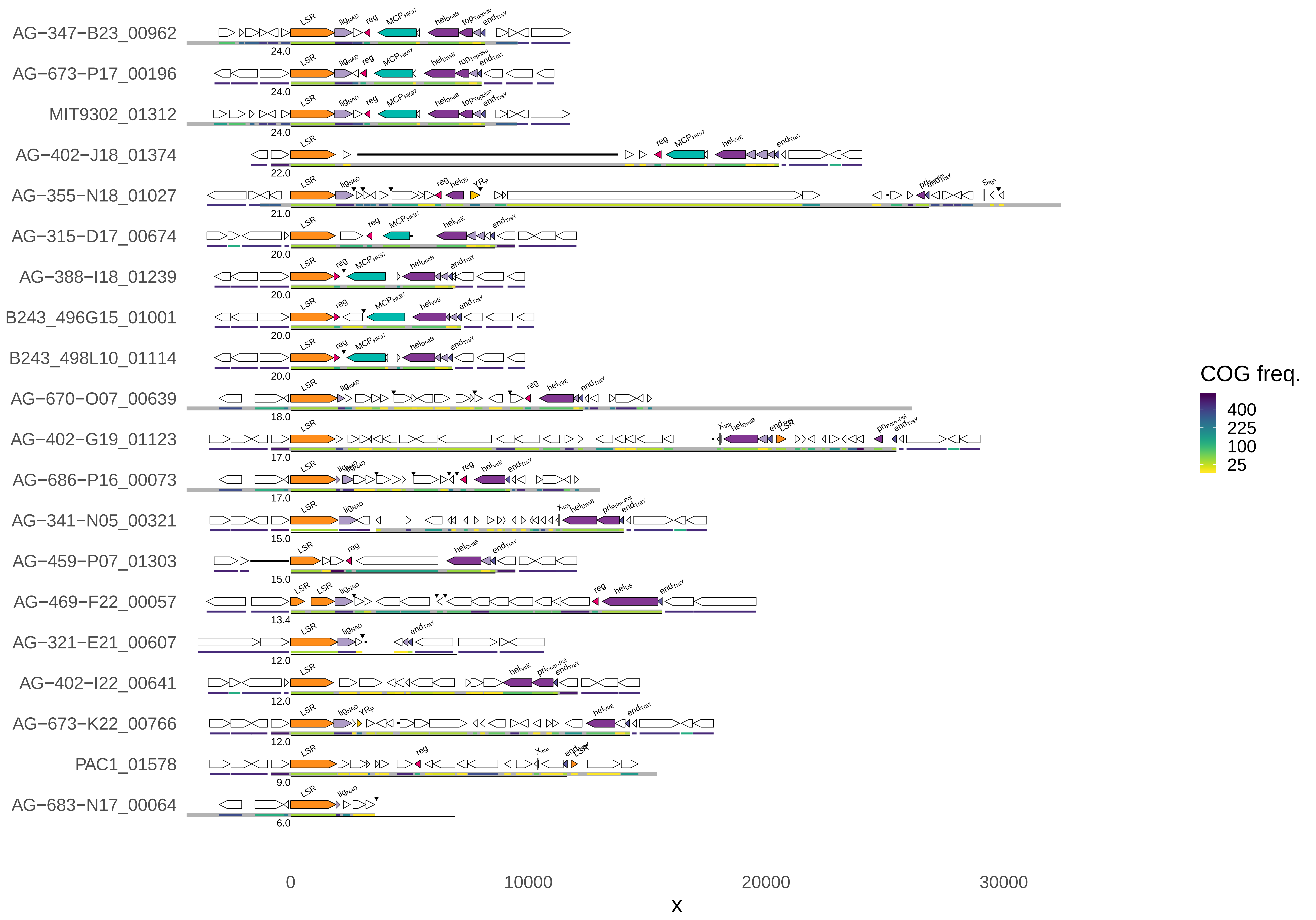
AG−347−B23_00962
AG−673−P17_00196
MIT9302_01312
AG−402−J18_01374
AG−355−N18_01027
AG−315−D17_00674
AG−388−I18_01239
B243_496G15_01001
B243_498L10_01114
AG−670−O07_00639
AG−402−G19_01123
AG−686−P16_00073
AG−341−N05_00321
AG−459−P07_01303
AG−469−F22_00057
AG−321−E21_00607
AG−402−I22_00641
AG−673−K22_00766
PAC1_01578
AG−683−N17_00064
24.0
24.0
24.0
22.0
21.0
20.0
20.0
20.0
20.0
18.0
17.0
17.0
15.0
15.0
13.4
12.0
12.0
12.0
9.0
6.0
0
10000
20000
30000
x
COG freq.
400
225
100
25

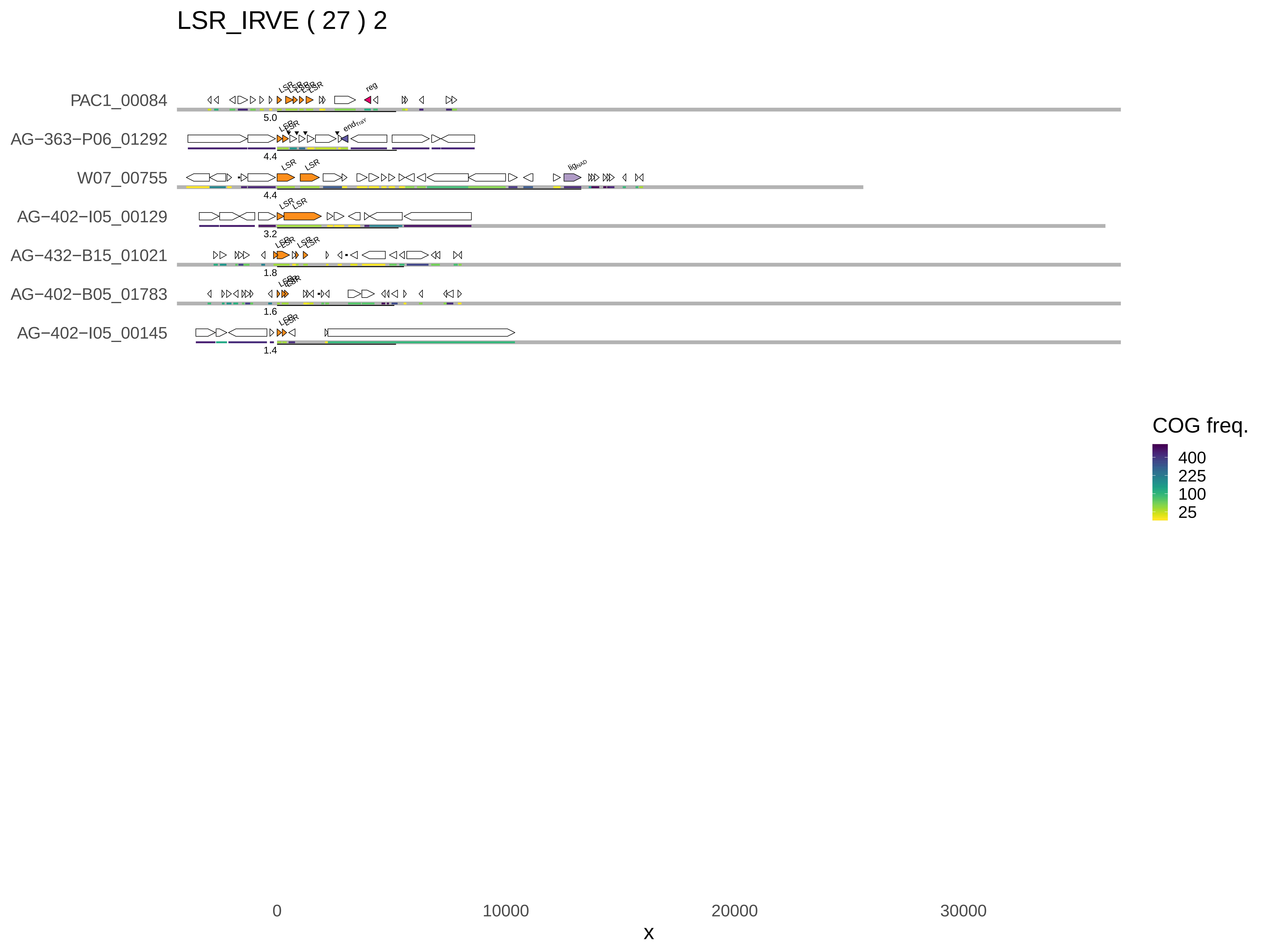
LSR_IRVE ( 27 ) 2
PAC1_00084
AG−363−P06_01292
W07_00755
AG−402−I05_00129
AG−432−B15_01021
AG−402−B05_01783
AG−402−I05_00145
5.0
4.4
4.4
3.2
1.8
1.6
1.4
reg
end
lig
COG freq.
400
225
100
25
0
10000
20000
30000
x

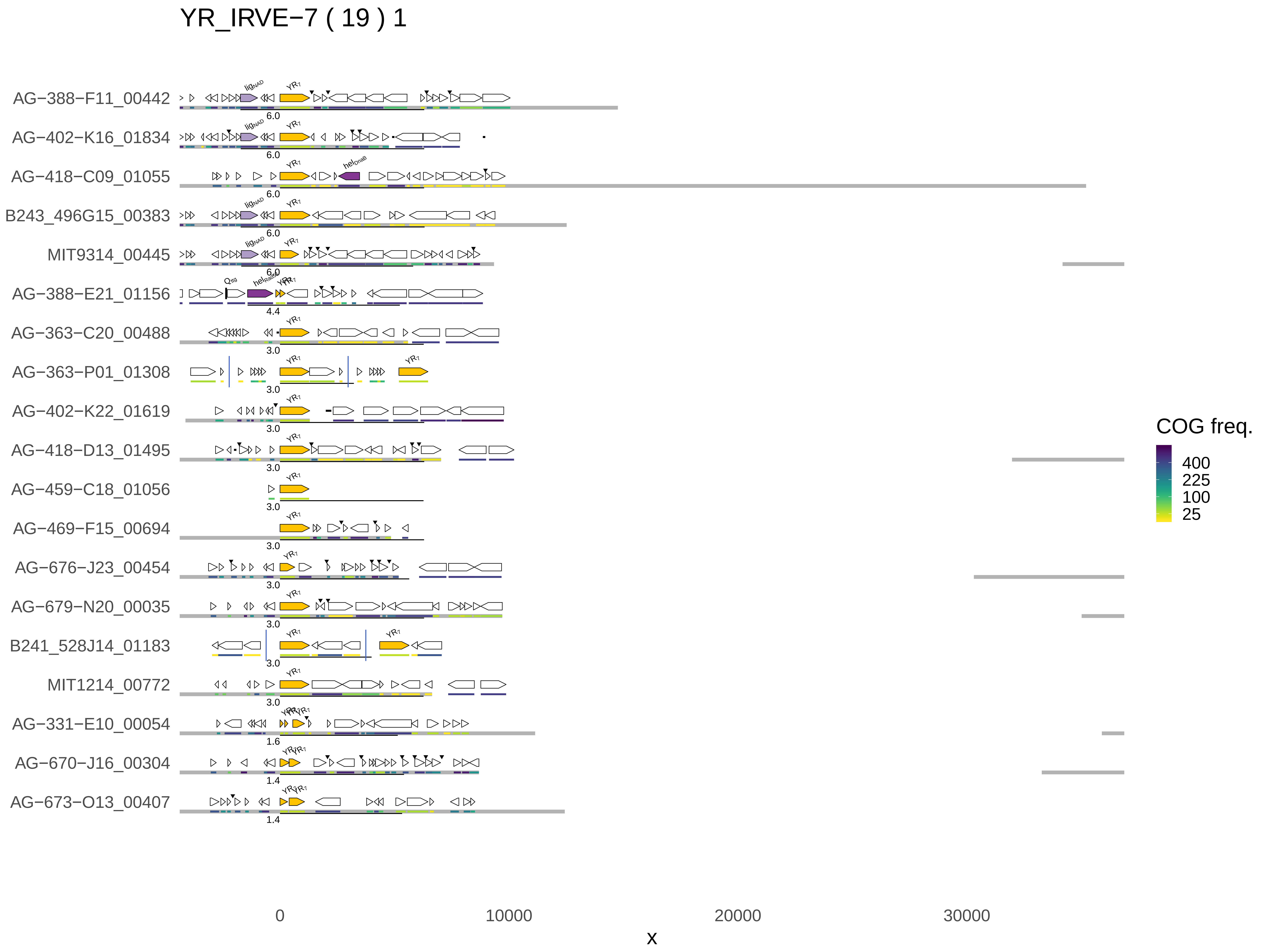
YR_IRVE−7 ( 19 ) 1
AG−388−F11_00442
AG−402−K16_01834
AG−418−C09_01055
B243_496G15_00383
MIT9314_00445
AG−388−E21_01156
AG−363−C20_00488
AG−363−P01_01308
AG−402−K22_01619
AG−418−D13_01495
AG−459−C18_01056
AG−469−F15_00694
AG−676−J23_00454
AG−679−N20_00035
B241_528J14_01183
MIT1214_00772
AG−331−E10_00054
AG−670−J16_00304
AG−673−O13_00407
6.0
6.0
6.0
6.0
6.0
4.4
3.0
3.0
3.0
3.0
3.0
3.0
3.0
3.0
3.0
3.0
1.6
1.4
1.4
COG freq.
400
225
100
25
0
10000
20000
30000
x

### YR_IRVE−9 ( 15 ) 1

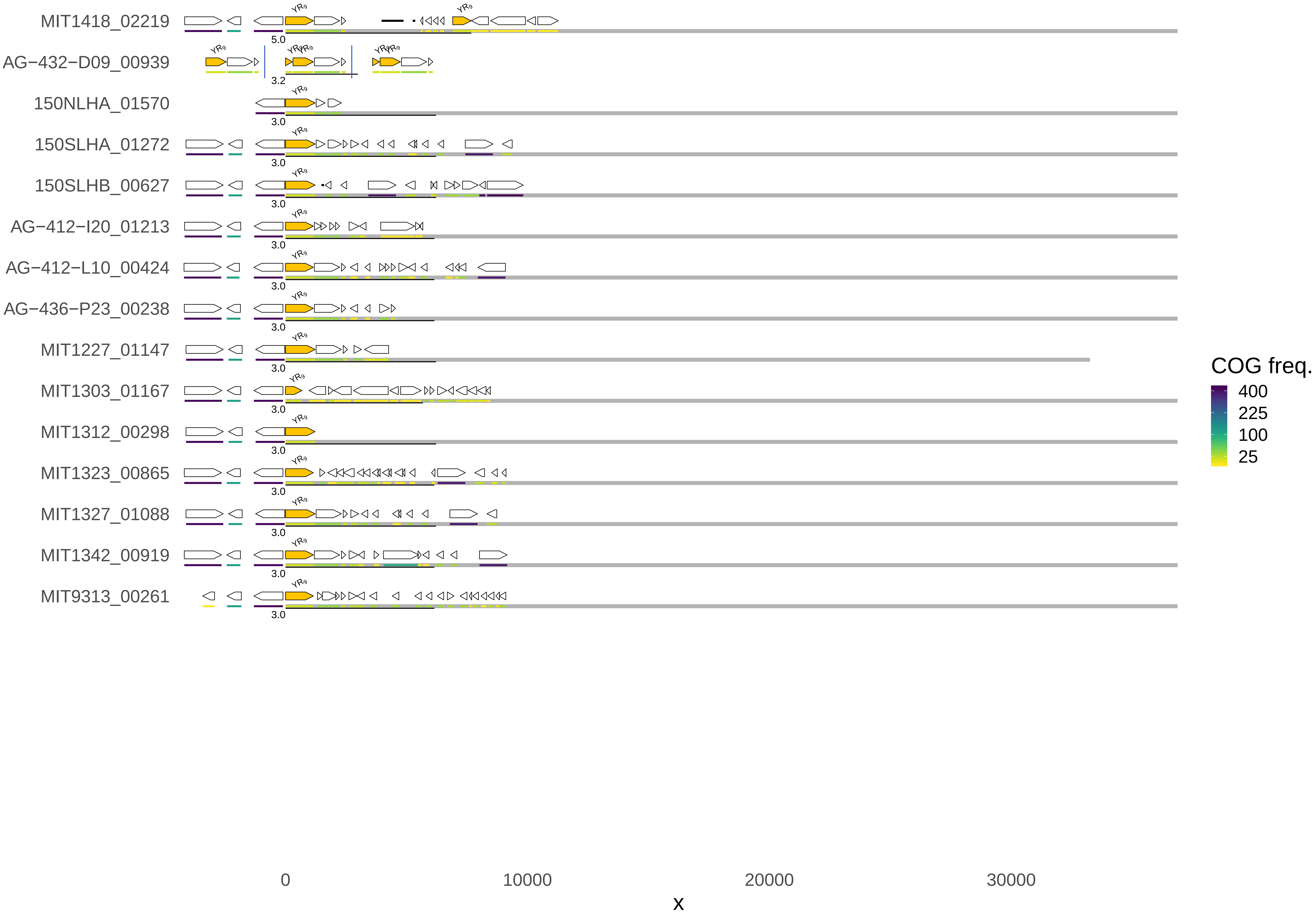
MIT1418_02219
AG−432−D09_00939
150NLHA_01570
150SLHA_01272
150SLHB_00627
AG−412−I20_01213
AG−412−L10_00424
AG−436−P23_00238
MIT1227_01147
MIT1303_01167
MIT1312_00298
MIT1323_00865
MIT1327_01088
MIT1342_00919
MIT9313_00261
YR9
5.0
3.2
3.0
0
10000
20000
30000
x
COG freq.
400
225
100
25

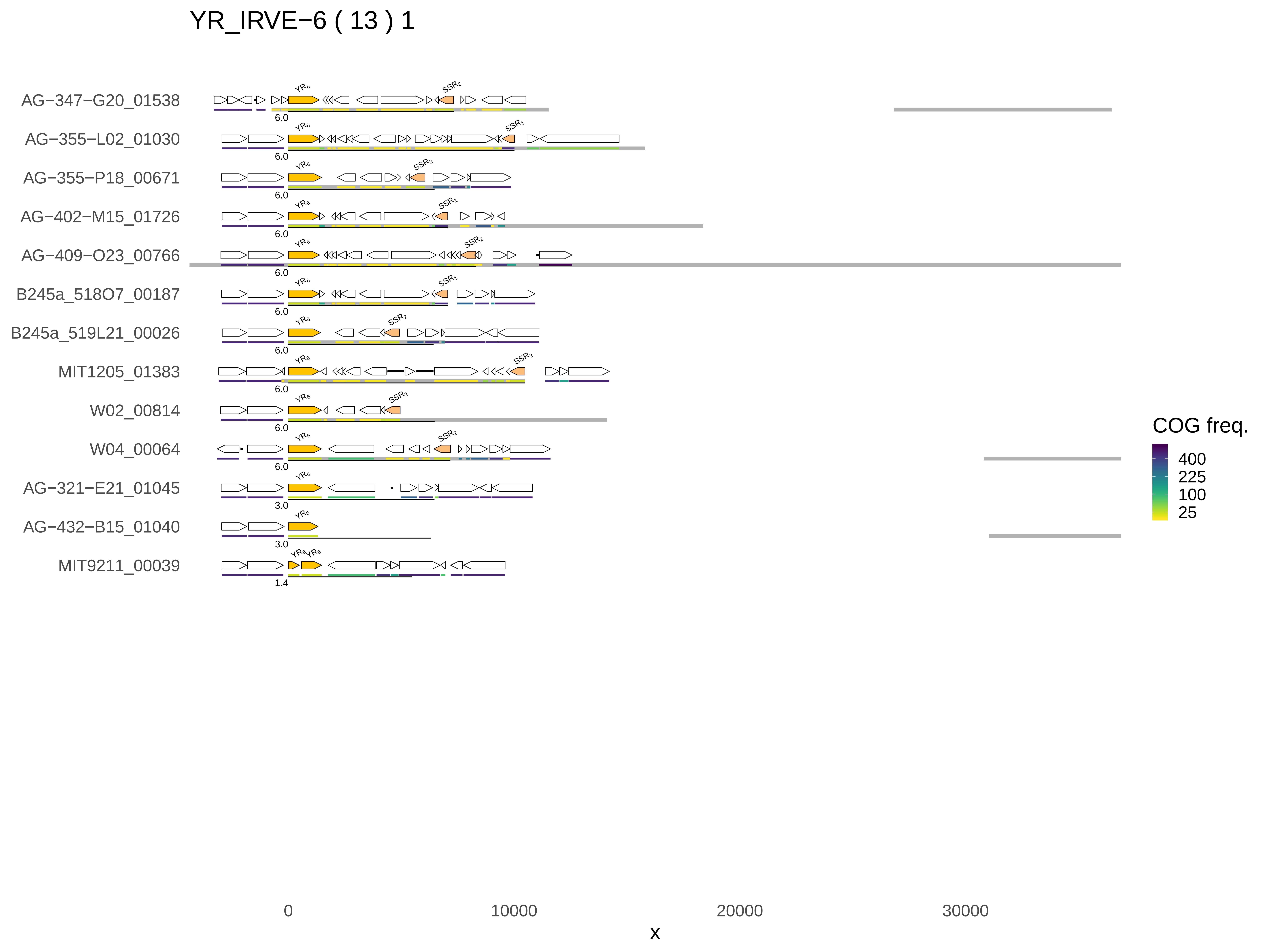
YR_IRVE−6 ( 13 ) 1
AG−347−G20_01538
AG−355−L02_01030
AG−355−P18_00671
AG−402−M15_01726
AG−409−O23_00766
B245a_518O7_00187
B245a_519L21_00026
MIT1205_01383
W02_00814
W04_00064
AG−321−E21_01045
AG−432−B15_01040
MIT9211_00039
YR6
SSR2
SSR1
6.0
3.0
1.4
0
10000
20000
30000
x
COG freq.
400
225
100
25

YR_IRVE−R ( 12 ) 1

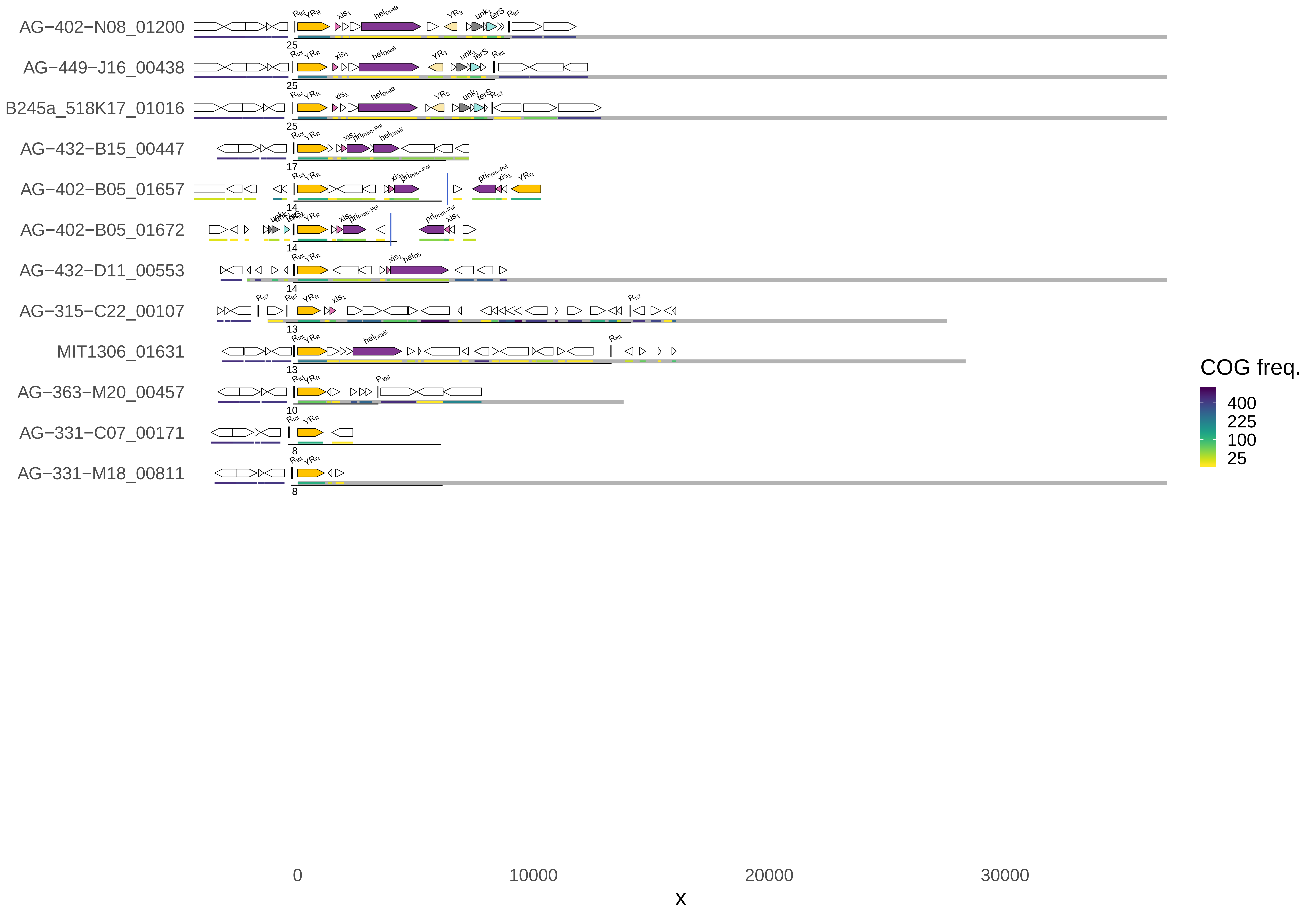
AG−402−N08_01200
AG−449−J16_00438
B245a_518K17_01016
AG−432−B15_00447
AG−402−B05_01657
AG−402−B05_01672
AG−432−D11_00553
AG−315−C22_00107
MIT1306_01631
AG−363−M20_00457
AG−331−C07_00171
AG−331−M18_00811
25
25
25
17
14
14
14
13
13
10
8
8
0
10000
20000
30000
x
COG freq.
400
225
100
25

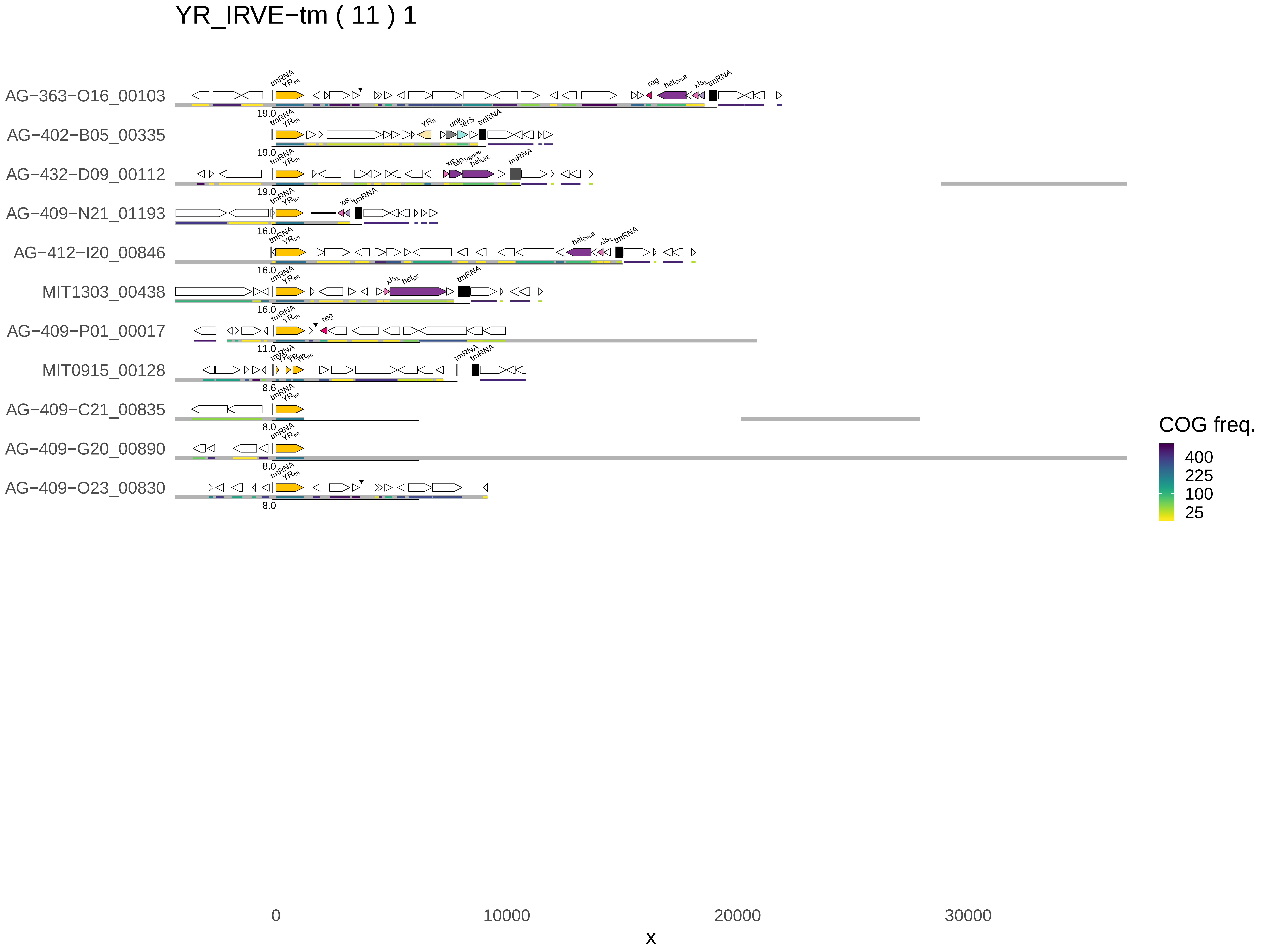
YR_IRVE−tm ( 11 ) 1
AG−363−O16_00103
AG−402−B05_00335
AG−432−D09_00112
AG−409−N21_01193
AG−412−I20_00846
MIT1303_00438
AG−409−P01_00017
MIT0915_00128
AG−409−C21_00835
AG−409−G20_00890
AG−409−O23_00830
tmRNA
reg
19.0
16.0
11.0
8.6
8.0
0
10000
20000
30000
x
COG freq.
400
225
100
25

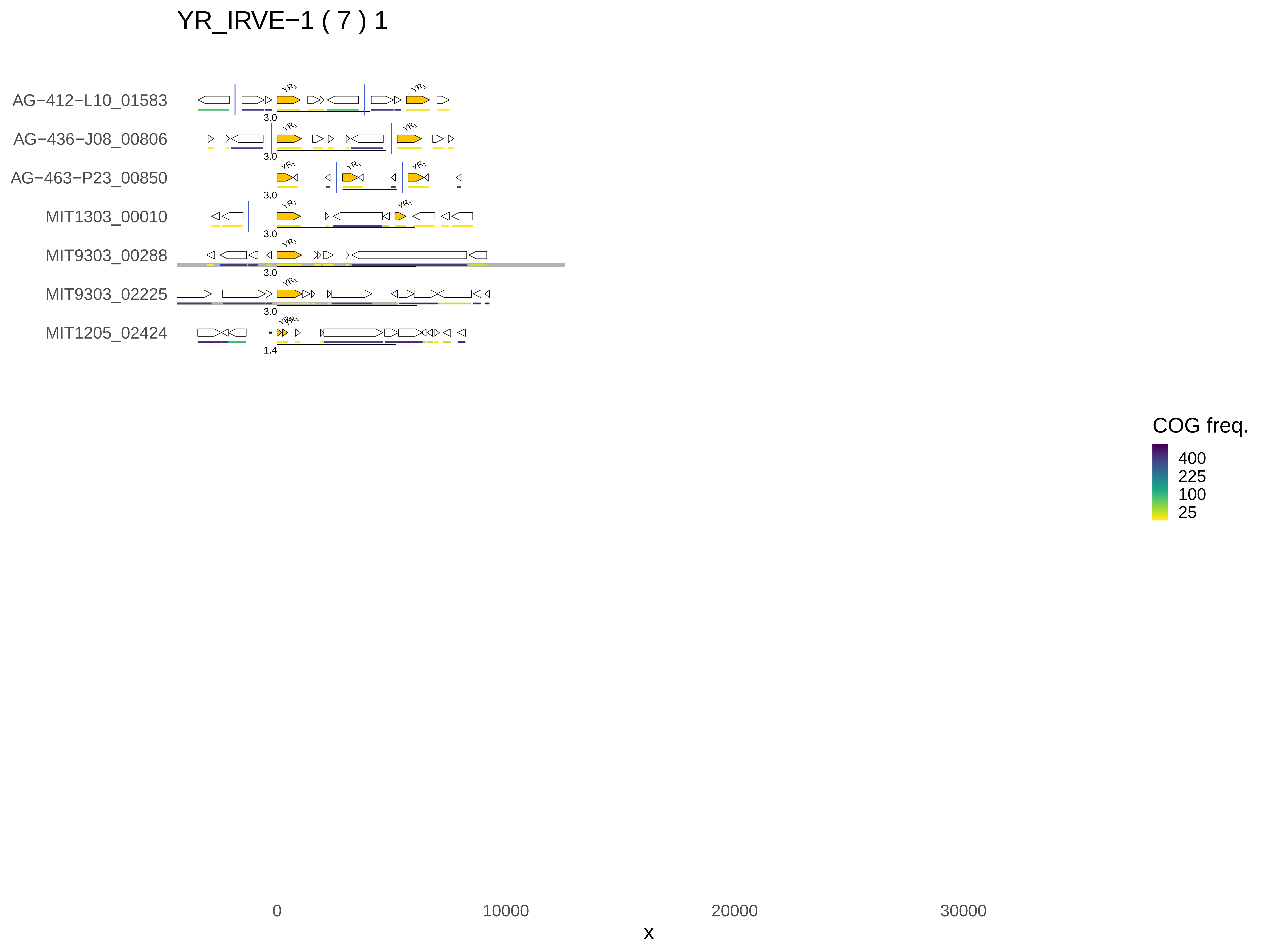

YR_IRVE−1 ( 7 ) 1
AG−412−L10_01583
AG−436−J08_00806
AG−463−P23_00850
MIT1303_00010
MIT9303_00288
MIT9303_02225
MIT1205_02424
YR1
3.0
1.4
0
10000
20000
30000
x
COG freq.
400
225
100
25

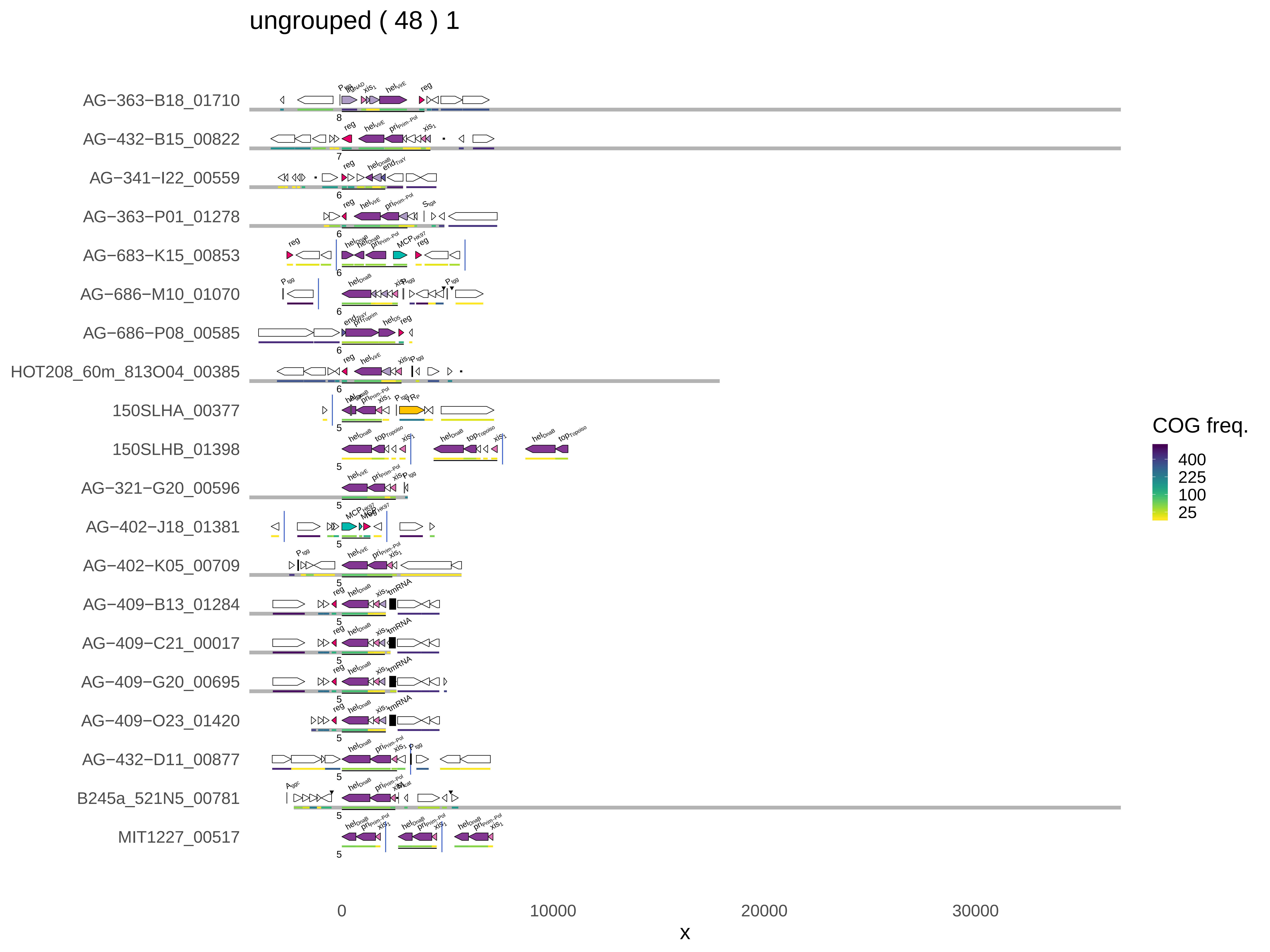
ungrouped ( 48 ) 1
AG−363−B18_01710
AG−432−B15_00822
AG−341−I22_00559
AG−363−P01_01278
AG−683−K15_00853
AG−686−M10_01070
AG−686−P08_00585
HOT208_60m_813O04_00385
150SLHA_00377
150SLHB_01398
AG−321−G20_00596
AG−402−J18_01381
AG−402−K05_00709
AG−409−B13_01284
AG−409−C21_00017
AG−409−G20_00695
AG−409−O23_01420
AG−432−D11_00877
B245a_521N5_00781
MIT1227_00517
reg
hel
pri
xis
tmRNA
MCP
top
0
10000
20000
30000
x
COG freq.
400
225
100
25

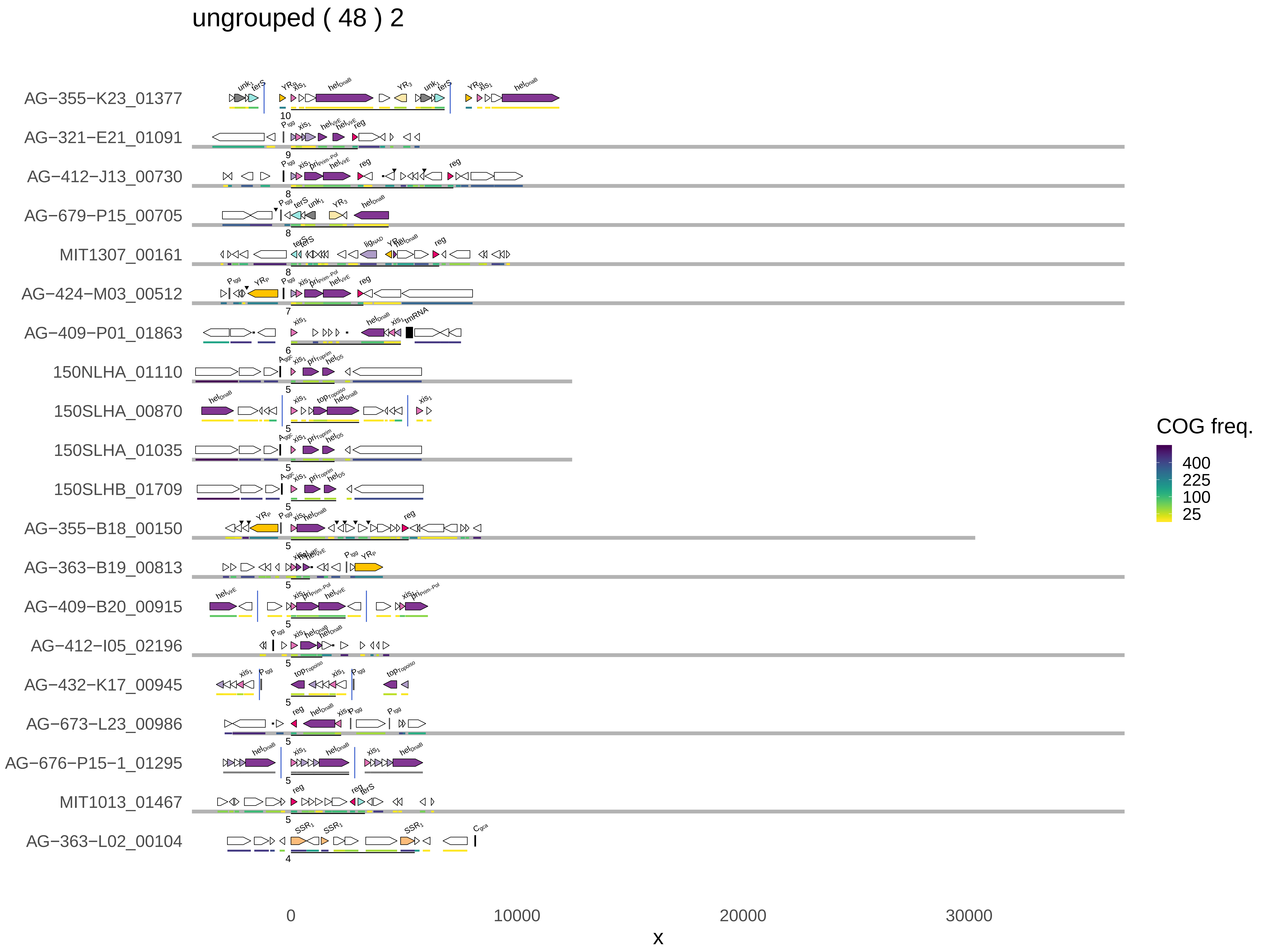
ungrouped ( 48 ) 2
AG−355−K23_01377
AG−321−E21_01091
AG−412−J13_00730
AG−679−P15_00705
MIT1307_00161
AG−424−M03_00512
AG−409−P01_01863
150NLHA_01110
150SLHA_00870
150SLHA_01035
150SLHB_01709
AG−355−B18_00150
AG−363−B19_00813
AG−409−B20_00915
AG−412−I05_02196
AG−432−K17_00945
AG−673−L23_00986
AG−676−P15−1_01295
MIT1013_01467
AG−363−L02_00104
0
10000
20000
30000
x
COG freq.
400
225
100
25

### ungrouped ( 48 ) 3

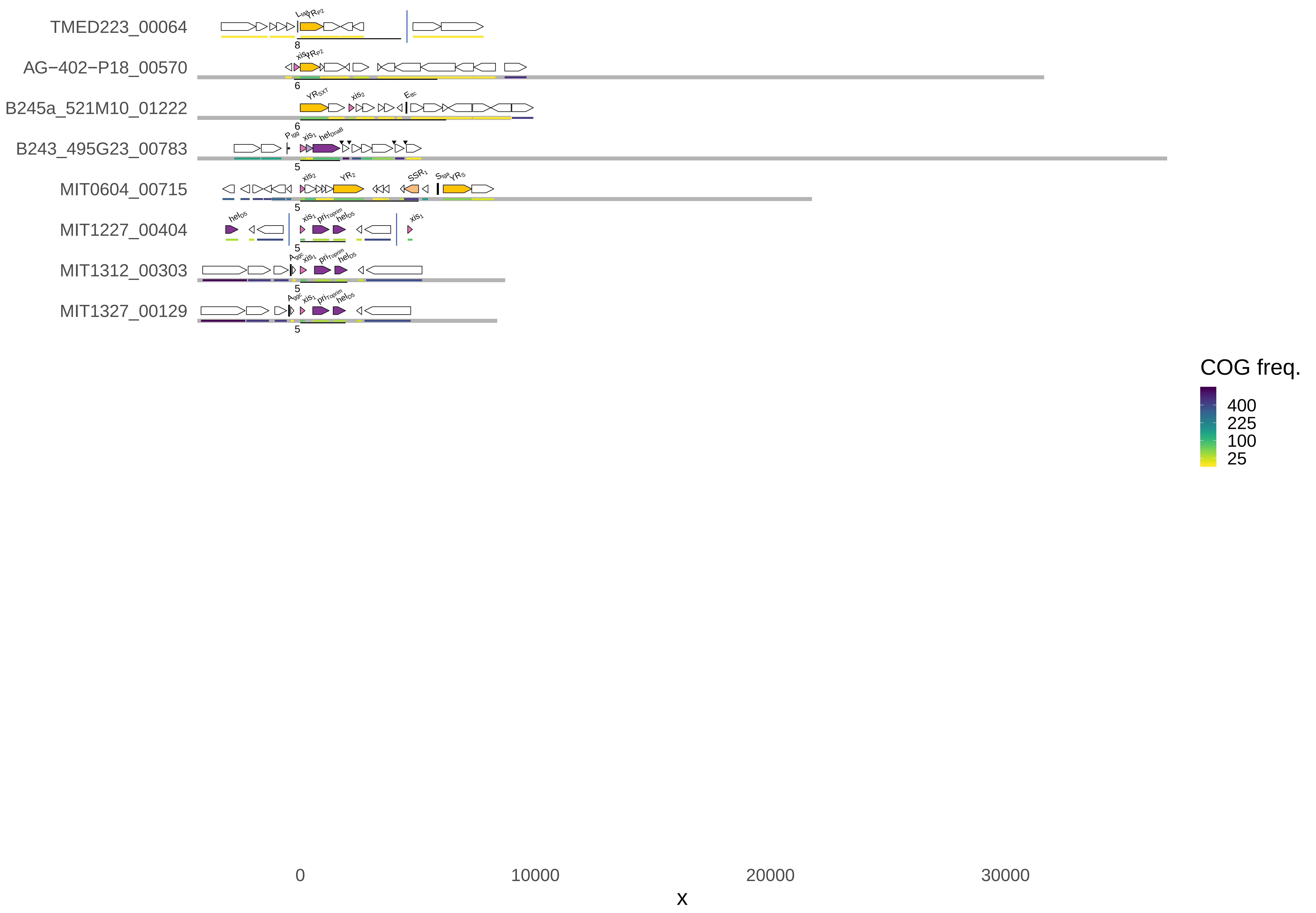
